# Relative genomic impacts of translocation history, hatchery practices, and farm selection in Pacific oyster *Crassostrea gigas* throughout the Northern Hemisphere

**DOI:** 10.1101/847467

**Authors:** Ben J. G. Sutherland, Claire Rycroft, Anne-Laure Ferchaud, Rob Saunders, Li Li, Sheng Liu, Amy M. Chan, Sarah P. Otto, Curtis A. Suttle, Kristina M. Miller

## Abstract

Pacific oyster *Crassostrea gigas*, endemic to coastal Asia, has been translocated globally throughout the past century, resulting in self-sustaining introduced populations (naturalized). Oyster aquaculture industries in many parts of the world depend on commercially available seed (hatchery-farmed) or naturalized/wild oysters to move onto a farm (naturalized-farmed). It is therefore important to understand genetic variation among populations and farm types. Here we genotype naturalized/wild populations from France, Japan, China, and most extensively in coastal British Columbia, Canada. We also genotype cultured populations from throughout the Northern Hemisphere to compare with naturalized populations. In total, 16,942 markers were identified using double-digest RAD-sequencing in 182 naturalized, 112 hatchery-farmed, and 72 naturalized-farmed oysters (n = 366). Consistent with previous studies, very low genetic differentiation was observed around Vancouver Island (mean F_ST_ = 0.0019), and low differentiation between countries in the Japan-Canada-France historical translocation lineage (France-Canada F_ST_ = 0.0024; Japan-Canada F_ST_ = 0.0060). Chinese populations were more differentiated (China-Japan F_ST_ = 0.0241). Hatchery-propagated populations had higher inter-individual relatedness suggesting family structure. Within-population inbreeding was not detected on farms, but nucleotide diversity and polymorphism rate was lower in one farm population. Moving oysters from nature onto farms did not result in strong within-generation selection. Private alleles at substantial frequency were identified in several hatchery populations grown in BC, suggesting non-local origins. Tests of selection identified outlier loci consistent with selective differences associated with domestication, in some cases consistently identified in multiple farms. Top outlier candidates were nearby genes involved in calcium signaling and calmodulin activity. Implications of potential introgression from hatchery-farmed oysters depends on whether naturalized populations are valued as a locally-adapted resource or as an introduced, invasive species. Given the value of the industry in BC and the challenges the industry faces (e.g., climate change, crop losses, biotic stressors), this remains an important question.

## INTRODUCTION

Wild populations contain standing genetic variation that is critical for adaptation to changing environments (Barrett & Schluter, 2008). When domestic strains of such species exist, genetic differences between wild and domestic lineages can provide an important source of variation for broodstock development (Guo, 2009). Identifying, preserving, and using these genetic resources effectively is essential for sustainable development of aquaculture species (Guo, 2009). Genetic markers that differentiate populations can be useful to assign unknown origin individuals to populations of origin (Beacham et al., 2017), to detect gene flow from domesticated populations (e.g., introgression of domesticated alleles) (Wringe et al., 2018), and to manage broodstock (Arbelaez et al., 2019). Once markers are identified, they can be used to generate robust, multi-purpose high throughput amplicon sequencing panels (Meek & Larson, 2019) or be added to SNP chips (Gutierrez et al., 2017). Better understanding of the genetics of cultured organisms co-existing with wild populations is needed to provide tools for monitoring the diversity of the species, to direct selective breeding efforts, and to ensure good husbandry practices.

Genetics is particularly useful to understand population connectivity in aquatic organisms due to the difficulty in directly observing organism movement (Gagnaire et al., 2015). However, marine invertebrates have high fecundity and dispersal, as well as large population sizes, and these factors often result in little to no signal of genetic differentiation (Gagnaire et al., 2015). High fecundity in particular can result in chaotic genetic patchiness, where genetic differentiation between patches does not correlate with spatial distance and can fluctuate over time (Eldon et al., 2016). This phenomenon is particularly pronounced when there is sweepstakes reproductive success, where a small number of individuals produce many offspring, due to random matching of reproductive timing and favourable environment conditions. Sweepstakes reproductive success can therefore result in a low effective population size (Ne) relative to census size (Nc) (Hedgecock & Pudovkin, 2011). Chaotic genetic patchiness has been observed in many marine invertebrates (Villacorta-Rath et al., 2017), including the Pacific oyster (Sun & Hedgecock, 2017). Nonetheless, fine-scale genetic structure has been identified in marine invertebrates such as the American lobster *Homarus americanus* (Benestan et al., 2015) and the sea scallop *Placopecten magellanicus* (Lehnert et al., 2018). The use of high-density markers combined with environmental metadata improves the likelihood of identifying such genetic structure.

High fecundity can also result in high polymorphism rate, and this has been observed in Pacific oyster (Plough, 2016). Gametogenesis in species with a large gametic output, consistent with sweepstakes reproductive success, have many mitotic cell divisions, prior to meiosis, during which mutations accumulate and elevate levels of heterozygosity (Williams, 1975; Launey & Hedgecock, 2001). Furthermore, because mutations can arise early in gametogenesis, leading to a high frequency among the gametes (“jackpot mutations”), deleterious mutations can occasionally be introduced at high frequency (Harrang et al., 2013). The enormous number of mitotic divisions leading to gamete production in species such as the Pacific oyster is expected to generate a high genetic load, with numerous deleterious recessive mutations. This can result in high rates of larval mortality (Launey & Hedgecock, 2001), and severe inbreeding effects (Hubert, 2004; Hedgecock et al., 2005). To counteract such genetic effects, breeders have been advised to cross inbred strains to benefit from heterosis among the offspring (Hedgecock & Davis, 2007). In a similar context, crossing divergent populations may provide heterozygote advantage, but only if the populations are genetically distinct. In a species with the potential for the emergence of deleterious recessive mutations, breeding programs are highly relevant and important to consider.

Bivalve aquaculture is a valuable resource globally. Most of the harvested production originates from aquaculture (89%), and the global, per annum value of farmed bivalves is 20.6 B USD (Wijsman et al., 2019). In BC in 2016, the estimated value of Pacific oyster aquaculture was 14.8 M CAD in revenue (Sun & Hallin, 2018). The oyster aquaculture industry depends on either commercial hatcheries for production, or natural recruitment events where naturally set oysters are transplanted to a farm. When commercial hatcheries are used, the exact country of origin of the broodstock is rarely known, and oysters from multiple sources are often combined together and grown on a farm. Whether naturalized (or wild) populations exhibit local adaptation has yet to be demonstrated, but such adaptation could allow populations to be more fit to local conditions in terms of food availability, disease, harmful algae, and levels of oxygen, temperature, and partial pressure of carbon dioxide (pCO2). Identifying genes involved in either domestication or local adaptation will be informative for guiding selective breeding. Identifying haplotypes specific to broodstock lines will also be useful for tracking introgression of international commercial hatchery populations into naturalized or wild populations. Best practices for broodstock development could both require maintenance of genetic diversity as well as the incorporation of alleles conferring higher survivorship under adverse conditions or higher fitness in specific environments. Although relatedness can be determined through pedigrees or low-density panels (Kijas et al., 2019), high-density genetic studies are necessary to pinpoint informative genes and markers.

Farmed Pacific oyster are often grown in areas in which they are not endemic (De Silva, 2012). Pacific oyster originated in East Asia (i.e., coastal Japan, Korea, North China) but has been translocated globally for over a century (Guo, 2009; De Silva, 2012). Pacific oyster has now been introduced to over 66 countries and grows in self-sustaining (i.e., naturalized) populations in at least 17 countries (Herbert et al., 2016). In Canada, Pacific oyster was introduced repeatedly in the early 1910s until the 1960s to supplement the declining Olympia oyster *Ostrea conchaphila* fishery (Guo, 2009). In France, Pacific oyster was introduced in the 1970s using broodstock (i.e., adult individuals) from British Columbia (BC) and spat (i.e., juveniles attached to substrate) from Japan (Grizel & Heral, 1991). Translocations have therefore resulted in many naturalized populations in coastal BC and Europe (e.g., France, Denmark) (Herbert et al., 2016; Anglès d’Auriac et al., 2017; Reise et al., 2017).

Population genetic analyses to date have not found evidence for bottlenecks during Pacific oyster introductions in BC or France (Sun & Hedgecock, 2017; Vendrami et al., 2018). Very low genetic differentiation between translocated naturalized populations and Japanese populations has been observed (Vendrami et al., 2018). Low differentiation has also been observed throughout China, with slightly elevated F_ST_ between China and Japan (Li et al., 2015). While not found in BC or France, bottlenecks have resulted in lower genetic diversity in regions that were recently colonized followed by the rapid expansion of oyster populations, such as the Wadden Sea of Denmark (Vendrami et al., 2018).

In the North East Pacific ocean (e.g. BC and Washington State, USA), low population differentiation among sites has been observed using 52 coding SNP markers (Sun & Hedgecock, 2017). This study also identified higher temporal variation than spatial variation, providing evidence for sweepstakes reproductive success (Sun & Hedgecock, 2017). It is possible, however, that with a higher density of markers, including non-coding loci, spatial population structure can be revealed in coastal BC.

Here we address three important aspects of the population genetics of Pacific oyster. First, we examine the genetic diversity and differentiation of oysters within BC by characterizing four naturalized populations, including two collections occurring across multiple brood years. Wild/naturalized oysters from Japan, France, and China were also genotyped to put the observed genetic differentiation into a broader context. All samples were genotyped using double-digest restriction-associated DNA sequencing (ddRAD-seq) of 16,942 RAD loci, representing ~0.255% of the Pacific oyster genome. Second, to better characterize differences between hatchery-farmed and naturalized oysters, several farm populations grown in BC, France, the United Kingdom (UK), and China were genotyped at these same markers. This permitted the identification of loci that may be subject to different selective pressures between hatcheries/farms and nature (i.e., subject to “domestication selection”). Third, to characterize the selective landscape associated with growth on a farm, naturalized oysters that had recently been moved to a farm were compared to their source populations in three parallel comparisons across the Northern Hemisphere (i.e., Canada, France, and China). Collectively, these analyses provide insight into the relative genomic impacts of hatchery practices, farming, and translocation history on the contemporary genetic diversity that exists in Pacific oysters throughout the Northern Hemisphere.

## METHODS

### Sample collections

Pacific oyster *Crassostrea gigas* individuals were sampled from a variety of sources, including those growing in nature, either as wild (endemic) populations or as naturalized (introduced) populations (both referred to as NAT), those growing in farms, either from commercial hatcheries (HAT-F) or from NAT populations moved from nature onto farms (NAT-F) as described by local farmers in each area, as well as those from hatcheries destined for farms but not yet transferred onto a farm (HAT). NAT samples were obtained from introduced populations (i.e., BC, France) or wild populations (i.e., China, Japan; Table 1). HAT-F samples were obtained from BC with origins from commercial hatcheries in Canada, US, and/or Chile (exact source unknown), as well as from China, with origins from wild Chinese populations.

**Table 1.** Sample overview, including collection location, sample type, short form identifier, and sample size and approximate length for retained, genotyped samples. Collection locations are shown in Figures 1 and 2. Origin of seed was in some cases difficult to determine, and these collections.

| Collection Location | Type | Origin of Seed | Identifier | Sample Size | Approx. Size (cm) |
| --- | --- | --- | --- | --- | --- |
| Hisnit Inlet, BC | NAT | BC | HIS | 39 | 5.0, 13.5 |
| Pipestem Inlet, BC | NAT | BC | PIP | 25 | 1.0-6.0, 19.0 |
| Serpentine River, BC | NAT | BC | SER | 23 | 4.0-7.7 |
| Pendrell Sound, BC | NAT | BC | PEN | 35 | 3.0-4.0, 5.0-6.0 |
| Baynes Sound*, BC | NAT-F | BC | PEN-F | 24 | 2.0-4.5 |
| Rosewall Creek, BC | HAT-F | US and/or BC | ROS | 23 | 10.0-14.0 |
| Deep Bay, BC | HAT-F | US, BC, and/or Chile | DPB | 20 | 4.5-6.5 |
| Guernsey, UK | HAT | BC | GUR | 22 | 8.0-11.0 |
| N. Brittany, France | NAT | France | FRA | 23 | 6.5-12.0 |
| SW. France | NAT-F |  | FRA-F | 24 | 9.0-11.0 |
| Hyogo and Miyagi Prefectures, Japan | *na* | Japan | JPN | 13 | *na* |
| Qingdao, China | NAT | China | CHN | 24 | 6.0-7.0 |
| Qingdao, China | NAT-F | China | CHN-F | 24 | 3.0-5.0 |
| Qingdao, China | HAT-F | China | QDC | 24 | 6.0-7.0 |
| Qingdao, China | HAT-F | China | RSC | 23 | 8.0-10.0 |

NAT = naturalized (or wild); NAT-F = naturalized moved to a farm; HAT-F = hatchery grown on a farm; HAT = hatchery not yet transplanted to farm; \* = transplant from Pendrell Sound; *na* = not available

All collections for this study were conducted in 2017 and used adult oysters. BC NAT collections were conducted from sites around Vancouver Island during summer (Figure 1A; Table 1). For all collections where possible, each oyster was measured for length and weight (see Table 1), and a section of the mantle was sampled in 95% ethanol. Naturalized populations were obtained from Pendrell Sound (PEN), Serpentine River (SER), Pipestem Inlet (PIP), and Hisnit Inlet (HIS; Figure 1). Different sizes of oysters for Pendrell Sound and Hisnit Inlet were collected and assumed to be of different brood years, although this should be considered a rough approximation as cohorts cannot be resolved perfectly on the basis of size (Hedgecock & Pudovkin, 2011). Size classes included small = 3.0-4.9 cm (size class = 2); medium = 5.1-7.0 cm (size class = 3); and large = 11.1-15.0 cm (size class = 6) oysters. Small and medium size classes were obtained for PEN (n = 15 and 20, respectively), and medium and large were collected for HIS (n = 19 and 20, respectively). BC HAT-F populations were sampled from farms located at Deep Bay (DPB) and Rosewall Creek (ROS), BC. The location on the farm that the DPB oysters were obtained is unknown (samples were provided directly by the farm), but those from ROS were obtained by sampling throughout the farm site on the beach. BC NAT-F samples were sampled from a population grown in mesh bags on a farm in Baynes Sound from several locations throughout the farm, with the naturalized source population being Pendrell Sound, BC.

**Figure 1.**
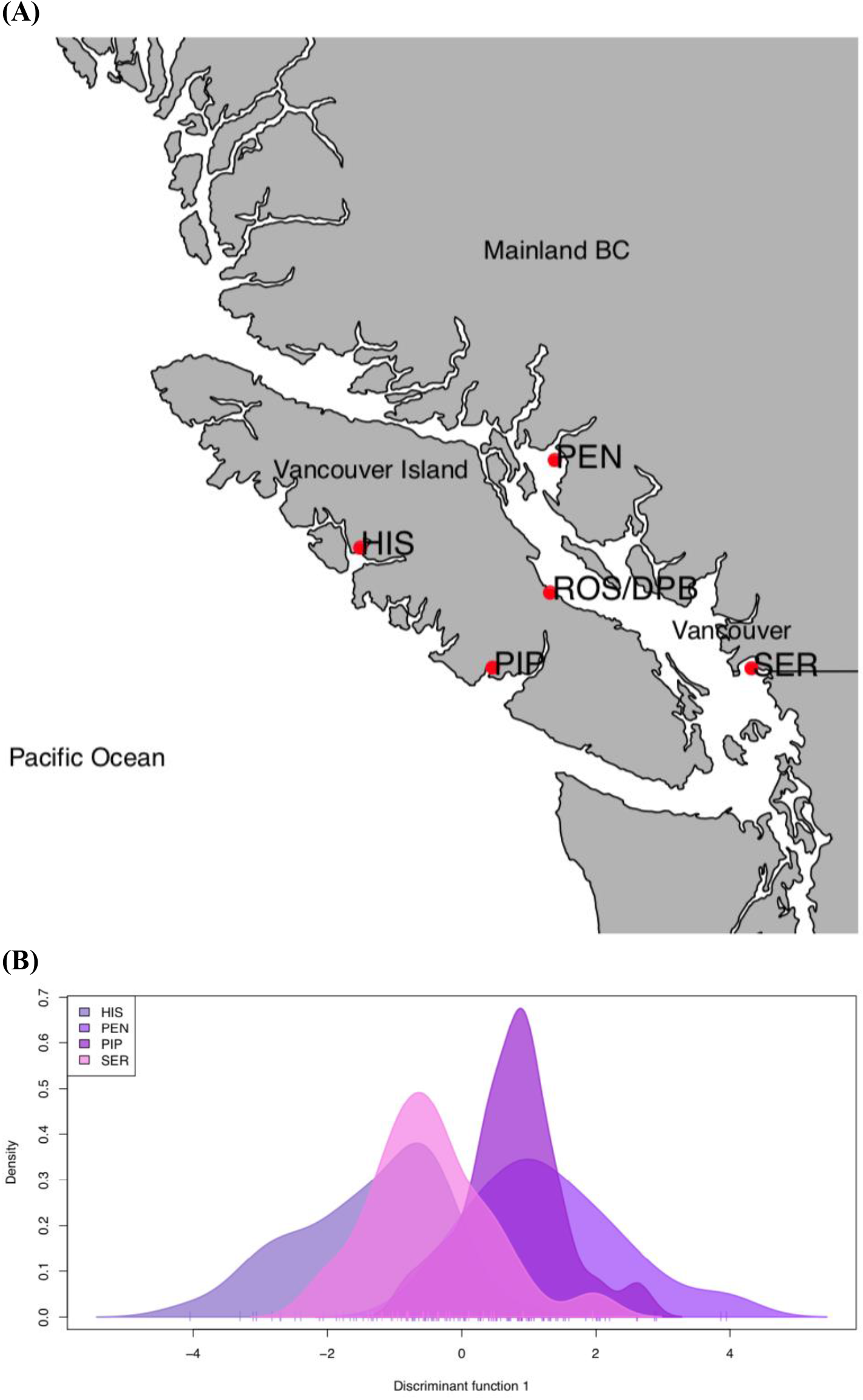
**(A)** Pacific oysters were collected from four locations in southwestern British Columbia. Pendrell Sound (PEN) and Serpentine River (SER) are on the mainland, while Pipestem Inlet (PIP), and Hisnit Inlet (HIS) are on Vancouver Island. **(B)** As shown by discriminant analysis of principal components (DAPC), there was very low to negligible genetic differentiation among these sites. Further, the naturalized populations PEN and PIP are both frequently used by industry for broodstock and are undifferentiated from each other. Populations of commercial-hatchery farms are grown at sites near Rosewall Creek (ROS) and Deep Bay (DPB), shown in (A).

International NAT samples included those from France and China. France (FRA) samples were collected as tissue samples of naturally set adult oysters in Northwest France from a beach near Sainte-Barbe (48.765167, - 2.994716). China (CHN) NAT samples were collected in the Qingdao region. NAT-F collections were also made from a farm in western France and a farm in China. China HAT-F samples were also collected, one from a short-term culture (QDC, 2^nd^ generation) and one from a long-term culture (RSC, 6^th^ generation).

HAT samples were obtained from a hatchery in Guernsey, UK (Guernsey Sea Farms), which maintains their own hatchery broodstock originally derived from BC stock (Mark Dravers and JP Hastey, *pers. comm*). Adult samples of unknown origin (farmed or wild) were also obtained from local markets in Japan. All international samples were collected by local researchers (see *Acknowledgements)* or obtained from commercial markets, obtained in accordance with local regulations, and shipped according to international regulations.

### DNA extraction, quantification and quality control

DNA was extracted using the BioSprint automated extraction protocol in plate format (QIAGEN), using a proteinase-K digestion and an RNase incubation to reduce RNA presence in genomic DNA, as per manufacturers’ instructions. Sections of tissue used for extraction were ~4 x 4 mm. Purified genomic DNA was quantified with a Qubit (High Sens; ThermoFisher) and quality checked for fragmentation using 1% agarose gel electrophoresis. Genomic DNA was normalized to 20 ng/μl for library preparation.

### Restriction site associated DNA (RAD) sequencing enzyme selection

To choose restriction enzymes and fragment sizes to produce an optimal number of loci, an *in silico* digest was simulated using the Pacific oyster reference genomes (i.e., GenBank GCA_000297895.1 (Zhang et al., 2012) and the chromosome level assembly (Gagnaire et al., 2018)) in SimRAD (Lepais & Weir, 2014). Initial digests with PstI and *MspI* indicated low cut frequency due to the low GC content in the oyster genome (33.5%); therefore, *NsiI*, a 6-base cutter with fewer GC recognition sites was used in place of PstI. Using NsiI and *MspI* with a size selection of 100-250 bp, 15,770 and 15,319 sites were predicted using the Zhang *et al*. (2012) and Gagnaire *et al*. (2018) genomes, assuming that the assembly was 93% and 47% complete, respectively (see code in Data Accessibility). The number of markers was calculated based on the 100% assembly expansion (i.e., a complete genome size of 600 Mb).

### Double-digest RAD-sequencing library preparation

In a 96 well plate, 10 μl of 20 ng/ul DNA per individual was fragmented using the high-fidelity enzymes *NsiI* / *MspI* (NEB) combined in a double digest, then ligated via sticky ends to a unique forward adapter and a common Y-adapter specific to the Ion Torrent Proton (Thermo Fisher) following previously published procedures (Mascher et al., 2013), using barcodes and size-selection adaptation as described previously (Recknagel et al., 2015). The entire ddRAD-seq protocol is provided on protocols.io (see *Data Availability*). In brief, genomic DNA was digested and ligated to barcodes, then 5 μl of each sample was multiplexed into libraries of between 24 and 48 samples, purified using PCR clean-up columns (QIAGEN), size-selected for a size range of 250-300 bp on a Pippin Prep (Sage Biosciences). After size-selection, four PCR reactions (16 cycles) were conducted and pooled to reduce PCR bias, similar to Mascher *et al*. (2013). Each PCR was pooled and then cleaned up with PCR clean-up columns (QIAGEN). Pools were quantified by Qubit, and diluted to 200 pM in preparation for sequencing on an Ion Torrent Proton.

Libraries were sequenced on an Ion Torrent Proton using an Ion PI Chip Kit v3 chip (Thermo Fisher), as per manufacturers’ instructions. Base calling was done with Torrent Suite 5.10.1 (Thermo Fisher) and multiplexed FASTQ output files with barcodes still present were exported using the Torrent *FileExporter* plugin.

### Sequence data pre-processing and alignment against reference genome for genotyping

All sequence processing and analysis occurred within a modified version of *stacks_workflow* (E. Normandeau,*pers. comm*.; see *Data Availability)* with custom analysis scripts and documentation within an additional, separate GitHub repository (see *Data Availability*). Raw FASTQ files were inspected before and after trimming using FastQC v0.11.8 (Andrews, 2010), with output aggregated by MultiQC (Ewels et al., 2016). Universal adapters were removed, and all reads greater than 50 bp were retained using cutadapt v1.18 (Martin, 2011) in parallel (Tange, 2011). All reads were quality trimmed (Phred > 10 in sliding windows of size 15% of the read length), truncated to 80 bp, and demultiplexed with the two enzyme setting (i.e., NsiI and MspI) using *process_radtags* of *Stacks* v2.3e (Catchen et al., 2011; Rochette et al., 2019) in parallel. Custom scripts were then used to combine the same individuals from different sequencing runs, and to prepare the population map for *Stacks* (see *Data Availability*).

Individual read files were aligned in parallel to the most complete Pacific oyster genome (Zhang et al., 2012) using *bwa mem* v0.7.12-r1039 (Li et al., 2009). Insertion and deletion sequencing errors are known to occur in Ion Proton systems, and therefore non-default flags were used in the alignment, specifically -O 0,0 (no gap open penalties); -E 2,2 (reduced gap extension penalty); -T 0 (no minimum score to produce output) (E. Normandeau, *pers. comm*.). Subsequently, *samtools view* v1.9 (Li et al., 2009) was used to remove unmapped reads (-F 4), non-primary alignments (-F 256 -F 2048), as well as low quality alignments (-q 1), and convert the sam to bam format. Samples with fewer than 1 M reads were removed from the analysis using automated custom scripts (see *Data Availability*), and all remaining samples were used for genotyping with *Stacks* v2.3e (Rochette et al., 2019). A new population map with only retained samples was generated, and the total number of reads per sample, total read alignments, and mapping percentage were calculated using *samtools* and custom scripts (see *Data Availability*).

### Genotyping and filtering

Loci were identified with *Stacks* v.2.3e using the reference-based approach with alignments against the most complete Pacific oyster reference genome, estimated at 93% complete (Zhang et al., 2012) with the *Stacks* module *gstacks* with program defaults. The *populations* module and custom scripts *(Data Availability*) were used to identify loci out of Hardy-Weinberg equilibrium in at least three populations (p < 0.01 in each population) to make a list of markers to exclude (i.e., a blacklist). The *populations* module using the blacklist was then used to output multi-locus genotype files with either a single SNP per RAD-tag or as microhaplotypes (i.e., all SNPs genotyped and passing filters in a RAD-tag are grouped in a microhaplotype). The single SNP output from *populations* used the following filtering parameters: -r 0.7 (i.e., minimum proportion of individuals genotyped per population); --min_populations 15 (i.e., genotyped in all populations in the population map); --min-maf 0.01 (i.e., global minor allele frequency filter); --write-single-snp (i.e., restricts data to the first SNP per RAD locus); a blacklist including all loci identified as out of HWE; --fasta-loci (i.e., output one locus per marker in fasta); --vcf (i.e., VCF output) and --plink (i.e., plink format output). For the microhaplotype data output from *Stacks* as a VCF, the parameters were the same as above except allowing multiple SNPs per marker and including the --radpainter flag to output into fineRADstructure input (i.e., SimpleMatrix format) (Malinsky et al., 2016).

### Population Differentiation

The single SNP per RAD-tag data was translated to plink format (Purcell *et al*. 2007) by *Stacks*, which was then used to recode the VCF (--recodeA) into the number of alleles per locus, per individual. The recoded VCF was input to *adegenet* (Jombart, 2008) in R (R Core Team 2018) for multivariate analyses (i.e., Principal Components Analysis (PCA) and Discriminant Analysis of Principal Components (dAPC)). The genotypes were input to *hierfstat* (Goudet, 2005) to calculate pairwise F_ST_ (Weir & Cockerham, 1984) using 1,000 bootstraps across loci. When the lower limit of the 95% confidence interval was negative, the lower limit was transformed to 0 and there was considered to be no differentiation between the collections.

### Genetic Diversity and Relatedness

Per population observed and expected heterozygosity and homozygosity, as well as inbreeding coefficient (F_IS_), were calculated using a single marker per locus (i.e., first variant of the locus) considering variant positions only, in the *populations* module of *Stacks*. Per population, per marker estimates of nucleotide diversity (π) were calculated using VCFtools (--site-pi) (Danecek et al., 2011) using the multi-locus genotypes containing a single SNP per marker, and analyzed in R (see *Data Availability*).

Relatedness based on shared polymorphisms was calculated using a single SNP per marker using *related* (Pew et al., 2015) by exporting data from *Stacks* into R via *plink* and *adegenet*, then converted to *demerelate* (Kraemer & Gerlach, 2017) to format to *related* input. Shared polymorphisms were calculated using *ritland* (Ritland, 1996), *wang* (Wang, 2002) and *quellergt* (Queller & Goodnight, 1989) statistics for coancestry analysis and inter-individual relatedness (only results using *ritland* are shown, as all showed similar trends). Relatedness was also calculated based on shared microhaplotypes using fineRADstructure (Malinsky et al., 2016). In brief, the coancestry matrix was calculated using *RADpainter*, then individuals were assigned to populations and a tree built using *finestructure*, then output was plotted using scripts provided with *fineRADstructure* (Malinsky et al., 2016).

After inspecting genetic similarity among collections, a single population per general region or broodstock line was selected and private alleles identified using *poppr* (Kamvar et al., 2014). To identify alleles specific to farms, all BC naturalized (including farm transplants) were combined together and compared against the hatchery-farmed populations.

### Detection of domestication outliers

Domestication outliers were considered as those with evidence of differential selection between HAT-F/HAT with NAT populations. Outliers were identified by comparing HAT-F/HAT populations with a NAT population that was most similar to the expected source of the broodstock, when possible (see top of Table 2). Four different approaches were applied to identify outliers. Candidate loci were first identified using *pcadapt* v4.1.0 (Luu et al., 2017). Individual comparisons were made for each HAT-F/HAT and NAT contrast (Table 2). The principal components (PCs) retained in the analysis (*k* in pcadapt) were retained based on the flattening of the slope on scree plots to keep only those to the left of the plateau as suggested, retaining *k* = 3, 5, 6, 6, 3 for DPB, ROS, GUR, QDC, and RSC, respectively (Additional File S1). Further, score plots within *pcadapt* were used to identify which PCs separated the samples based on the HAT-F/HAT vs. NAT comparison (Additional File S2). Outliers were retained that were significant, as determined by *get.pc* of *pcadapt*, along the following PCs: DPB =PC1,3; ROS = PC1,3; 1,2,3,4; GUR = PC1; QDC = 1,2,3,4; and RSC = PC1. Outlier p-values were generated, and false discovery rates controlled using *qvalue* (Storey & Tibshirani, 2003; Storey et al., 2019). Markers with q < 0.01 for the PCs described above were considered significant. Second, *BayeScan* v2.1 with a prior odds of 100 (Foll & Gaggiotti, 2008) was also used to identify outliers in these contrasts. Third, a Redundancy Analysis (RDA) was conducted using the HAT-F/HAT and NAT collections in Table 2 combined by source type (HAT-F/HAT vs. NAT) as a multi-locus genotype-environment association (GEA) using *vegan* v2.5-5 (Oksanen et al., 2019). RDA, an analog of multivariate linear regression analysis, used the multi-locus genotype data per individual as a dependent variable and grouping (HAT-F or NAT) as the explanatory variable. The function *rda* was used to compute RDA on the model separating samples based on the grouping factor (e.g., Laporte et al., 2016). An analysis of variance (ANOVA; 1,000 permutations) was performed to determine the significance of the RDAs, and the percentage of variance explained (PVE) was calculated using *RsquareAdj* in the *vegan* package. The identification of outliers was conducted using instructions as per previous work and online tutorials (Forester et al., 2018; Forester, 2019), considering outliers as the markers with loading values greater than 3.5 standard deviations of the mean. The fourth approach identified markers in the top 99^th^ percentile of the F_ST_ distribution and considered these as potential outliers.

**Table 2.** Outlier overview, including values for both the domestication and on-farm selection comparisons. Outliers were identified per grouping by RDA (single value per grouping comparison; > 3.5x SD of the mean), and per contrast by *pcadapt*. (q < 0.01). Outliers in the 99^th^ percentile of the F_ST_ per contrast were also identified. F_ST_ lower limits that overlapped with 0 resulted in the entire interval and average being transformed to 0. The number of outliers per contrast that were detected by at least two methods were enumerated per contrast (‘Common outliers’). Note: this includes all outliers, including those that do not align to the chromosome-level assembly.

| Type | Comparison | $F_{ST}$ (avg.) | $F_{ST}$ (95% CI) | Outliers (*pcadapt*) | Outliers ($F_{ST}$) | Outliers (RDA) | Common outliers |
| --- | --- | --- | --- | --- | --- | --- | --- |
| **Domestication**<br>(HAT/HAT-F vs. NAT) |  |  |  |  |  | 170 |  |
|  | DPB-PEN | 0.0604 | 0.0574-0.0636 | 203 | 137 |  | 53 |
|  | ROS-PIP | 0.0080 | 0.0069-0.0090 | 309 | 135 |  | 64 |
|  | GUR-FRA | 0.0281 | 0.0270-0.0299 | 82 | 132 |  | 50 |
|  | QDC-CHN | 0.0196 | 0.0181-0.0212 | 367 | 129 |  | 61 |
|  | RSC-CHN | 0.0087 | 0.0074-0.0100 | 83 | 134 |  | 31 |
| **On-farm selection**<br>(NAT-F vs. NAT) |  |  |  |  |  | 175 |  |
|  | CHN-CHNF | 0.0003 | 0-0.0012 | NA | 134 |  | 18 |
|  | FRA-FRAF | 0.0004 | 0-0.0014 | NA | 140 |  | 13 |
|  | PEN-PENF | 0.0008 | 0-0.0015 | NA | 144 |  | 26 |

Outliers significant using both RDA and *pcadapt*, or outliers that showed significance using any two statistical tests as well as being observed in multiple comparisons, were considered as potential loci subjected to differential selection between naturalized and farm populations and were flagged for further investigation. These top outliers were used to search against the reference genome to determine whether the they were within or near predicted genes. The annotation from NCBI, NCBI_annot_rel_101_2017-02-02 from the Zhang et al. (2012) genome was principally used, and genes containing the outlier (e.g., in introns or exons), or those within approximately 10 kb of the outlier position were recorded. The 10 kb distance was used given the rapid decay of linkage disequilibrium observed in the Pacific oyster (Kijas et al., 2019).

### Detection of outliers under selection from growing on farm

To identify markers that changed significantly in allele frequency following transplantation from naturalized populations onto farms, a pairwise comparison was conducted between NAT and NAT-F oysters for populations in BC, China and France. The comparison was between a nearby, naturalized population and the transplanted farm population (i.e., FRA vs. FRA-F; CHN vs. CHN-F; PEN vs. PEN-F; see bottom of Table 2). Outliers were identified using the same approach as above (i.e.,*pcadapt, BayeScan*, RDA, 99^th^ percentile F_ST_). However, scree plots did not indicate a plateau (Additional File S1), suggesting that no PCs explained a majority of the variation, and so *pcadapt* was not used for these comparisons. To determine whether similar signatures of selection were found between different NAT vs NAT-F contrasts, suggesting parallel selection, per locus F_ST_ was directly compared between each pair of countries (i.e., BC, FRA, CHN).

### Alignment of outliers to the chromosome-level assembly

All markers identified using the most complete reference genome (Zhang et al., 2012) that were exported from *Stacks* in *fasta* format were collectively aligned to the chromosome-level assembly (Gagnaire et al., 2018) using *bwa mem* and *samtools*. Only those that aligned to the chromosome-level assembly were used for plotting along chromosomes but for all other purposes both aligned and unaligned markers against the chromosome-level assembly were included. Per locus F_ST_ values calculated on the pairs of populations used in the *pcadapt* contrasts (Table 2) were plotted along chromosomes per contrast, with outliers identified by each tool indicated in the plot using custom scripts (see *Data Availability*).

## RESULTS

### Sample sequencing and genotyping

In total, 182 naturalized/wild individuals (NAT) were genotyped from coasts of Vancouver Island, British Columbia (BC; n = 122), France (FRA n = 23), China (CHN; n = 24), and Japan (JPN; n = 13; Table 1; Figure 1A). For naturalized oysters transplanted to a farm (NAT-F), 72 oysters were genotyped, with parallel comparisons in Canada, France, and China (n = 24 for each), to compare to expected source populations. Finally, 112 hatchery-origin oysters grown on a farm (HAT-F or HAT) were genotyped (Table 1).

On average, samples had a total of 2.17 M reads (min = 1.01 M; median = 2.06 M; max = 6.17 M). Aligning to the most complete reference genome (NCBI: GCA_000297895.1; oyster_v9; Zhang et al. 2012) resulted in an average alignment rate of 69.7%. Genotyping with *gstacks* resulted in a total of 723,034 unfiltered loci, with an average effective per-sample coverage of 20.2x (sd = 5.7x, min = 10.3x, max = 48.6x). Loci were then filtered for missing data by only retaining markers that were genotyped in at least 70% of the individuals per population in all populations, that had a minor allele frequency (global MAF > 0.01), and that were not out of HWE equilibrium in three or more populations (p < 0.01 in each). This resulted in a total of 81,443 SNPs found in 17,084 loci when considering microhaplotypes. When filters were applied to only the first SNP per RAD locus (see above), a total of 16,942 markers were retained. These 16,942 markers comprised 1,532,384 bp of total sequence, representing 0.255% of the expected 600 Mbp Pacific oyster genome.

Filtered loci were subsequently aligned to the chromosome-level reference genome (Gagnaire et al., 2018) for the purpose of plotting outlier statistics (see below), indicating that loci occurred throughout the genome (Table 3). A total of 9,286 of the filtered RAD-loci mapped to the chromosome-level assembly (54.8% of filtered loci), which fits with the expectation of the chromosome-level genome being approximately 50% present in the assembly (Gagnaire et al., 2018). All markers, including those not aligned to the chromosome assembly, were retained for downstream analyses, with the exception of plots by chromosome.

**Table 3.** Markers aligning to the chromosome-level assembly or only to the contig-level assembly are shown as either as numbers or percentages of aligned markers per chromosome. All RAD-loci (n = 16,942) were used for all analysis except for plotting purposes along chromosomes.

| Chromosome | Number of markers | Percentage of total aligned |
| --- | --- | --- |
| A | 1,058 | 11.4% |
| B | 732 | 7.9% |
| C | 882 | 9.5% |
| D | 1,471 | 15.8% |
| E | 1,234 | 13.3% |
| F | 978 | 10.5% |
| G | 1,027 | 11.1% |
| H | 581 | 6.3% |
| I | 762 | 8.2% |
| J | 561 | 6.0% |
| **Total aligned** | 9,286 | - |
| Percent aligned | 54.8% | - |
| Not aligned | 7,656 | - |
| **Total markers** | 16,942 | - |

### Genetic differentiation among naturalized locations in British Columbia

Naturalized populations from around Vancouver Island, BC (Figure 1A) ranged from no to very low differentiation from one another (Table 4A). On the mainland, very low differentiation was observed between Pendrell Sound (PEN) and Serpentine River (SER) (F_ST_ 95% CI = 0.0017-0.0033; Table 4A; Table S1A for average F_ST_). Moreover, no differentiation was observed (i.e., F_ST_ = 0 could not be rejected) between PEN and Pipestem Inlet (PIP), even though they are on opposite sides of Vancouver Island. Discriminant Analysis of Principle Components (DAPC) also demonstrated highly overlapping genotypic distributions among these locations (Figure 1B).

**Table 4.** Genetic differentiation (F_ST_) between **(A)** naturalized oyster populations collected from around Vancouver Island, BC; **(B)** between size classes representing different brood years and locations for naturalized populations around Vancouver Island, BC; **(C)** all pairs of populations in the study, using a single representative for each set of similar populations (e.g. BC naturalized represented by PEN). Statistics are shown as 95% confidence intervals, with lower estimates below the diagonal and upper estimates above the diagonal. Any negative lower limits were transformed to 0, and that contrast is assumed to be not differentiated. Full population names and locations are shown in Table 1 and Figures 1 and 2, respectively.

**(A)**
|  | HIS | PEN | PIP | SER |
| --- | --- | --- | --- | --- |
| HIS | - | 0.0035 | 0.0031 | 0.0025 |
| PEN | 0.0022 | - | 0.0012 | 0.0033 |
| PIP | 0.0014 | 0 | - | 0.0025 |
| SER | 0.0009 | 0.0017 | 0.0007 | - |

**(B)**
|  | PEN_2 | PEN_3 | HIS_3 | HIS_6 |
| --- | --- | --- | --- | --- |
| PEN_2 | - | 0.0011 | 0.0046 | 0.0043 |
| PEN_3 | 0 | - | 0.0048 | 0.0043 |
| HIS_3 | 0.0018 | 0.0024 | - | 0.0027 |
| HIS_6 | 0.0017 | 0.0019 | 0.0004 | - |

(C)
|  | PEN | CHN | QDC | RSC | DPB | ROS | GUR | FRA | JPN |
| --- | --- | --- | --- | --- | --- | --- | --- | --- | --- |
| **PEN** | - | 0.0229 | 0.0424 | 0.0147 | 0.0636 | 0.0096 | 0.0307 | 0.0032 | 0.0074 |
| **CHN** | 0.0179 | - | 0.0212 | 0.0100 | 0.0839 | 0.0308 | 0.0488 | 0.0218 | 0.0272 |
| **QDC** | 0.0371 | 0.0181 | - | 0.0285 | 0.1007 | 0.0505 | 0.0671 | 0.0403 | 0.0450 |
| **RSC** | 0.0119 | 0.0074 | 0.0248 | - | 0.0756 | 0.0223 | 0.0405 | 0.0135 | 0.0185 |
| **DPB** | 0.0574 | 0.0756 | 0.0920 | 0.0689 | - | 0.0722 | 0.0760 | 0.0640 | 0.0668 |
| **ROS** | 0.0075 | 0.0257 | 0.0445 | 0.0188 | 0.0660 | - | 0.0372 | 0.0095 | 0.0145 |
| **GUR** | 0.0270 | 0.0431 | 0.0600 | 0.0360 | 0.0697 | 0.0332 | - | 0.0299 | 0.0336 |
| **FRA** | 0.0016 | 0.0168 | 0.0346 | 0.0105 | 0.0582 | 0.0071 | 0.0261 | - | 0.0063 |
| **JPN** | 0.0047 | 0.0211 | 0.0383 | 0.0148 | 0.0604 | 0.0114 | 0.0291 | 0.0034 | - |

To evaluate the relative impacts of temporal versus spatial effects on population signatures in these BC collections, multiple size classes that were assumed to be different brood years (but see Methods and Discussion for criticisms to this approach) were obtained for HIS and PEN (Table 4). Small (size class = 2), medium (size class = 3), and large (size class = 6) oysters were used in the comparisons (see Methods). Small and medium size classes were obtained for PEN (n = 15 and 20, respectively), and medium and large were collected for HIS (n = 19 and 20, respectively). The difference between sizes was therefore greater for HIS than for PEN. There was no differentiation between PEN_2 and PEN_3 (F_ST_ not significantly different from 0 (F_ST_ 95% CI: 0-0.0011), and very low differentiation between HIS_3 and HIS_6 (F_ST_ 95% CI: 0.0004-0.0027; Table 4B or Table S1B for average F_ST_). Size-matched spatial differences between PEN_3 and HIS_3 (F_ST_ 95% CI: 0.0024-0.0048) were greater than temporal differences within a site for PEN but not significantly different for HIS

### Genetic differentiation in naturalized/wild oysters throughout the Northern Hemisphere

The maximum differentiation among naturalized sites in BC was F_ST_ = 0.0022-0.0035 (95% CI) between PEN and HIS. This level of genome-wide differentiation was similar to that between naturalized oysters in BC and France (F_ST_ = 0.0016-0.0032; Table 4C or Table S1C for average F_ST_; Figure 2A). Japan and BC had slightly higher, but still low differentiation (F_ST_ = 0.0047-0.0074; DAPC in Figure 2B).

**Figure 2.**
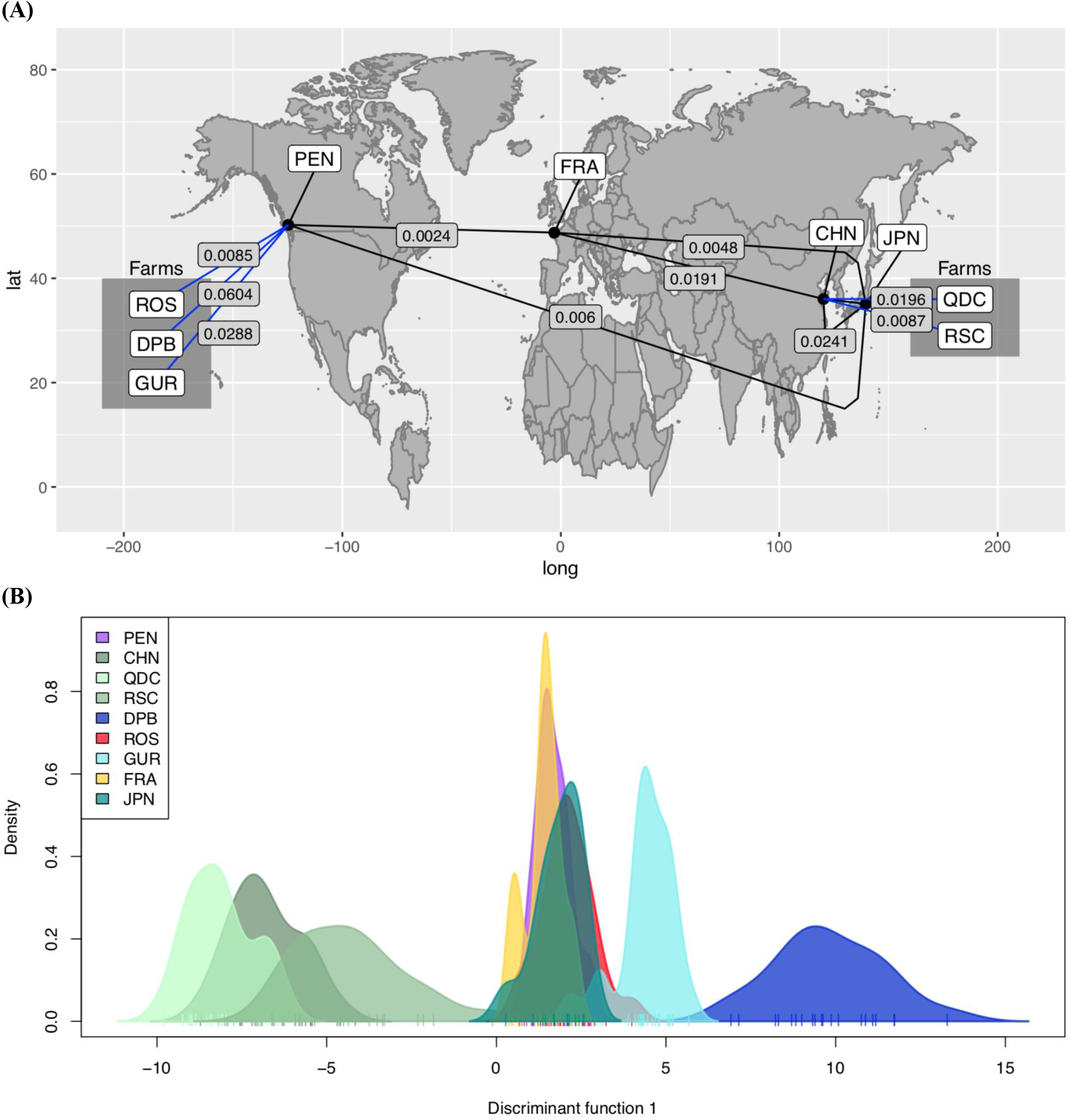
**(A)** Genetic differentiation among the self-recruiting oysters from Pendrell Sound (PEN), France (FRA), China (CHN), and Japan (JPN) compared to oysters sampled from farms thought to be growing strains descended from the same lineage (ROS, DPB, GUR). China is compared to the farms growing oysters thought to originate from China (QDC, RSC). **(B)** Discriminant analysis of Principal Components (DAPC) using all populations in the analysis indicates the similarity among BC populations, Japan, and France (peaks between 0-4 along discriminant function), among the Chinese populations (peaks between −4 to −10), and the distinct GUR (peak at 5) and DPB populations (peak at 9).

In contrast, wild Pacific oysters from China were more divergent from the JPN-CDN-FRA lineage (F_ST_ 95% CI: PEN-CHN = 0.0179-0.0229; FRA-CHN=0.0168-0.0218; JPN-CHN = 0.0211-0.0272; Table 4C). The CHN population was found to be more similar to the CHN hatchery populations (i.e., RSC, QDC; DAPC in Figure 2B). For comparison, F_ST_ between BC and China is ~8-fold higher than the most divergent of the BC populations.

### Divergence between naturalized and hatchery-farmed oysters

The most genetically distinct oyster populations in this study were consistently hatchery propagated (HAT/HAT-F). In general, to understand this divergence, HAT/HAT-F oysters were compared to their closest wild/naturalized (NAT) populations in terms of physical distance (Table 2).

In BC, the least divergent HAT-F population was ROS (PEN-ROS F_ST_ = 0.0075-0.0096), with a level of differentiation slightly higher than that observed between PEN and JPN (Figure 2B). GUR was more distinct (GUR-FRA F_ST_ = 0.0261-0.0299) and formed a single grouping along the first axis of the DAPC (Figure 2B). The most distinct population in the BC farms was DPB (DPB-PEN F_ST_ = 0.0574-0.0636; see DAPC in Figure 2B).

To identify private alleles by group, similar populations were grouped together to be analyzed, producing collections BC, DPB, ROS, GUR, RSC, QDC, CHN, FRA, JPN (Table 5A). FRA and JPN were included, but private alleles were not expected given the lack of differentiation observed between BC and these collections and the much higher sample size in the BC grouping. Using these groupings, all identified private alleles had more than five observations in the data. In the BC meta-population, 38 private alleles were found, with seven at high frequency (i.e., considered when there were more than 10 observations). Even with lower sample sizes, HAT/HAT-F populations had private alleles, with DPB having the most in the comparison (n = 49; 23 alleles at high frequency), followed by QDC (n = 17; four alleles at high frequency), and GUR (n = 11; two alleles at high frequency). Other HAT-F populations, such as ROS and RSC did not have a substantial number of private alleles (< 2).

**Table 5.** Private allele counts of the 16,942 markers identified using either (A) a single representative from each of the general regions in the dataset, with all other populations in that region included into the same group; and (B) a single population from each of the global regions. Population full names, and locations are given in Table 1 and Figures 1 and 2, respectively

|  | BC | DPB | ROS | GUR | RSC | QDC | CHN | FRA | JPN |
| --- | --- | --- | --- | --- | --- | --- | --- | --- | --- |
| **Private alleles** | 38 | 49 | 0 | 11 | 1 | 17 | 1 | 0 | 0 |
| **Sample size** | 146 | 20 | 23 | 22 | 23 | 24 | 48 | 47 | 13 |

(B)
|  | PEN | CHN | DPB | GUR |
| --- | --- | --- | --- | --- |
| **Private Alleles** | 35 | 24 | 20 | 22 |
| **Sample Size** | 35 | 24 | 20 | 22 |

Farmed oysters in China were distinct from those in the France/UK and Canada (see DAPC in Figure 2B). However, the farms were not highly different from the presumed wild population in China (F_ST_ 95% CI: CHN-RSC = 0.0074-0.0100; CHN-QDC = 0.0181-0.0212; Table 2). For example, RSC and CHN had a similar level of differentiation as ROS from PEN in Canada, and QDC and CHN was slightly less divergent than the GUR-FRA comparison in Europe (Table 2). The most distinct farmed oysters from nearby naturalized oysters was therefore the DPB population (Figure 2B; Table 5A).

In agreement with F_ST_ and DAPC analyses, a Principal Components Analysis (PCA) indicates that the majority of the variation in the genotypic data (single SNP per locus, all RAD-loci) is explained by PC1-3, although the percent variation explained (PVE) was low (Figure S1). Most samples clustered tightly in the center, and DPB and CHN separated on either side of PC1 (Figure S1A), and GUR separated on PC3 (Figure S1B). Many loci contribute to the differentiation observed in the PCA and DAPC, as observed by marker loading values for the DAPC, although a small number of markers contributed most to this difference (specifically, 16 markers exhibited loading values > 0.002; Figure S2).

### Genetic diversity, heterozygosity and relatedness

Genetic diversity as measured by nucleotide diversity (π) was lowest in the DPB HAT-F population, whereas all other collections had similar median π (Figure S3). JPN was not retained in this analysis because of its lower sample size (n = 13) and the effect of sample size on π. Lower π in DPB was not associated with elevated within population inbreeding (F_IS_); in fact, DPB had the lowest F_IS_ observed in the study, although its 95% confidence interval overlapped with that of all other populations except France (Table S2A). Several of the HAT/HAT-F collections had the lowest percentage of polymorphic sites, with DPB and GUR having 0.94% and 1.13%, respectively, compared to an average of 1.43% for the other populations (+/- 0.10%) (Table S2).

Comparing different loci, per locus nucleotide diversity was highly correlated across populations, with some loci showing consistently high or low π in multiple populations, even outside of the JPN-CDN-FRA lineage (e.g., within CHN; Figure S4). The least correlated populations with the BC wild populations were DPB and GUR (Figure S4).

While some hatcheries may attempt to cross inbred lines for heterosis (leading to low F_IS_), the resulting offspring grown together on a farm tend to be comprised of highly related individuals, due to family structure (Figure 3 and 4). The highest relatedness was observed in DPB, followed by GUR and QDC, then ROS, and finally the rest of the populations (Figure 3). The family structure observed in the HAT/HAT-F samples was also observed using shared microhaplotypes (Figure 4). The HAT-F samples that were less divergent from the NAT samples (i.e., ROS and RSC) also had elevated inter-individual relatedness relative to the NAT populations, although not as high as the more divergent HAT/HAT-F populations (Figure 4). In contrast, NAT and NAT-F samples were all equally dissimilar from each other.

**Fig. 3.**
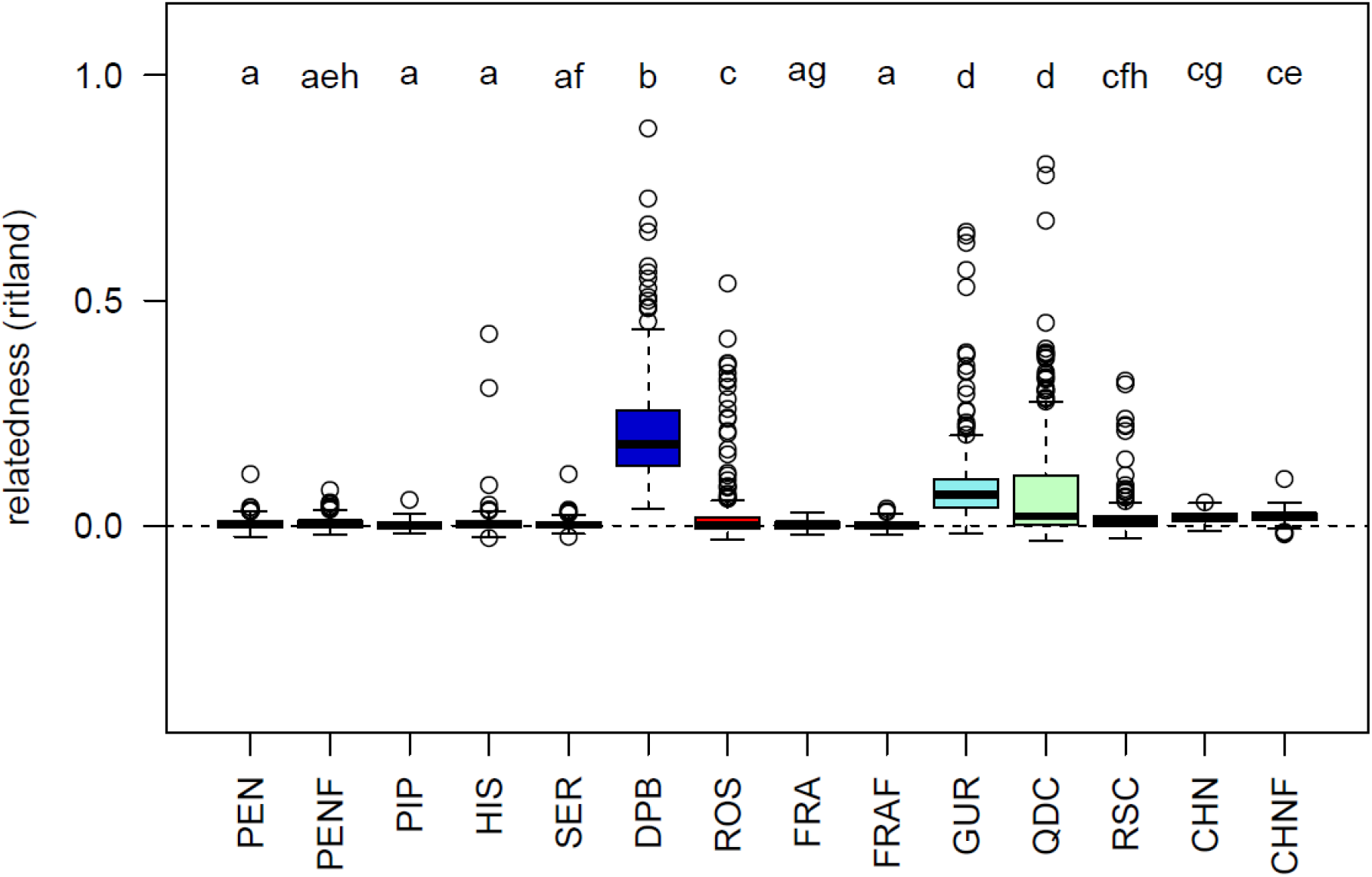
Inter-individual relatedness of individuals within each collection as calculated by the *ritland* statistic (Ritland, 1996; Wang, 2002). Commercial hatchery-farmed (HAT-F) populations include DPB, QDC, ROS, and RSC, and the hatchery population includes GUR. Significant differences (Tukey HSD p ≤ 0.01) are indicated when populations do not share any letters above the boxplot. The collection from JPN is not included in this analysis due to lower sample size and impact of sample size on the statistics.

**Figure 4.**
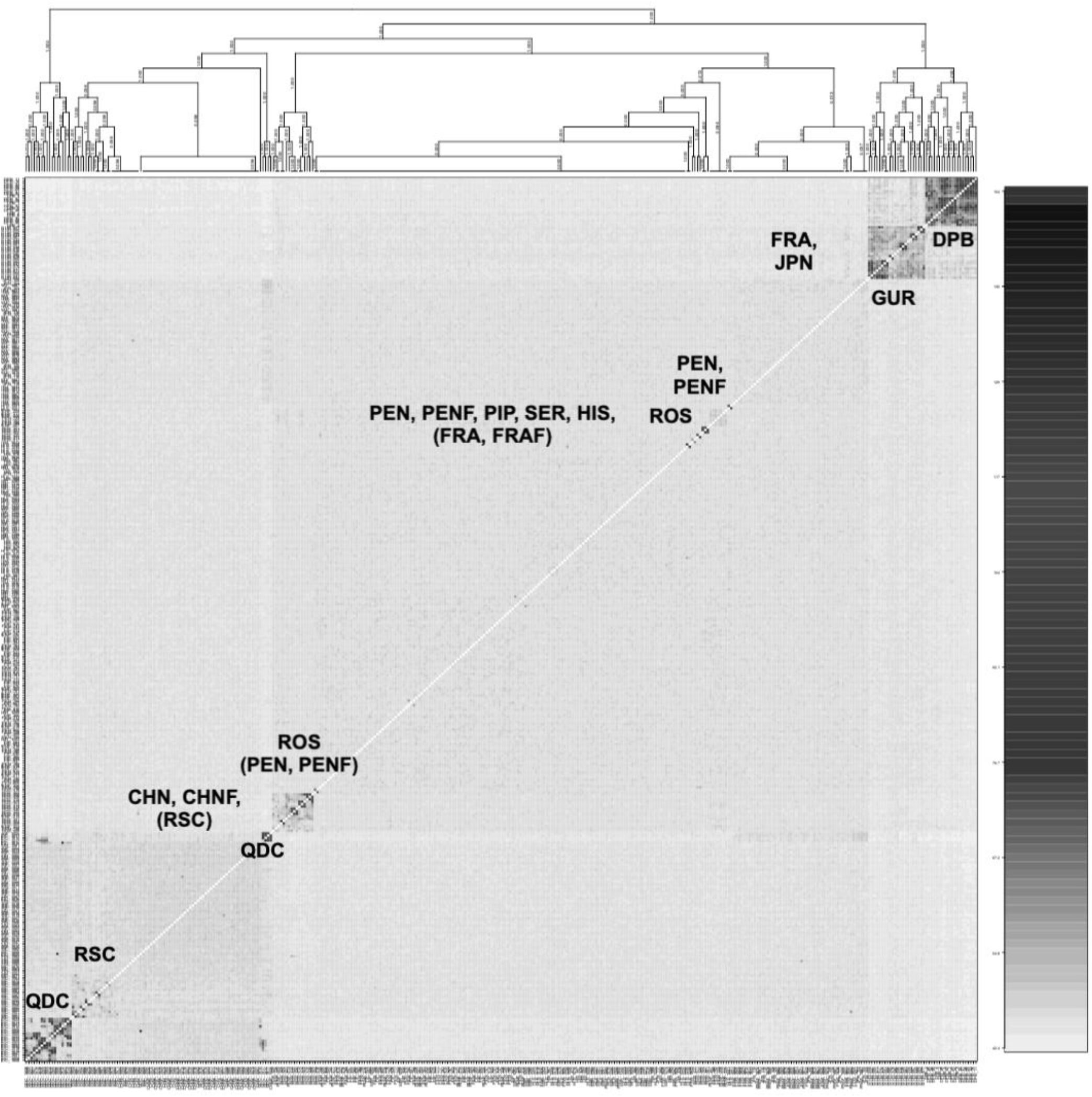
FineRADstructure analysis indicating family structure of hatchery populations and general similarity among groups, as calculated by shared microhaplotypes between samples, including a total of 17,084 loci. Increased shading reflects increased relatedness between individuals.

### Signatures of domestication selection

Here we investigate domestication selection, which we define as any markers that are outliers between HAT/HAT-F oysters and NAT oysters. This could reflect either directional selection or relaxed selection in HAT/HAT-F. Using *pcadapt*, between 82 and 367 outliers (false discovery rate q < 0.01) were identified per contrast between paired populations of HAT/HAT-F oysters and associated NAT populations (see top of Table 2). Using RDA, 170 outliers were identified between HAT/HAT-F and NAT populations. RDA indicated that HAT/HAT-F populations could be significantly separated along RDA1 from nearby NAT populations (Figure S5). Using *BayeScan* no outliers were identified. On average, 133 outliers were identified within the 99^th^ percentile of F_ST_ per contrast. The comparisons involving QDC, DPB, and ROS had the most outliers using *pcadapt*, with fewer identified for GUR and RSC. In general, outliers were found throughout the genome (Figure 5).

**Figure 5.**
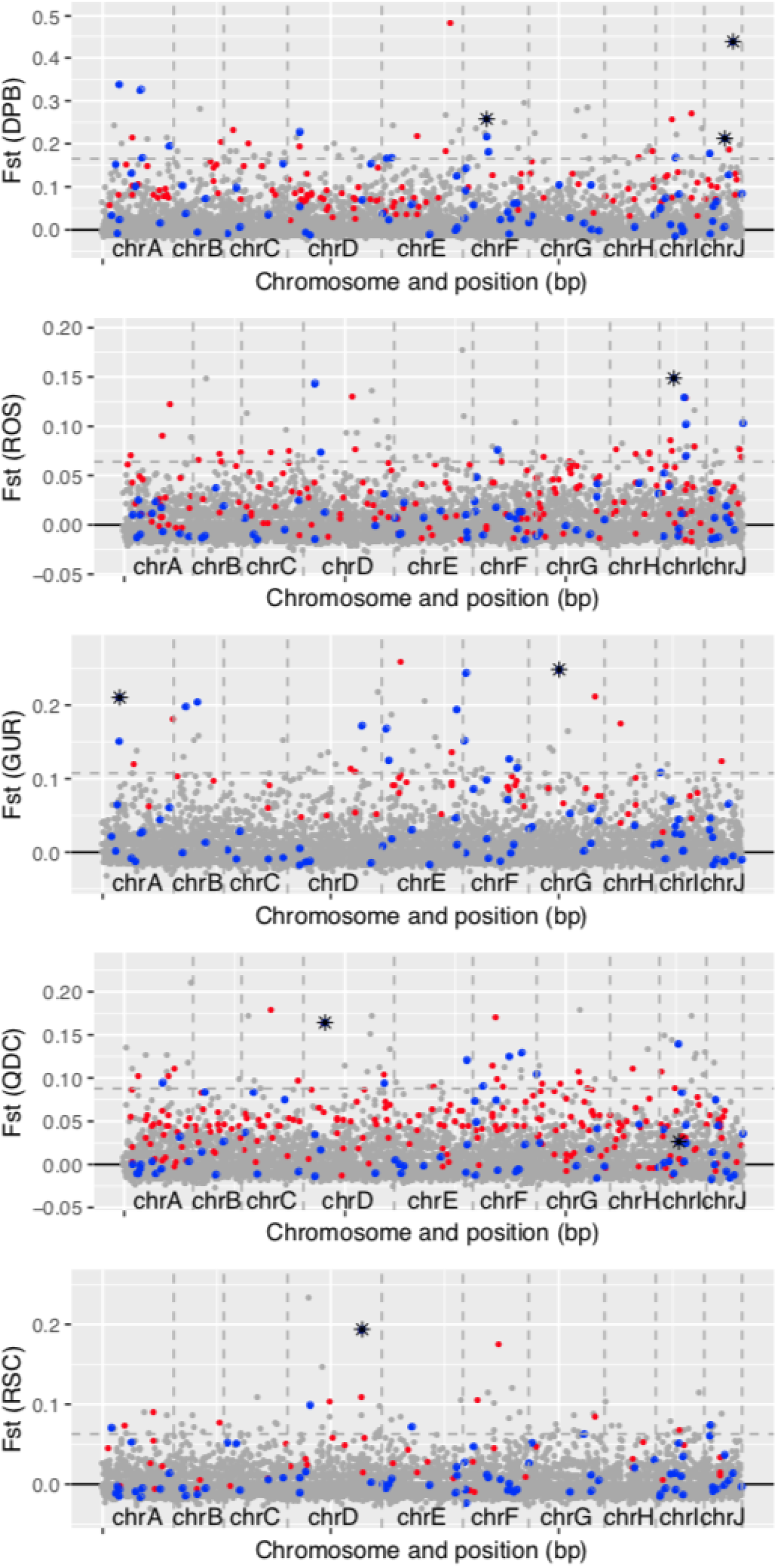
Putative domestication outliers in each comparison shown with per-locus F_ST_ values alongside significant *pcadapt* (red), RDA (blue), and both (black asterisks) indicators. The naturalized/wild population being compared alongside each contrast is the most geographically proximal population (see Table 2). Vertical grey dashed lines separate the chromosomes, and the horizontal dashed line is placed at the 99^th^ percentile of the F_ST_ values for the contrast.

Top outliers were identified as those that had the strongest evidence per comparison (i.e., significant for both *pcadapt* and RDA), or those that were consistently identified in multiple population comparisons with at least two pieces of evidence (i.e., evidence includes significant for F_ST_ 99^th^ percentile, RDA, or *pcadapt*). Top outliers with the strongest evidence and consistent observations included three markers: CLocus_104468, CLocus_625182, and CLocus_704500, which were identified in RSC and GUR, QDC and RSC, and QDC and DPB, respectively. CLocus_104468 was found to be in an intron of *fibrocystin-L*, CLocus_625182 was found to be in an intron of *prion-like-(Q/N-rich) domain bearing protein 25*, and CLocus_704500 was not near any genes. Both CLocus_104468 and CLocus_625182, significant using all three outlier approaches for RSC and QDC, respectively, were found on chrD, at position 32,708,807 bp and 9,881,897 bp, respectively. An additional 10 outliers were identified using two statistical approaches (see Additional File S3), and an additional 17 were found with at least two pieces of any evidence among the multiple population comparisons. Marker names, evidence type, contrasts, and the genomic location including containing or nearby genes (within 10 kb) are shown in Additional File S3 for further investigation. Putative functions and relation to the Pacific oyster and domestication in general are presented in the Discussion (see below).

### Signatures of wild-to-farm movement selection

Signatures of selection were investigated in oysters moved from naturalized populations onto farms (NAT-F). Three parallel comparisons were made from farms and source populations in Canada, France, and China. As expected, oysters that remained in the source population versus moved onto a farm typically exhibited negligible genetic differentiation (see the bottom of Table 2).

Outlier loci were not identifiable for these comparisons with *pcadapt* because scree plots did not indicate any PCs explaining significant proportions of the variation (Additional File S1). RDA found a total of 175 outliers between NAT and NAT-F samples (Figure S6), and these markers were able to differentiate NAT and NAT-F populations (Figure S7). *BayeScan* did not find any outliers. To characterize markers putatively under parallel selection in multiple regions, per locus F_ST_ values were compared among regions (Figure S8). In this way, several markers were found to have F_ST_ values greater than the 99^th^ percentile in pairs of comparisons. However, the amount of parallelism was low, as observed by the lack of markers with high F_ST_ in both comparisons (Figure S8). Two markers were found to be consistent outliers, CLocus_24819 (unknown chromosome) found by RDA and 99^th^ percentile F_ST_ in CHN and BC, and CLocus_608966 (chrB) found in FRA and BC.

## DISCUSSION

The identification of several thousand genetic markers throughout the genome of the Pacific oyster *C. gigas* allowed us to obtain a detailed overview of the genetic differentiation of oysters in populations across the Northern Hemisphere, including whether consistent evidence exists for reduced diversity in farmed populations. There were three major findings that emerged from our study. First, the most distinct populations in terms of genetic differentiation were commercial hatchery farmed populations (HAT-F) as well as populations from China. HAT-F populations often had elevated relatedness as has been observed previously (Hedgecock & Sly, 1990), and in one case had substantially reduced genetic diversity (DPB). There was no indication of large effects from on-farm, within-generation selection, and therefore the differentiation observed in these HAT-F populations is probably due to demographic effects (e.g., founder effects). Second, the genetic structure around Vancouver Island was very low as previously observed (Sun & Hedgecock, 2017). Even further, there was essentially no genome-wide differentiation between sites on either side of Vancouver Island (i.e., Pendrell and Pipestem), two regions commonly used for broodstock collections for farming in BC. Third, oysters from China, both farmed and wild, were distinct from the other oysters in the Japan-Canada-France translocation lineage, providing further evidence to differences between these two groups (Li et al., 2015). In addition, this work has also identified private alleles specific to distinct farm populations, as well as putative domestication outliers, in some case being identified in more than one farm population (Additional File S3).

### Genetic differentiation around Vancouver Island, British Columbia

Very weak to negligible population differentiation was observed across BC, as expected for populations of Pacific oyster (Sun & Hedgecock, 2017), and typical of other marine invertebrates (Gagnaire et al., 2015). Low F_ST_ values may be due to weak genetic drift (large effective population sizes) and/or high dispersal, for example as observed in blue mussel (Bierne, 2010). Given the expected transfer of Pacific oysters by human activity around Vancouver Island, such as the common use of both Pipestem Inlet and Pendrell Sound for farmers to collect seed (BC Shellfish Growers Association, 2019), the low differentiation is not surprising. In fact, these two populations were the only pair whose F_ST_ value was not significantly different from zero, potentially reflecting the reduced genetic differentiation due to increased human-mediated transfer.

Stock identification of oysters from the regions around Vancouver Island would not be possible given the results observed here and previously reported (Sun & Hedgecock, 2017). Increasing sample size drastically for populations or using additional markers putatively under selection could potentially increase power for stock identification (Gagnaire et al., 2015), but given the biology and dispersal ecology of Pacific oyster (i.e., larvae can disperse for up to 20 days at 22°C (Quayle, 1988)) and the anthropogenic movements of stocks, the probability of genetic stock identification of Pacific oysters around coastal BC is very low.

Given the weak differentiation around Vancouver Island, crosses involving oysters sourced from different locations are not expected to confer heterosis. The slight differentiation between Hisnit Inlet and Pendrell Sound could potentially provide this benefit, but without conducting trials of genetically similar or divergent crosses exposed to a variety of environments and perturbations, it is not possible to know if such crosses would generate heterosis. By contrast, crossing oysters from Pendrell Sound and Pipestem Inlet is not expected to provide any heterosis benefits, given the lack of any differentiation between these two areas. The similarity of these areas may change over time however, considering the stochasticity inherent in species with sweepstakes reproductive success, like Pacific oyster.

Spatial effects between the most distant locations by sea outweighed the temporal effects at Pendrell Sound but was not significantly different from the temporal effect for Hisnit Inlet (i.e., the location that is assumed to have more years between the collections based on size differences). The present study uses short time scales and has expected noise that comes with assuming brood year by oyster size (Hedgecock & Pudovkin, 2011). This is in contrast with previous work with decades between brood years that found that temporal variation outweighed spatial variation using a coding gene SNP panel with 52 markers (Sun & Hedgecock, 2017). Sweepstakes reproductive success and chaotic genetic patchiness can cause genetic differentiation to vary over time, with genetic bottlenecks being more pronounced when conditions are less favourable (Hedgecock & Pudovkin, 2011). For example, in sea scallops, *Placopecten magellanicus*, differentiation between different life stages was observed at one site but not at another (Lehnert et al., 2018). The question of spatial compared to temporal variation is therefore expected to depend on oceanic conditions, number of generations involved, and the spatial distance of the comparison.

### Genetic differentiation among countries

There was very low differentiation among distant regions, including Japan, France and Canada, implying little if any bottleneck during these introductions (Sun & Hedgecock, 2017; Gagnaire et al., 2018), potentially due to repeated introductions and high propagule pressure (Gagnaire et al., 2018). The introductions of Pacific oyster to France occurred in 1971-1977 either by transplant of adult oysters from BC (~562 tons (t)) or as spat from Japan (e.g., oyster or scallop shell on which spat had been set) (~10,015 t of spat collectors) (Quayle, 1988; Grizel & Heral, 1991). In contrast, transplantations from Japan to Canada probably stopped in the 1950s (Quayle, 1988). The differentiation among Japan, France and Canada reflects the translocation history and time since introduction, where France is similar to both BC and Japan, and BC is more genetically distant from Japan than BC is from France (Figure 2).

Relative to oysters from Canada, France, and Japan, oysters from China were more genetically distinct (Figure 2). This was previously identified in a comparison of populations around China and Japan, where the authors found no population structure in China and only differentiation between China and Japan (Li et al., 2015). Pacific oysters originate from East Asia, and therefore non-farmed oysters from Japan and China could represent wild oysters, with differences between them pre-dating human use. Although most of the cultured oysters in China are thought to have originally been introduced from Japan (Guo et al., 1999), the close similarity of farmed and wild oysters from China observed in this study indicates either that the farmed oysters in the present study originated from wild populations in China, or that the wild collections have experienced gene flow from the hatchery farmed populations in China, following a period of differentiation from the Japan population. Further work on the East Asian populations is needed to determine the timing and history of genetic differentiation between oysters from China and the JPN-CDN-FRA lineage, as well as the potential role of selective breeding and aquaculture.

### Signatures associated with divergent hatchery populations

Signatures of selection that are different between natural and farmed populations, including markers under domestication selection, can indicate specific genomic regions or markers that have been positively selected during the domestication or selective breeding process (Gutierrez et al., 2016; Gutierrez et al., 2018). Here it is not possible to determine whether the outliers identified were under selection on the farms or hatchery or whether they have experienced relaxed selection relative to natural populations. In any case, the outliers that we identified are good candidates for regions that have been under differential selection between natural and farm populations and therefore might provide information on the different selective pressures in these two locations.

In the current study, the most divergent hatchery populations were those with identifiable family structure, and this was consistent for all countries (e.g., DPB in Canada, GUR in the UK, QDC in China). This may indicate founder effects of the broodstock line, selection within hatcheries or farms, or genetic drift due to low effective population size of the brood line. Interestingly, the most divergent hatchery strain (DPB) also had low nucleotide diversity, which may indicate a lower number of founding individuals or more ongoing drift in this particular lineage. Hatchery propagation is known to potentially impact both the genetic diversity and the allelic composition of Pacific oyster. For example, hatcheries may cull the slowest growing oysters to keep the crop synchronized, but given that this growth rate is in part genetically determined, this imposes selection on the hatchery population and has been shown to reduce diversity and induce temporally varying genetic structure (Taris et al., 2006). Furthermore, many aspects of reproduction can result in large differences in Ne and N in hatchery or wild populations, including gamete quality, sperm-egg interaction, and differential viability among genotypes (Boudry et al., 2002). This is a strong argument to avoid having too few progenitors in breeding programs and to be aware that the total individuals put into the breeding program may not reflect the genetic diversity that results. Although best practices would suggest avoiding extreme differences in reproductive success from individual broodstock parental pairs, this requires specific steps to avoid (e.g., not combining sperm from multiple fathers during fertilization to reduce sperm competition). Given the strong differences between hatchery and wild populations in terms of inter-individual relatedness, and to a lesser extent in nucleotide diversity and polymorphism rate, the impacts of these factors on ecological aspects such as response to environmental perturbation or disease are worthwhile investigating. However, it remains unknown whether farms with lower diversity than others (i.e., lower F_IT_) are less likely to survive perturbations, or whether they have in fact undergone more selection and may be fitter under local conditions. These questions are important to address in future work.

The reduced inbreeding coefficient (F_IS_) in DPB may reflect the practice of crossing inbred lines, possibly with the aim to reduce the negative impact of recessive mutations in the Pacific oyster (Williams, 1975; Launey & Hedgecock, 2001; Ellegren & Galtier, 2016). The lower nucleotide diversity in DPB may reflect either a smaller number of founding individuals, ongoing drift, and/or selection. The lower F_IS_ observed may result in a single generation of heterozygote advantage (Launey & Hedgecock, 2001), but if these individuals cross again within their cohort, they would have overall reduced nucleotide diversity and thus less chance of carrying beneficial alleles for subsequent adaptation. Further, if they introgress into naturalized populations, they would bring this reduced genetic diversity. The amount of introgression from hatchery populations that are highly divergent from naturalized populations is worth further investigation. However, whether this is of concern is dependent on whether conservation of naturalized populations, such as for local broodstock, is a goal, given that the populations outside of Asia are introduced.

Higher inter-individual genetic variation was observed using multivariate approaches (i.e., PCA, DAPC) within farm populations, particularly those with family structure, than within naturalized populations (Figure S1). A similar result was found in selectively bred farmed populations of Atlantic salmon (~12 generation of farming), which showed higher inter-individual variation relative to wild populations (Gutierrez et al., 2016). This finding was attributed to both the admixture that occurred when the farm population was created and the repeated introgression of other lineages into the broodstock line over time. This increased variation was not observed in all farmed populations, with some salmon populations being more similar to wild populations (Gutierrez et al., 2016). This is similar to the differences in inter-individual variance observed in the present study for DPB (more inter-individual variance than naturalized populations) and ROS (similar to naturalized populations). Further, similar to the present study, many populations of farmed Atlantic salmon had low inbreeding values (F_IS_).

Breeding practices can influence the underlying genetics of hatchery strains, which can affect their potential to undergo selective breeding, as well as to have the necessary genetic variation to handle various biotic and abiotic stressors or threats, such as disease, when there is a genotype-environment interaction. An Australian hatchery strain was found to have lower genetic variation but to have less family structure and no sign of inbreeding; other strains in the same study showed higher family effects than this strain, thus this is likely occurring through the careful selection of unrelated individuals for crosses (Kijas et al., 2019). Similarly, in the present study, the RSC population from China, even though it has been bred for at least six generations, shows less family structure than the QDC strain. The QDC strain has a similar level of inter-individual relatedness to the DPB population, but the DPB population has the additional issue of reduced overall nucleotide diversity, indicative of a potential bottleneck or founder effect. This DPB strain would have lower adaptive potential and lower potential to undergo further selective breeding without introgression from other divergent populations.

### Domestication Outliers

The identification of outliers in multiple populations of oysters has highlighted regions of the genome that may have been subject to different selection pressures under domestication. Many outlier markers were near or within genes involved in either calcium signalling or resilience to environmental stressors (see Additional File S3). Calcium signalling may play an important roles in response to abiotic stress and defense. Genes nearby domestication outliers with calcium signalling related functions include *GTP-binding protein REM 1, calcium-binding mitochondrial carrier protein Aralar1-like, IQ domain-containing protein K* (may bind calmodulin), *calmodulin-like, calcineurin subunit B type 1* (stimulated by calmodulin and conferring calcium sensitivity), and *GTP-binding protein RAD* (involved in calmodulin binding) (Additional File S3). Calmodulin is a multifunctional calcium-binding protein, and given that intracellular Ca^2+^ can function in such diverse responses as response to abiotic stressors such as mechanical stimuli, osmotic stress, and heat and cold shock, as well as being involved in defending against pathogenic infections, this general category of calcium-binding genes is of particular note. Other genes involved in cellular stability, such as T-complex protein 1 subunit zeta, mitochondrial uncoupling protein 2, tumor suppressor p53-binding protein 1 were also within this list. The immune-related (antiviral) *toll-like receptor 3* was also found to be near an outlier.

An outlier was found in *transient receptor potential cation channel subfamily M member* 6; interestingly, *member 3-like* from the same cation channel subfamily member was found to be a parallel domestication outlier in both Scottish and Canadian domestic Atlantic salmon (López et al., 2019). Several other genes identified here have also been identified as being involved in domestication processes, such as *mitochondrial uncoupling protein 2*, which has undergone loss of function mutations in the domestication of pigs *(Sus scrofa domesticus)* and potentially has led to the reduction of cold tolerance (Berg et al., 2006), as well as being found to be an outlier in domestic dogs *Canis lupus familiaris* (Udagawa et al., 2014).

The role of these genes in terms of calmodulin function, calcium signalling, and resilience to abiotic stressors may provide insight into the selective differences experienced in the farm environment and the domestication process in Pacific oyster. Continued efforts to understand the differences in selective landscapes, including the genes involved, between hatchery populations and naturalized populations will inform the pressures that the oysters face in both environments.

### Within-generation impacts of transplanting oysters to a farm compared to nature

Within-generation outliers putatively under differential selection in naturalized oysters moved onto farms relative to those remaining in nature were minimal, and showed very low parallelism across regions. Aside from the present study, the effect of within-generation selection associated with moving oysters from nature onto a farm is largely uncharacterized. Sources of selection on farms include more crowding and different substrate conditions (e.g. oyster shell substrate held in mesh bags). Alternatively, the outliers may reflect selection in the oysters that remained in the naturalized populations, with relaxed selection in farmed oysters.

The lack of genome-wide differentiation between the naturalized and the transplanted populations indicates that the sampling of the naturalized population to be moved onto the farm did not impose a bottleneck or founder effect, and that specific families were not greatly enriched on the farm during the transplantation. Given that mesh bags of naturally set oysters are moved onto farms, and that these were sampled in the present study, suggests that oysters naturally settling near one another are not highly related. This is also reflected by the lack of family structure in the NAT-F populations relative to that viewed in the HAT/HAT-F populations.

## Conclusions

Consistent with previous studies, Pacific oysters along the Japan-Canada–France translocation lineage showed very low genetic differentiation. However, Chinese oysters, including wild individuals, were more distinct from this group. This could be due to long-term genetic isolation and/or the effect of farmed oysters sourced from Chinese broodstock introgressing into wild populations. Whether due to founder effects or drift the magnitude of the genetic differentiation between some commercial hatchery populations and naturalized populations were similar to differentiation in the most distinct populations across the sampled Northern Hemisphere range. As previously observed, commercial hatchery populations had more distinct family structure and higher inter-individual relatedness, and one farm exhibited lower genetic diversity. The most differentiated hatchery populations also had a higher prevalence of private alleles. Whether the distinctiveness and low genetic diversity of some hatchery-propagated populations is considered to be a risk for introgression into naturalized populations depends on whether Pacific oyster in Canada is considered a genetic resource for broodstock and a harvestable food resource, or whether it is considered an invasive species. Commonly used broodstock sourcing sites Pendrell Sound and Pipestem Inlet was the only pair of populations that had no genetic differentiation. Valuable next steps would include controlled challenge studies to determine the impact on survival of the candidate loci identified as outliers here and studies to examine the strength of heterosis as a function of the differentiation documented between populations.

## Supporting information

Supplemental Results

## Acknowledgements

This work was funded by the Gordon and Betty Moore Foundation (GBMF#5600). Thanks to all those that contributed samples for this project, including Nicolas Bierne and Ismaël Bernard (Eurêka Modélisation), David Howell, J.P. Hastey (Nova Harvest Ltd.), Keith Reid (Odyssey Shellfish), Brian Yip (Fanny Bay Oysters), Joe Tarnowski (Baynes Sound Oyster co.), and Bruce Evans. Thanks to Nicolas Bierne for comments on the manuscript, to Tim Green for discussion on oyster breeding approaches, to Pierre-Alexandre Gagnaire for providing an early version of the chromosome-level assembly, to Brian Boyle from IBIS for discussions on ddRAD-seq approaches, and to Thierry Gosselin and Eric Normandeau for population genetics discussions. Thanks to the Twitter population genetics communities for discussions around population differentiation in low structured species, domestication selection, and other population genetic concepts. Thanks to the Editor and two anonymous reviewers for valuable comments on an earlier version of the manuscript.

## Data Availability

RAD-seq protocol: https://www.protocols.io/view/nsil1-msp1-rad-digest-detailed-protocol-2019-01-18-w95fh86

Complete analysis pipeline: https://github.com/bensutherland/ms_oyster_popgen

Genotyping pipeline (original): https://github.com/enormandeau/stacks_workflow (see fork: https://github.com/bensutherland/stacks_workflow)

Raw data: NCBI Short Read Archive (SRA) BioProject PRJNA550437

## Supplemental Materials

**Table S1.** Mean genetic differentiation (F_ST_) shown between pairs of populations for **(A)** populations in BC; and **(B)** populations in BC separated by size class; and (C) global populations.

**Table S2.** Population summary statistics with standard error for the single SNP per marker data from *Stacks* using either **(A)** all variant sites; or **(B)** all variant and fixed RAD loci.

**Figure S1.** Principal Components Analysis (PCA) of all samples using a single SNP per locus with all loci. Sample IDs are plotted along **(A)** PC 1 and 2; or **(B)** PC3 and 2.

**Figure S2.** Discriminant Analysis of Principal Components (DAPC) marker loading values.

**Figure S3.** Per population distribution of nucleotide diversity (π), shown as frequencies of specific bins of nucleotide diversity as calculated by vcftools (--site-pi).

**Figure S4.** Nucleotide diversity (π) for each marker (dot) correlated between each population with the BC wild population.

**Figure S5.** Comparison between Redundancy Analysis (RDA) and PCA in differentiating populations.

**Figure S6.** Outliers identified between naturalized/wild oysters moved onto a farm and naturalized populations (NAT-F vs. NAT) using RDA analysis.

**Figure S7.** Comparison between Redundancy Analysis (RDA) and PCA in differentiating populations.

**Figure S8.** Per locus F_ST_ values compared between different countries to detect signatures of parallel selection from being moved onto a farm.

**Additional File S1.** Scree plot showing percent variation explained for all domestication or on-farm selection contrasts.

**Additional File S2.** Score plots showing sample position within PCA based on each PC for all domestication or on-farm selection contrasts.

**Additional File S3.** Top outliers, either those identified as being an outlier via both *pcadapt* and RDA, or those with at least two outlier tests (*pcadapt*, RDA, 99^th^ percentile F_ST_) and present in more than one comparison. Information such as marker name, chromosome-level assembly alignment chromosome where possible, and nearby genes in the reference assembly annotation are provided. Additionally, the general known functions of the outliers from resources such as UniProt are provided for each gene nearby the outlier markers.

## REFERENCES

Andrews, S. (2010). FastQC: A quality control tool for high throughput sequence data. Retrieved from https://www.bioinformatics.babraham.ac.uk/projects/fastqc/

Anglès d’Auriac, M. B., Rinde, E., Norling, P., Lapègue, S., Staalstrøm, A., Hjermann, D. Ø., & Thaulow, J. (2017). Rapid expansion of the invasive oyster Crassostrea gigas at its northern distribution limit in Europe: Naturally dispersed or introduced? PLoS ONE, 12(5), e0177481–e0177481. doi:10.1371/journal.pone.0177481

Arbelaez, J. D., Dwiyanti, M. S., Tandayu, E., Llantada, K., Jarana, A., Ignacio, J. C., Platten, J. D., Cobb, J., Rutkoski, J. E., Thomson, M. J., & Kretzschmar, T. (2019). 1k-RiCA (1K-Rice Custom Amplicon) a novel genotyping amplicon-based SNP assay for genetics and breeding applications in rice. Rice, 12(1), 55. doi:10.1186/s12284-019-0311-0

Association, B. S. G. (2019). Oysters. Retrieved from http://bcsga.ca/shellfish-farming-101/shellfish-we-farm/oysters/

Barrett, R. D. H., & Schluter, D. (2008). Adaptation from standing genetic variation. Trends in Ecology & Evolution, 23(1), 38–44. doi:papers3://publication/doi/10.1016/j.tree.2007.09.008

Beacham, T. D., Wallace, C., MacConnachie, C., Jonsen, K., McIntosh, B., Candy, J. R., Devlin, R. H., & Withler, R. E. (2017). Population and individual identification of coho salmon in British Columbia through parentage-based tagging and genetic stock identification: an alternative to coded-wire tags. Canadian Journal of Fisheries and Aquatic Sciences, 74(9), 1391–1410. doi:10.1139/cjfas-2016-0452

Benestan, L., Gosselin, T., Perrier, C., Sainte-Marie, B., Rochette, R., & Bernatchez, L. (2015). RAD genotyping reveals fine-scale genetic structuring and provides powerful population assignment in a widely distributed marine species, the American lobster (Homarus americanus). Molecular Ecology, 24(13), 3299–3315. doi:papers3://publication/doi/10.1111/mec.13245

Berg, F., Gustafson, U., & Andersson, L. (2006). The uncoupling protein 1 gene (UCP1) is disrupted in the pig lineage: a genetic explanation for poor thermoregulation in piglets. PLOS Genetics, 2(8), e129. doi:10.1371/journal.pgen.0020129

Bierne, N. (2010). The distinctive footprints of local hitchhiking in a varied environment and global hitchhiking in a subdivided population. Evolution, 64(11), 3254–3272. doi:10.1111/j.1558-5646.2010.01050.x

Boudry, P., Collet, B., Cornette, F., Hervouet, V., & Bonhomme, F. (2002). High variance in reproductive success of the Pacific oyster (Crassostrea gigas, Thunberg) revealed by microsatellite-based parentage analysis of multifactorial crosses. Aquaculture, 204(3), 283–296. doi:https://doi.org/10.1016/S0044-8486(01)00841-9

Catchen, J. M., Amores, A., Hohenlohe, P., Cresko, W., & Postlethwait, J. H. (2011). Stacks: building and genotyping loci de novo from short-read sequences. G3 - Genes|Genomes|Genetics, 1(3), 171–182. doi:papers3://publication/doi/10.1534/g3.111.000240

Danecek, P., Auton, A., Abecasis, G., Albers, C. a., Banks, E., DePristo, M. a., Handsaker, R. E., Lunter, G., Marth, G. T., Sherry, S. T., McVean, G., Durbin, R., & Genomes Project Analysis, G. (2011). The variant call format and VCFtools. Bioinformatics, 27(15), 2156–2158. doi:papers3://publication/doi/10.1093/bioinformatics/btr330

De Silva, S. S. (2012). Aquaculture: a newly emergent food production sector—and perspectives of its impacts on biodiversity and conservation. Biodiversity and Conservation, 21(12), 3187–3220. doi:papers3://publication/doi/10.1007/s10531-012-0360-9

Eldon, B., Riquet, F., Yearsley, J., Jollivet, D., & Broquet, T. (2016). Current hypotheses to explain genetic chaos under the sea. Current Zoology, 62(6), 551–566. doi:papers3://publication/doi/10.1093/cz/zow094

Ellegren, H., & Galtier, N. (2016). Determinants of genetic diversity. Nature Reviews Genetics, 1–13.doi:papers3://publication/doi/10.1038/nrg.2016.58

Ewels, P., Magnusson, M., Lundin, S., & Käller, M. (2016). MultiQC: summarize analysis results for multiple tools and samples in a single report. Bioinformatics, 32(19), 3047–3048. doi:papers3://publication/doi/10.1093/bioinformatics/btw354

Foll, M., & Gaggiotti, O. (2008). A Genome-Scan Method to Identify Selected Loci Appropriate for Both Dominant and Codominant Markers: A Bayesian Perspective. Genetics, 180(2), 977. doi:10.1534/genetics.108.092221

Forester, B. R. (2019). Detecting multilocus adaptation using Redundancy Analysis (RDA). Population Genetics in R. Retrieved from https://popgen.nescent.org/2018-03-27_RDA_GEA.html

Forester, B. R., Lasky, J. R., Wagner, H. H., & Urban, D. L. (2018). Comparing methods for detecting multilocus adaptation with multivariate genotype–environment associations. Molecular Ecology, 27(9), 2215–2233. doi:10.1111/mec.14584

Gagnaire, P.-A., Broquet, T., Aurelle, D., Viard, F., Souissi, A., Bonhomme, F., Arnaud-Haond, S., & Bierne, N. (2015). Using neutral, selected, and hitchhiker loci to assess connectivity of marine populations in the genomic era. Evolutionary Applications, 8(8), 769–786. doi:papers3://publication/doi/10.1111/eva.12288

Gagnaire, P.-A., Lamy, J.-B., Cornette, F., Heurtebise, S., Dégremont, L., Flahauw, E., Boudry, P., Bierne, N., & Lapègue, S. (2018). Analysis of genome-wide differentiation between native and introduced populations of the cupped oysters Crassostrea gigas and Crassostrea angulata. Genome Biology and Evolution, 10(9), 2518–2534. doi:papers3://publication/doi/10.1093/gbe/evy194

Goudet, J. (2005). hierfstat, a package for R to compute and test hierarchical F-statistics. Molecular Ecology Notes, 5, 184–186. doi:papers3://publication/doi/10.1111/j.1471-8278

Grizel, H., & Heral, M. (1991). Introduction into France of the Japanese oyster (Crassostrea gigas). ICES Journal of Marine Science, 47(3), 399–403. doi:papers3://publication/doi/10.1093/icesjms/47.3.399

Guo, X. (2009). Use and exchange of genetic resources in molluscan aquaculture. Reviews in Aquaculture, 1(3-4), 251–259. doi:papers3://publication/doi/10.1111/j.1753-5131.2009.01014.x

Guo, X., Ford, S. E., & Zhang, F. (1999). Molluscan aquaculture in China. Journal of Shellfish Research, 18(1), 19–31.

Gutierrez, A. P., Matika, O., Bean, T. P., & Houston, R. D. (2018). Genomic Selection for Growth Traits in Pacific Oyster (Crassostrea gigas): Potential of Low-Density Marker Panels for Breeding Value Prediction. Frontiers in Genetics, 9, 391.

Gutierrez, A. P., Turner, F., Gharbi, K., Talbot, R., Lowe, N. R., Peñaloza, C., McCullough, M., Prodöhl, P. A., Bean, T. P., & Houston, R. D. (2017). Development of a Medium Density Combined-Species SNP Array for Pacific and European Oysters (<em>Crassostrea gigas</em> and <em>Ostrea edulis</em>). G3: Genes|Genomes|Genetics, 7(7), 2209. doi:10.1534/g3.117.041780

Gutierrez, A. P., Yáñez, J. M., & Davidson, W. S. (2016). Evidence of recent signatures of selection during domestication in an Atlantic salmon population. Marine Genomics, 26, 41–50. doi:papers3://publication/doi/10.1016/j.margen.2015.12.007

Harrang, E., Lapègue, S., Morga, B., & Bierne, N. (2013). A high load of non-neutral amino-acid polymorphisms explains high protein diversity despite moderate effective population size in a marine bivalve with sweepstakes reproduction. G3 - Genes|Genomes|Genetics, 3(2), 333–341. doi:papers3://publication/doi/10.1534/g3.112.005181

Hedgecock, D., & Davis, J. P. (2007). Heterosis for yield and crossbreeding of the Pacific oyster Crassostrea gigas. Aquaculture, 272, S17–S29. doi:https://doi.org/10.1016/j.aquaculture.2007.07.226

Hedgecock, D., Gaffney, P. M., Goulletquer, P., & Guo, X. (2005). The case for sequencing the Pacific oyster genome. Journal of Shellfish &. doi:papers3://publication/doi/10.2983/0730-8000(2005)24%255B429:tcfstp%255D2.0.co;2

Hedgecock, D., & Pudovkin, A. I. (2011). Sweepstakes reproductive success in highly fecund marine fish and shellfish: a review and commentary. Bulletin of Marine Science, 87(4), 971–1002. doi:papers3://publication/doi/10.5343/bms.2010.1051

Hedgecock, D., & Sly, F. (1990). Genetic drift and effective population sizes of hatchery-propagated stocks of the Pacific oyster, *Crassostrea gigas*. Aquaculture, 88(1), 21–38. doi:https://doi.org/10.1016/0044-8486(90)90316-F

Herbert, R. J. H., Humphreys, J., Davies, C. J., Roberts, C., Fletcher, S., & Crowe, T. P. (2016). Ecological impacts of non-native Pacific oysters (Crassostrea gigas) and management measures for protected areas in Europe. Biodiversity and Conservation, 25(14), 2835–2865. doi:papers3://publication/doi/10.1007/s10531-016-1209-4

Hubert, S. (2004). Linkage Maps of Microsatellite DNA Markers for the Pacific Oyster Crassostrea gigas. Genetics, 168(1), 351–362. doi:papers3://publication/doi/10.1534/genetics.104.027342

Jombart, T. (2008). adegenet: a R package for the multivariate analysis of genetic markers. Bioinformatics, 24(11), 1403–1405. doi:papers3://publication/doi/10.1093/bioinformatics/btn129

Kamvar, Z. N., Tabima, J. F., & Grünwald, N. J. (2014). Poppr: an R package for genetic analysis of populations with clonal, partially clonal, and/or sexual reproduction. PeerJ, 2, e281. doi:10.7717/peerj.281

Kijas, J. W., Gutierrez, A. P., Houston, R. D., McWilliam, S., Bean, T. P., Soyano, K., Symonds, J. E., King, N., Lind, C., & Kube, P. (2019). Assessment of genetic diversity and population structure in cultured Australian Pacific oysters. Animal Genetics, 50(6), 686–694. doi:10.1111/age.12845

Kraemer, P., & Gerlach, G. (2017). Demerelate: calculating interindividual relatedness for kinship analysis based on codominant diploid genetic markers using R. Molecular Ecology Resources, 17(6), 1371–1377. doi:10.1111/1755-0998.12666

Laporte, M., Pavey, S. A., Rougeux, C., Pierron, F., Lauzent, M., Budzinski, H., Labadie, P., Geneste, E., Couture, P., Baudrimont, M., & Bernatchez, L. (2016). RAD sequencing reveals within-generation polygenic selection in response to anthropogenic organic and metal contamination in North Atlantic Eels. Molecular Ecology, 25(1), 219–237. doi:10.1111/mec.13466

Launey, S., & Hedgecock, D. (2001). High genetic load in the Pacific oyster Crassostrea gigas. Genetics, 159(1), 255–265. doi:papers3://publication/uuid/3F859A91-47C3-4FE1-99BF-5DDEA789B9AB

Lehnert, S. J., DiBacco, C., Van Wyngaarden, M., Jeffery, N. W., Lowen, J., Sylvester, E. V. A., Wringe, B. F., Stanley, R. R. E., Hamilton, L. C., & Bradbury, I. R. (2018). Fine-scale temperature-associated genetic structure between inshore and offshore populations of sea scallop (Placopecten magellanicus). Heredity, 1–12. doi:papers3://publication/doi/10.1038/s41437-018-0087-9

Lepais, O., & Weir, J. T. (2014). SimRAD: an R package for simulation-based prediction of the number of loci expected in RADseq and similar genotyping by sequencing approaches. Molecular Ecology Resources, 14(6), 1314–1321. doi:papers3://publication/doi/10.1111/1755-0998.12273

Li, H., Handsaker, B., Wysoker, A., Fennell, T., Ruan, J., Homer, N., Marth, G., Abecasis, G., Durbin, R., & Genome Project Data Processing, S. (2009). The Sequence Alignment/Map format and SAMtools. Bioinformatics, 25(16), 2078–2079. doi:papers3://publication/doi/10.1093/bioinformatics/btp352

Li, S., Li, Q., Yu, H., Kong, L., & Liu, S. (2015). Genetic variation and population structure of the Pacific oyster *Crassostrea gigas* in the northwestern Pacific inferred from mitochondrial COI sequences. Fisheries Science, 81(6), 1071–1082. doi:10.1007/s12562-015-0928-x

López, M. E., Benestan, L., Moore, J.-S., Perrier, C., Gilbey, J., Di Genova, A., Maass, A., Diaz, D., Lhorente, J.-P., Correa, K., Neira, R., Bernatchez, L., & Yáñez, J. M. (2019). Comparing genomic signatures of domestication in two Atlantic salmon (Salmo salar L.) populations with different geographical origins. Evolutionary Applications, 12(1), 137–156. doi:10.1111/eva.12689

Luu, K., Bazin, E., & Blum, M. G. B. (2017). pcadapt: an R package to perform genome scans for selection based on principal component analysis. Molecular Ecology Resources, 17(1), 67–77. doi:10.1111/1755-0998.12592

Malinsky, M., Trucchi, E., Lawson, D., & Falush, D. (2016). RADpainter and fineRADstructure: population inference from RADseq data. 1–6. doi:papers3://publication/doi/10.1101/057711

Martin, M. (2011). Cutadapt removes adapter sequences from high-throughput sequencing reads. EMBnet journal, 17(1), 10. doi:papers3://publication/doi/10.14806/ej.17.1.200

Mascher, M., Wu, S., Amand, P. S., Stein, N., & Poland, J. (2013). Application of genotyping-by-sequencing on semiconductor sequencing platforms: a comparison of genetic and reference-based marker ordering in barley. PLoS ONE, 8(10), e76925. doi:papers3://publication/doi/10.1371/journal.pone.0076925

Meek, M. H., & Larson, W. A. (2019). The future is now: Amplicon sequencing and sequence capture usher in the conservation genomics era. Molecular Ecology Resources, 19(4), 795–803. doi:10.1111/1755-0998.12998

Oksanen, J., Blanchet, G., Friendly, M., Kindt, R., Legendre, P., McGlinn, D., Minchin, P. R., O’Hara, R. B., Simpson, G. L., Solymos, P., Stevens, H. H., Szoecs, E., & Wagner, H. (2019). vegan: Community Ecology Package (Version 2.5-5). Retrieved from https://CRAN.R-project.org/package=vegan

Pew, J., Muir, P. H., Wang, J., & Frasier, T. R. (2015). related: an R package for analysing pairwise relatedness from codominant molecular markers. Molecular Ecology Resources, 15(3), 557–561. doi:10.1111/1755-0998.12323

Plough, L. V. (2016). Genetic load in marine animals: a review. Current Zoology, 62(6), 567–579.doi:papers3://publication/doi/10.1093/cz/zow096

Quayle, D. B. (1988). Pacific oyster culture in British Columbia (Vol. 218). Nanaimo, British Columbia: Fisheries and Oceans Canada.

Queller, D. C., & Goodnight, K. F. (1989). Estimating relatedness using genetic markers. Evolution, 43(2), 258–275. doi:10.1111/j.1558-5646.1989.tb04226.x

Recknagel, H., Jacobs, A., Herzyk, P., & Elmer, K. R. (2015). Double-digest RAD sequencing using Ion Proton semiconductor platform (ddRADseq-ion) with nonmodel organisms. Molecular Ecology Resources, 15(6), 1316–1329. doi:papers3://publication/doi/10.1111/1755-0998.12406

Reise, K., Buschbaum, C., Büttger, H., Rick, J., & Wegner, K. M. (2017). Invasion trajectory of Pacific oysters in the northern Wadden Sea. Marine biology, 164(4), 68–68. doi:10.1007/s00227-017-3104-2

Ritland, K. (1996). Estimators for pairwise relatedness and individual inbreeding coefficients. Genetical Research, 67(2), 175–185. doi:10.1017/S0016672300033620

Rochette, N. C., Rivera-Colón, A. G., & Catchen, J. M. (2019). Stacks 2: analytical methods for paired-end sequencing improve RADseq-based population genomics. bioRxiv. doi:papers3://publication/doi/10.1101/615385

Storey, J. D., Bass, A. J., Dabney, A., & Robinson, D. (2019). qvalue: Q-value estimation for false discovery rate control. (Version 2.16.0). Retrieved from http://github.com/jdstorey/qvalue

Storey, J. D., & Tibshirani, R. (2003). Statistical significance for genomewide studies. Proceedings of the National Academy of Sciences, 100(16), 9440. doi:10.1073/pnas.1530509100

Sun, D., & Hallin, L. (2018). BC Ministry of Agriculture - British Columbia’s Fisheries and Aquaculture Sector, 2016 Edition. British Columbia: BC Stats Retrieved from https://www2.gov.bc.ca/assets/gov/farming-natural-resources-and-industry/agriculture-and-seafood/statistics/industry-and-sector-profiles/sector-reports/british_columbias_fisheries_and_aquaculture_sector_2016_edition.pdf.

Sun, X., & Hedgecock, D. (2017). Temporal genetic change in North American Pacific oyster populations suggests caution in seascape genetics analyses of high gene-flow species. Marine Ecology Progress Series, 565, 79–93. doi:papers3://publication/doi/10.3354/meps12009

Tange, O. (2011). GNU Parallel - The Command-Line Power Tool. The USENIX Magazine, 36(1), 42–47.doi:papers3://publication/doi/http://dx.doi.org/10.5281/zenodo.16303

Taris, N., Ernande, B., McCombie, H., & Boudry, P. (2006). Phenotypic and genetic consequences of size selection at the larval stage in the Pacific oyster (Crassostrea gigas). Journal of Experimental Marine Biology and Ecology, 333(1), 147–158. doi:https://doi.org/10.1016/j.jembe.2005.12.007

Udagawa, C., Tada, N., Asano, J., Ishioka, K., Ochiai, K., Bonkobara, M., Tsuchida, S., & Omi, T. (2014). The genetic association study between polymorphisms in uncoupling protein 2 and uncoupling protein 3 and metabolic data in dogs. BMC Research Notes, 7(1), 904. doi:10.1186/1756-0500-7-904

Vendrami, D. L. J., Houston, R. D., Gharbi, K., Telesca, L., Gutierrez, A. P., Gurney-Smith, H., Hasegawa, N., Boudry, P., & Hoffman, J. I. (2018). Detailed insights into pan-European population structure and inbreeding in wild and hatchery Pacific oysters (Crassostrea gigas) revealed by genome-wide SNP data. Evolutionary Applications. doi:papers3://publication/doi/10.1111/eva.12736

Villacorta-Rath, C., Souza, C. A., Murphy, N. P., Green, B. S., Gardner, C., & Strugnell, J. M. (2017). Temporal genetic patterns of diversity and structure evidence chaotic genetic patchiness in a spiny lobster. Molecular Ecology, 27(1), 54–65. doi:papers3://publication/doi/10.1111/mec.14427

Wang, J. (2002). An Estimator for Pairwise Relatedness Using Molecular Markers. Genetics, 160(3), 1203–1215.

Weir, B. S., & Cockerham, C. C. (1984). Estimating F-statistics for the analysis of population structure. Evolution, 38(6), 1358–1370. doi:10.1111/j.1558-5646.1984.tb05657.x

Wijsman, J. W. M., Troost, K., Fang, J., & Roncarati, A. (2019). Global Production of Marine Bivalves. Trends and Challenges. In A. C. Smaal, J. G. Ferreira, J. Grant, J. K. Petersen, & Ø. Strand (Eds.), Goods and Services of Marine Bivalves (pp. 7–26). Cham: Springer International Publishing.

Williams, G. (1975). Sex and evolution. Princeton: Princeton University Press.

Wringe, B. F., Anderson, E. C., Jeffery, N. W., Stanley, R. R. E., & Bradbury, I. R. (2018). Development and evaluation of SNP panels for the detection of hybridization between wild and escaped Atlantic salmon (Salmo salar) in the western Atlantic. Canadian Journal of Fisheries and Aquatic Sciences, 76(5), 695–704. doi:10.1139/cjfas-2017-0394

Zhang, G., Fang, X., Guo, X., Li, L., Luo, R., Xu, F., Yang, P., Zhang, L., Wang, X., Qi, H., Xiong, Z., Que, H., Xie, Y., Holland, P. W. H., Paps, J., Zhu, Y., Wu, F., Chen, Y., Wang, J., Peng, C., Meng, J., Yang, L., Liu, J., Wen, B., Zhang, N., Huang, Z., Zhu, Q., Feng, Y., Mount, A., Hedgecock, D., Xu, Z., Liu, Y., Domazet-Lošo, T., Du, Y., Sun, X., Zhang, S., Liu, B., Cheng, P., Jiang, X., Li, J., Fan, D., Wang, W., Fu, W., Wang, T., Wang, B., Zhang, J., Peng, Z., Li, Y., Li, N., Wang, J., Chen, M., He, Y., Tan, F., Song, X., Zheng, Q., Huang, R., Yang, H., Du, X., Chen, L., Yang, M., Gaffney, P. M., Wang, S., Luo, L., She, Z., Ming, Y., Huang, W., Zhang, S., Huang, B., Zhang, Y., Qu, T., Ni, P., Miao, G., Wang, J., Wang, Q., Steinberg, C. E. W., Wang, H., Li, N., Qian, L., Zhang, G., Li, Y., Yang, H., Liu, X., Wang, J., Yin, Y., & Wang, J. (2012). The oyster genome reveals stress adaptation and complexity of shell formation. Nature, 490(7418), 49–54. doi:papers3://publication/doi/10.1038/nature11413

