## Supplementary figures and images for "Relative genomic impacts of translocation history, hatchery practices, and farm selection in Pacific oyster *Crassostrea gigas* throughout the Northern Hemisphere"

### Additional File S1

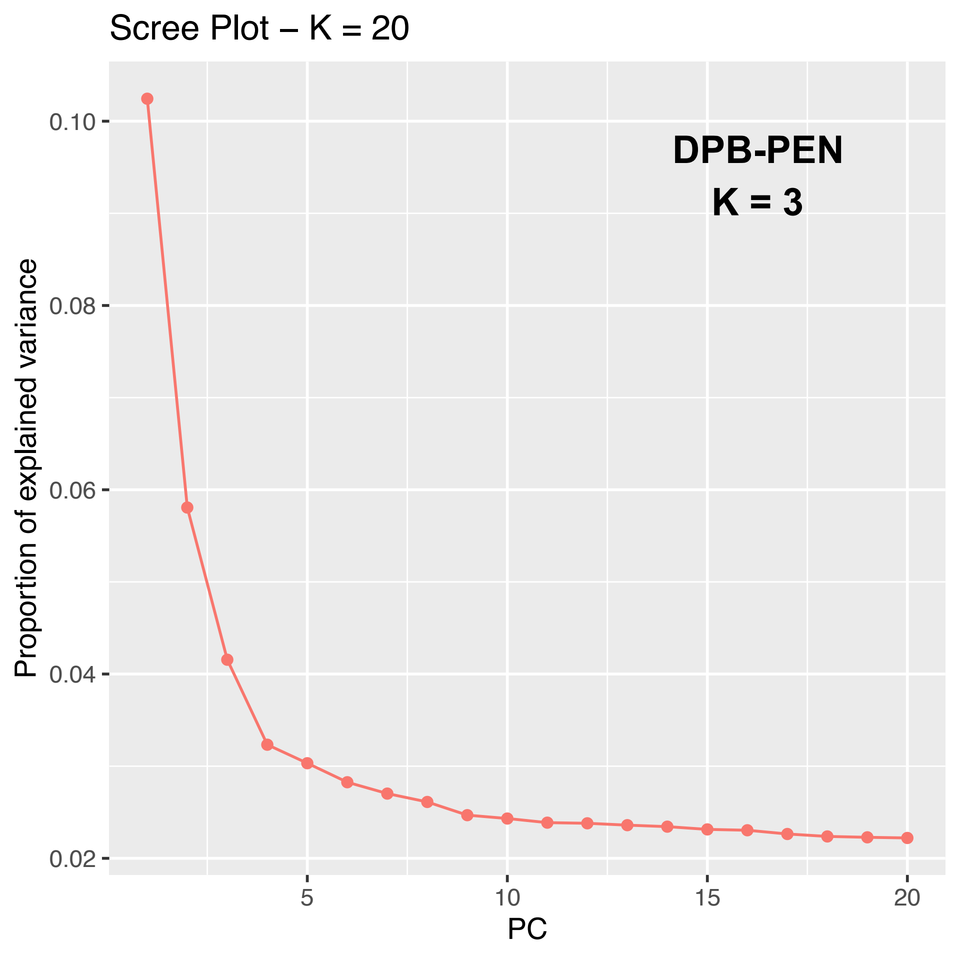

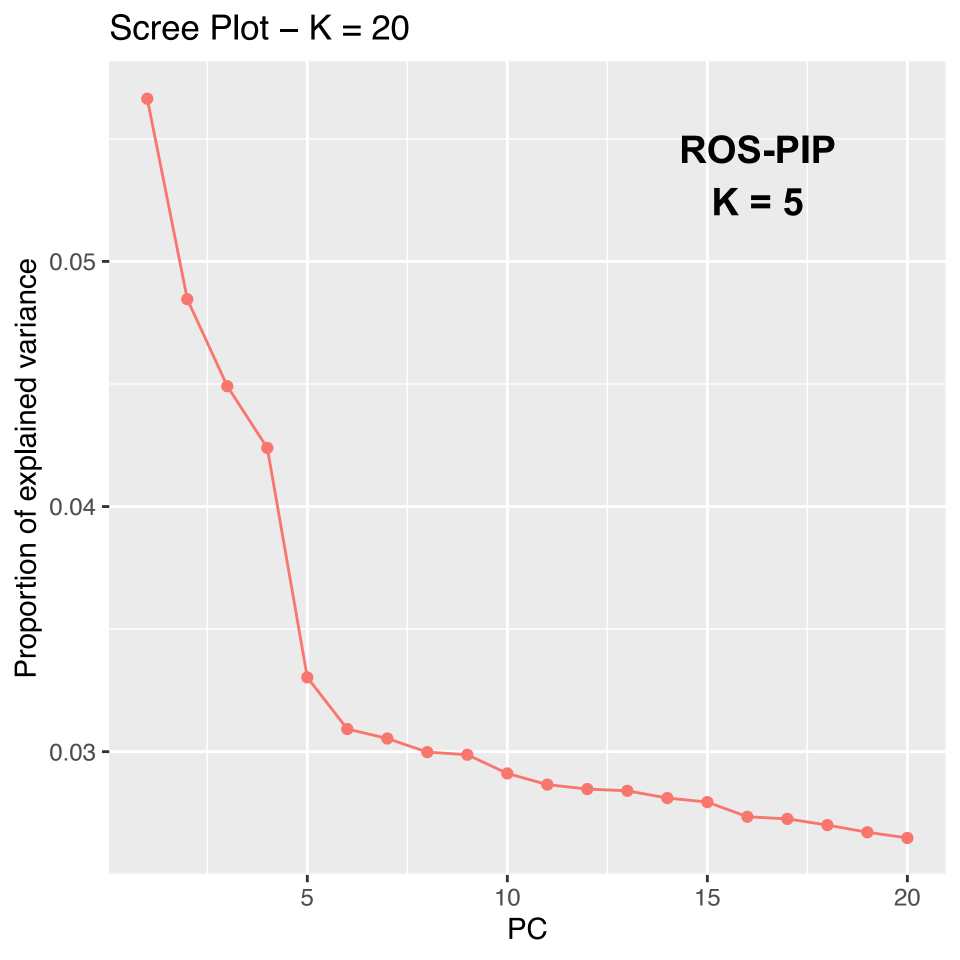

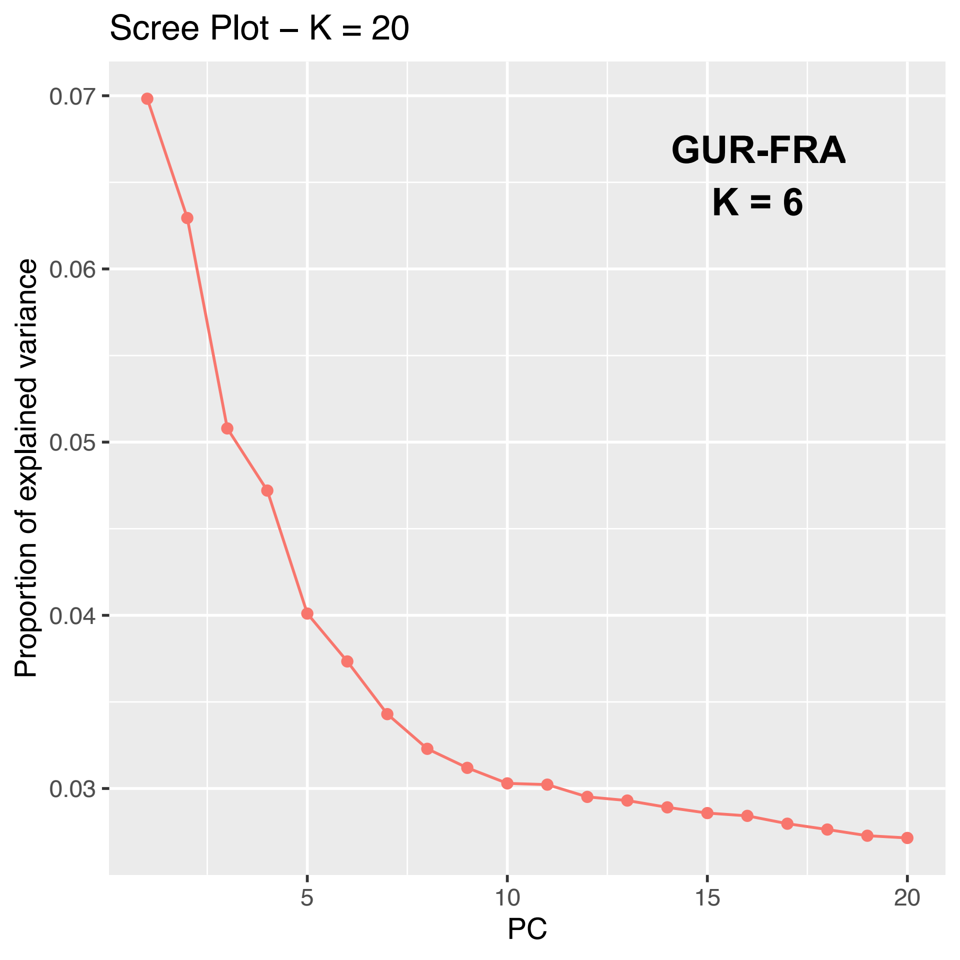

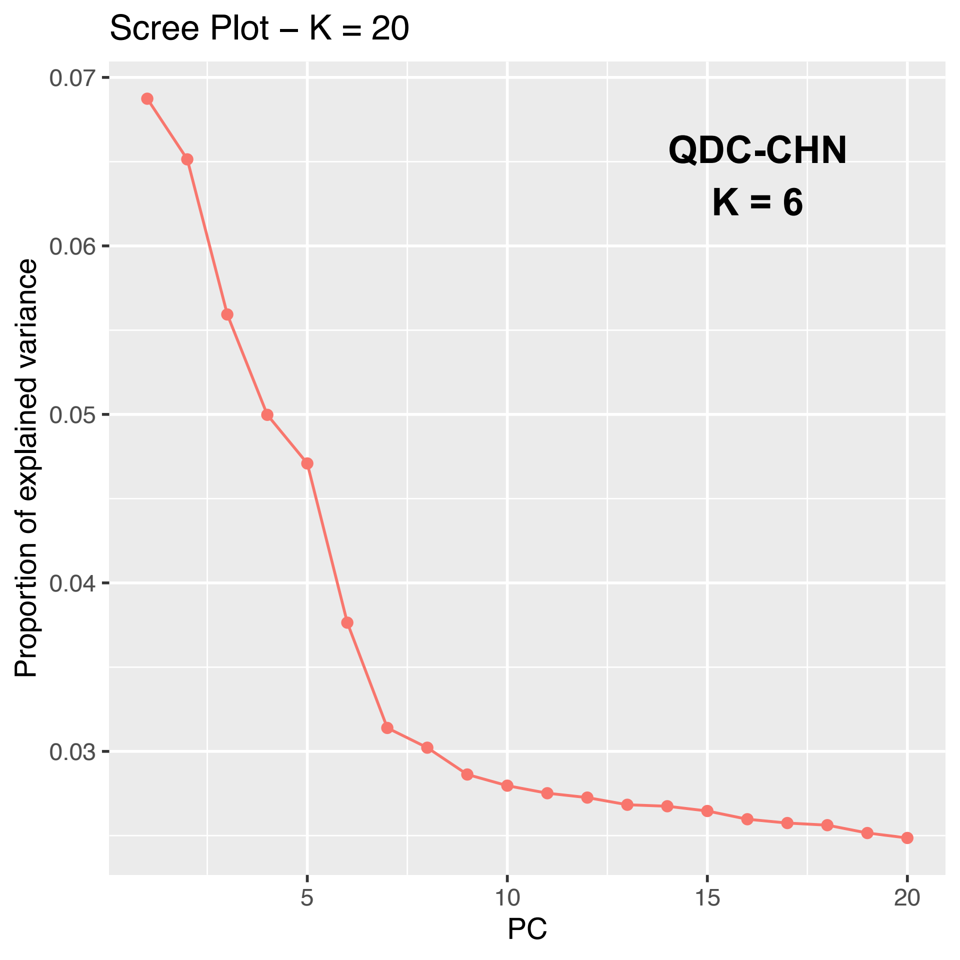

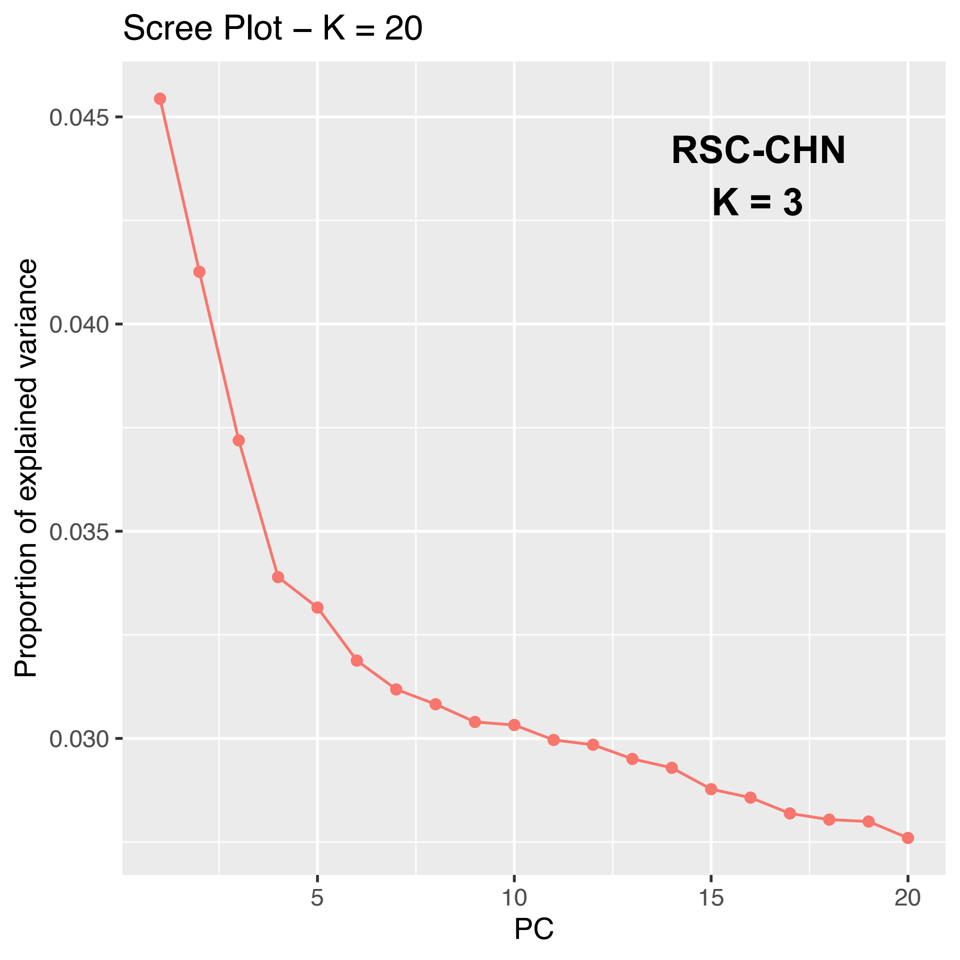

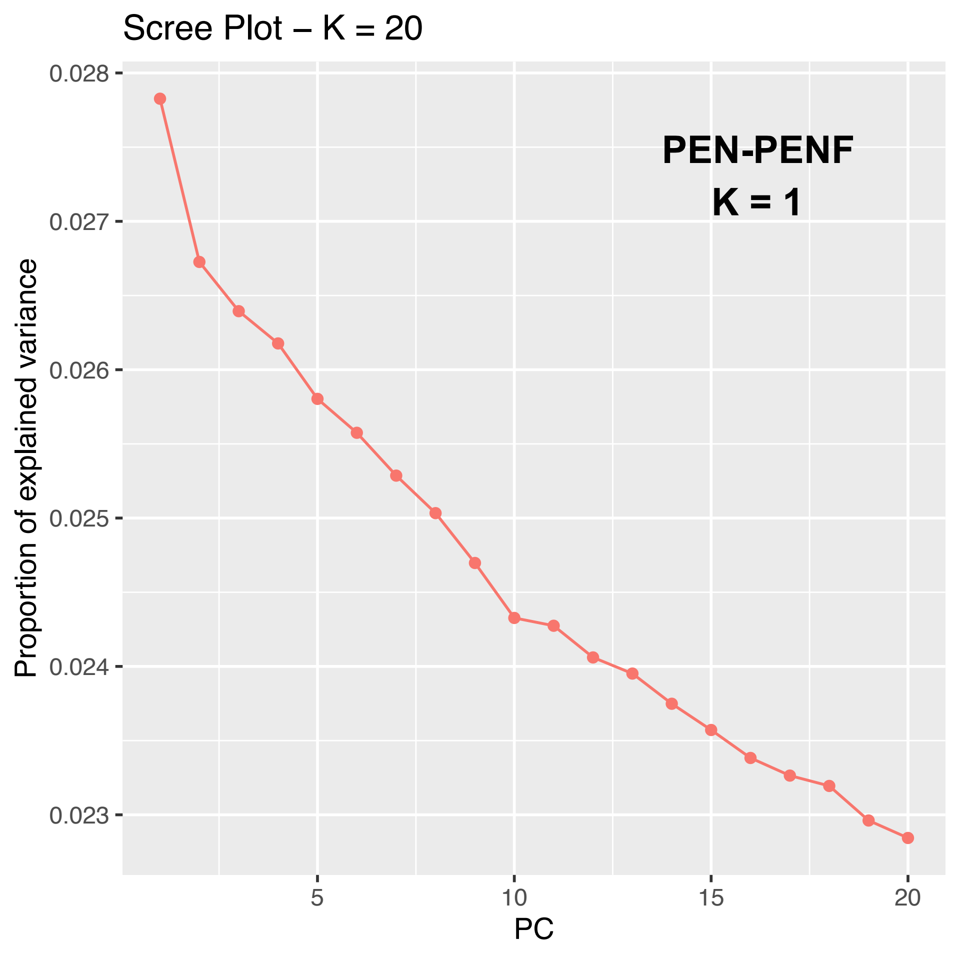

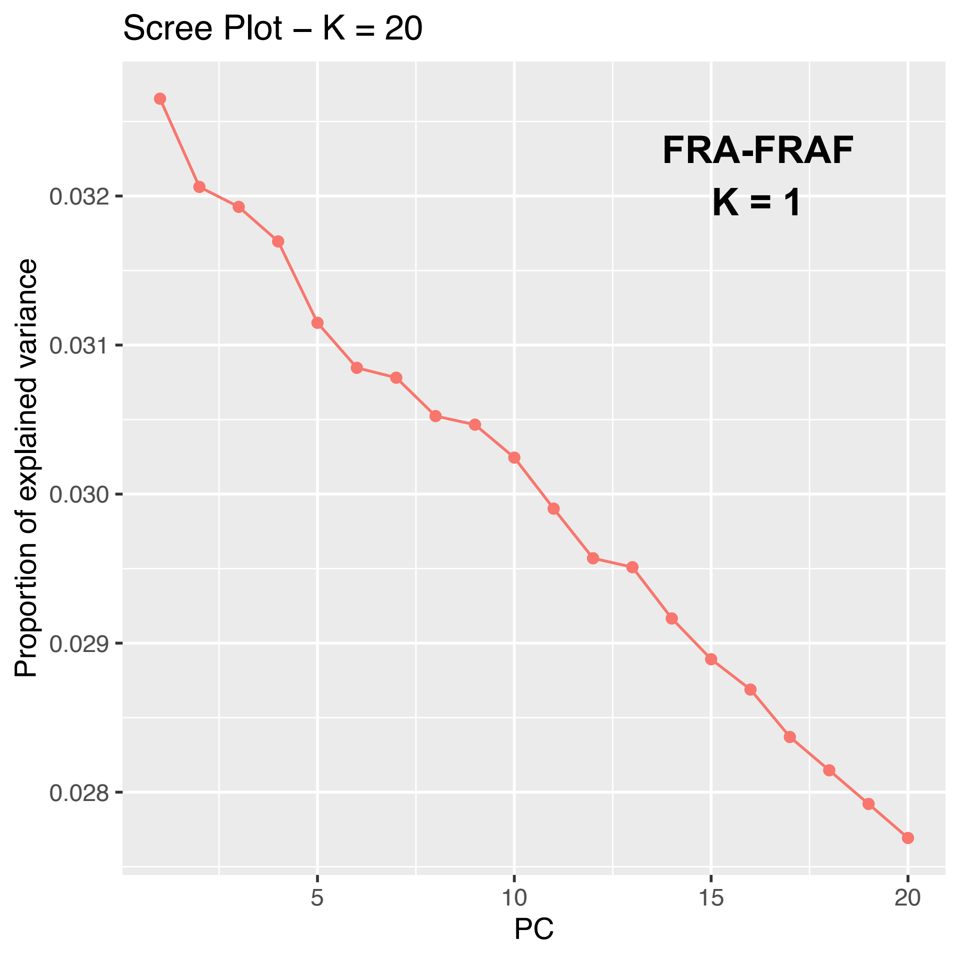

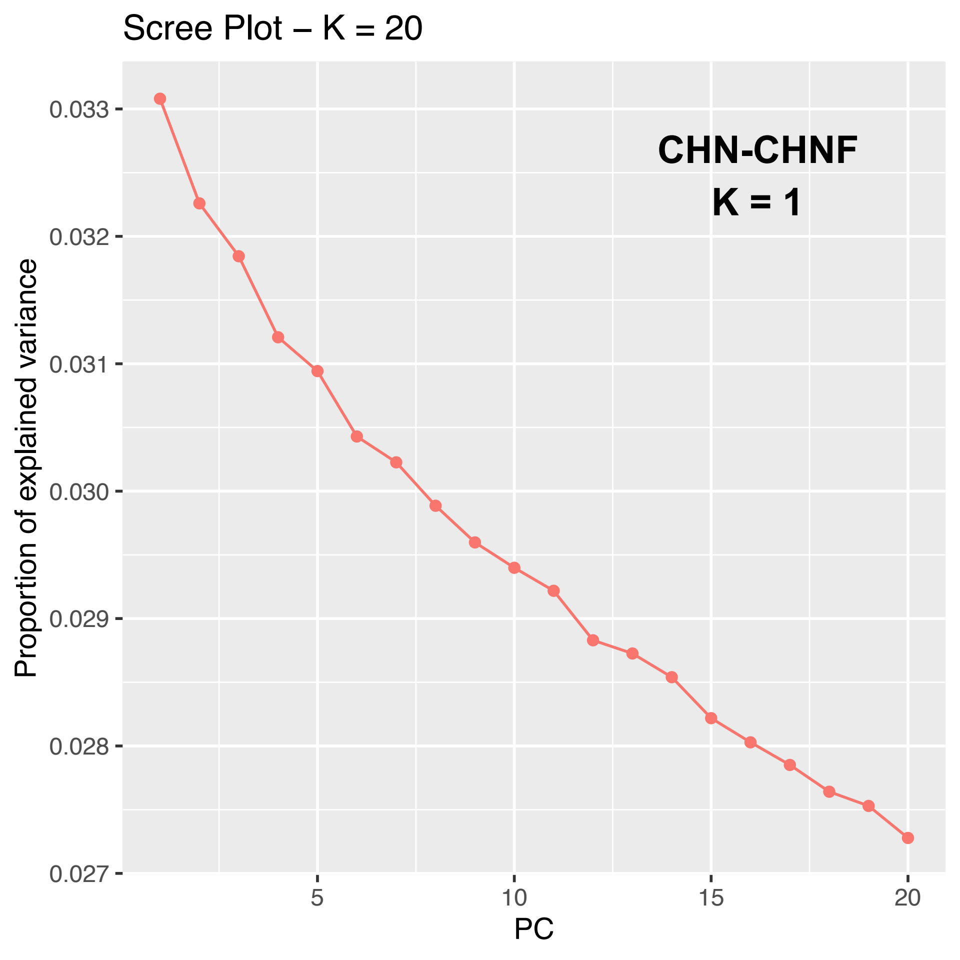
