## Additional File S2 for "Relative genomic impacts of translocation history, hatchery practices, and farm selection in Pacific oyster *Crassostrea gigas* throughout the Northern Hemisphere"

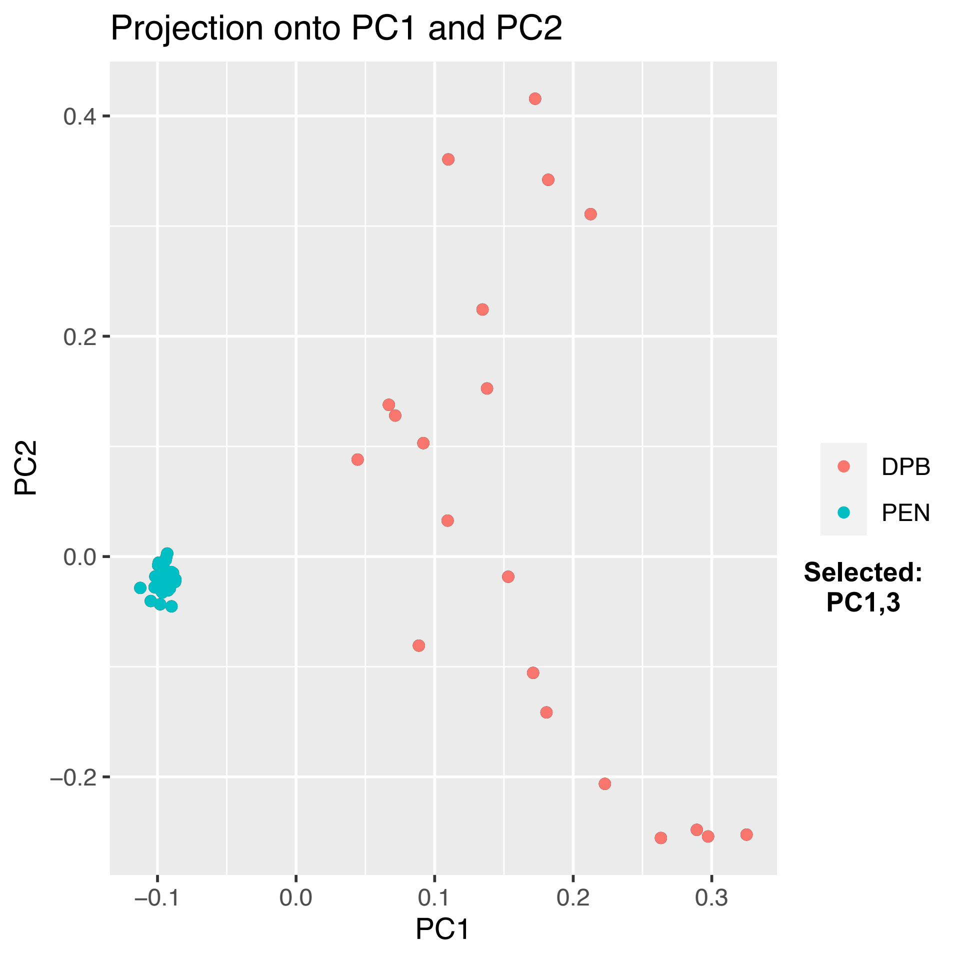

Projection onto PC1 and PC2
PC2
PC1
0.4
0.2
0.0
−0.2
−0.1
0.0
0.1
0.2
0.3
DPB
PEN
Selected:
PC1,3

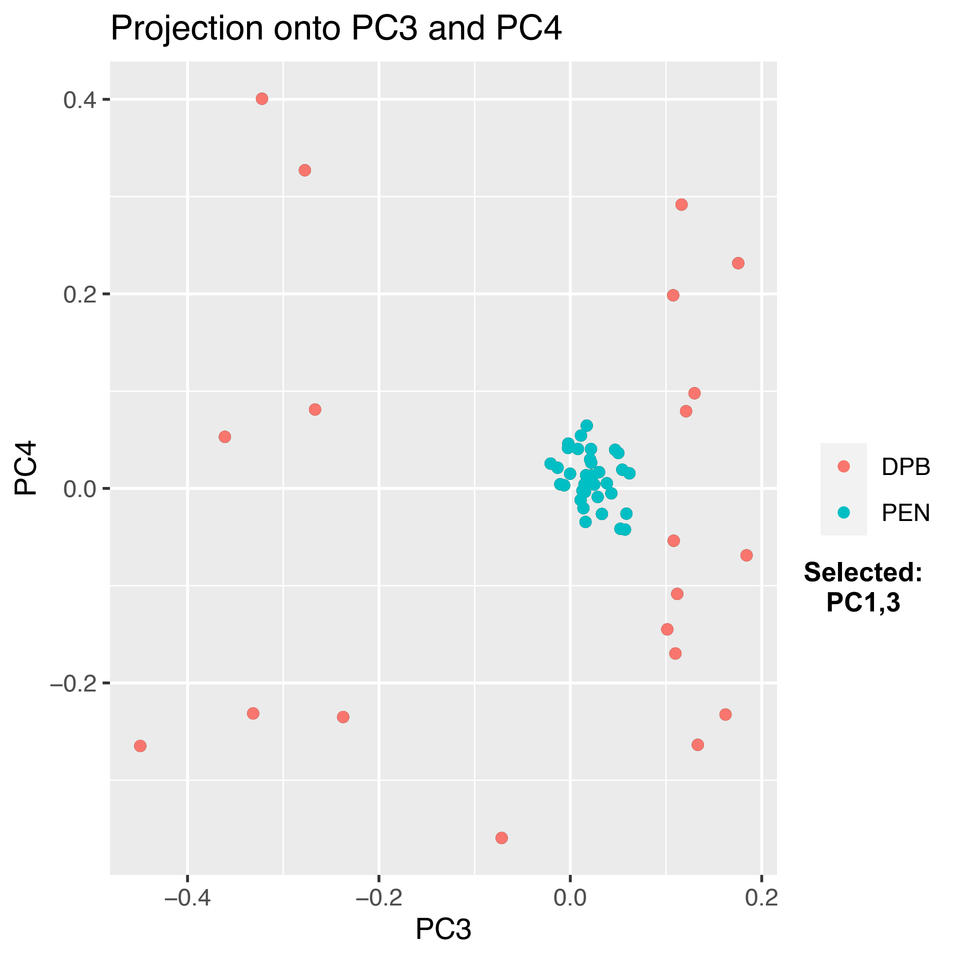
Projection onto PC3 and PC4
PC4
0.4
0.2
0.0
−0.2
PC3
−0.4
−0.2
0.0
0.2
DPB
PEN
Selected:
PC1,3

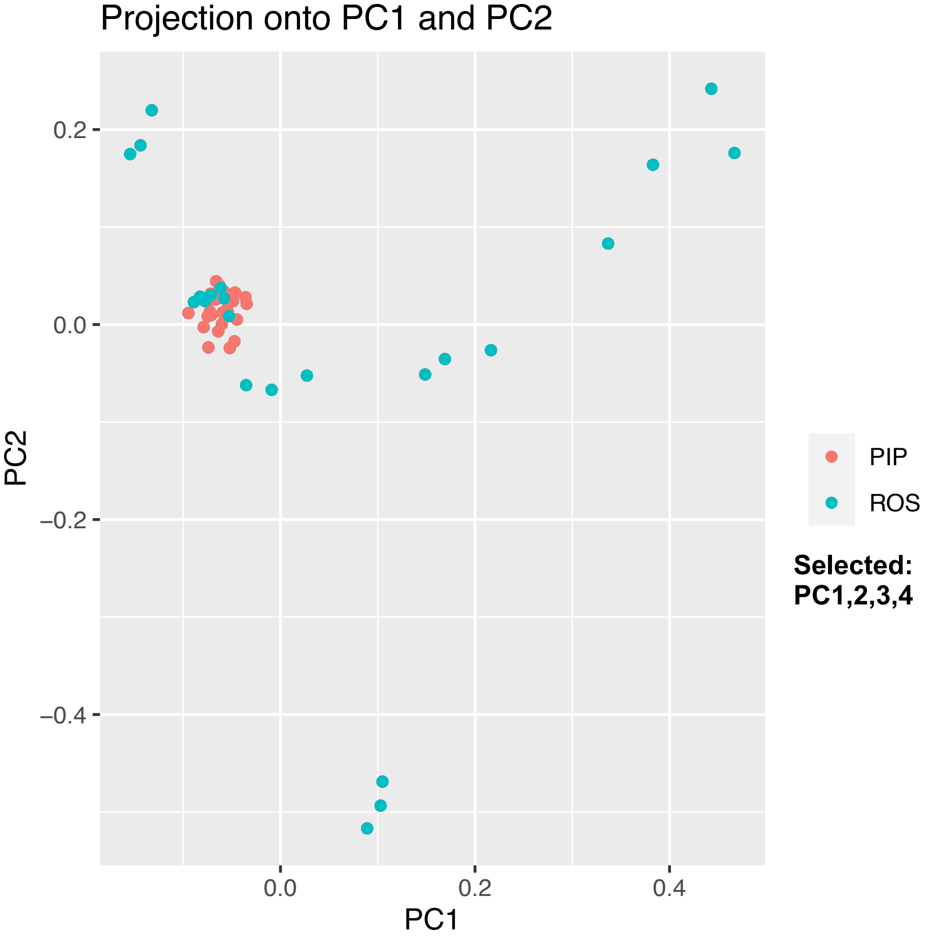
Projection onto PC1 and PC2
0.2
0.0
−0.2
−0.4
PC2
0.0
0.2
0.4
PC1
PIP
ROS
Selected:
PC1,2,3,4

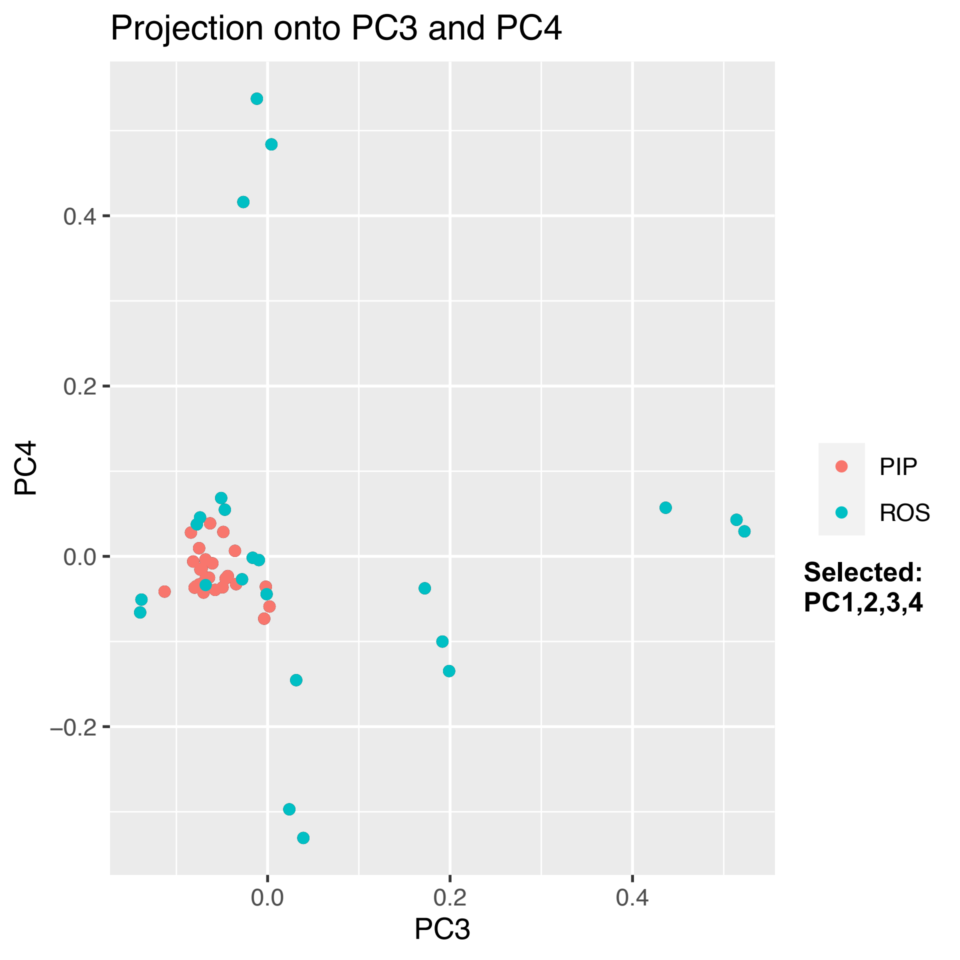
Projection onto PC3 and PC4
PC4
PC3
0.4
0.2
0.0
−0.2
0.0
0.2
0.4
PIP
ROS
Selected:
PC1,2,3,4

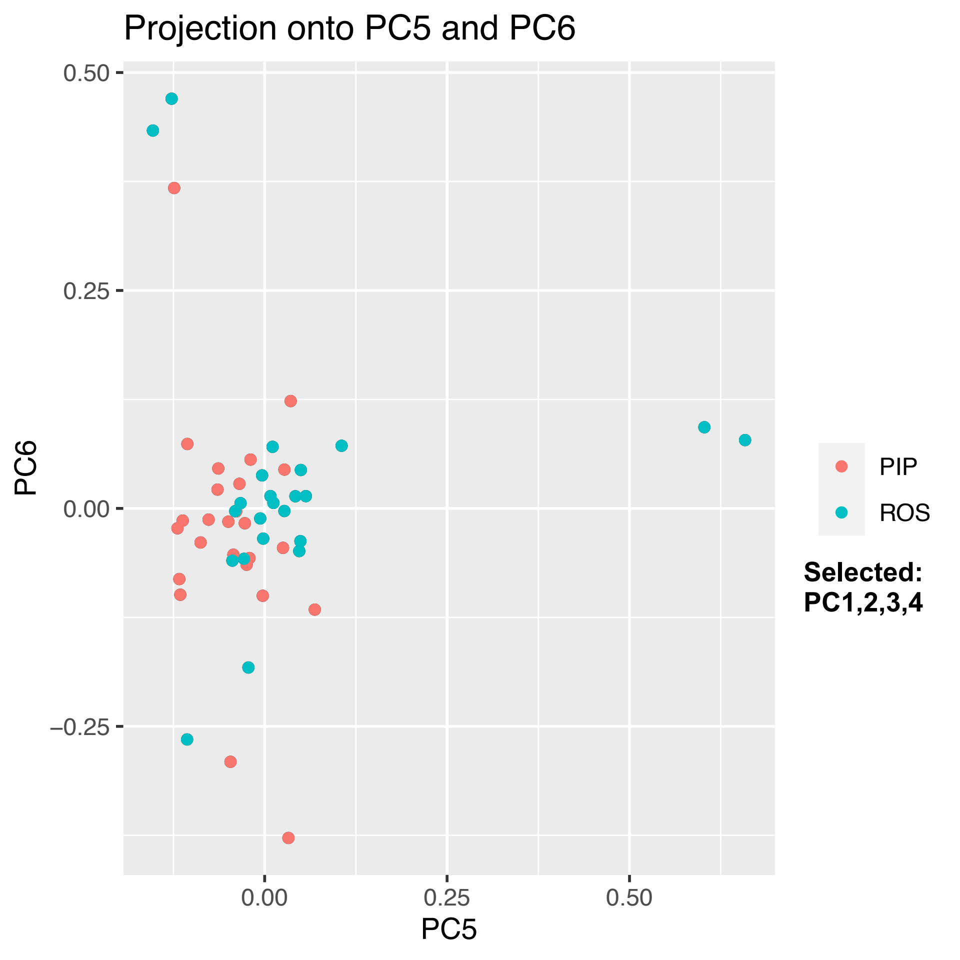
Projection onto PC5 and PC6
PC6
0.50
0.25
0.00
−0.25
0.00
0.25
0.50
PC5
PIP
ROS
Selected:
PC1,2,3,4

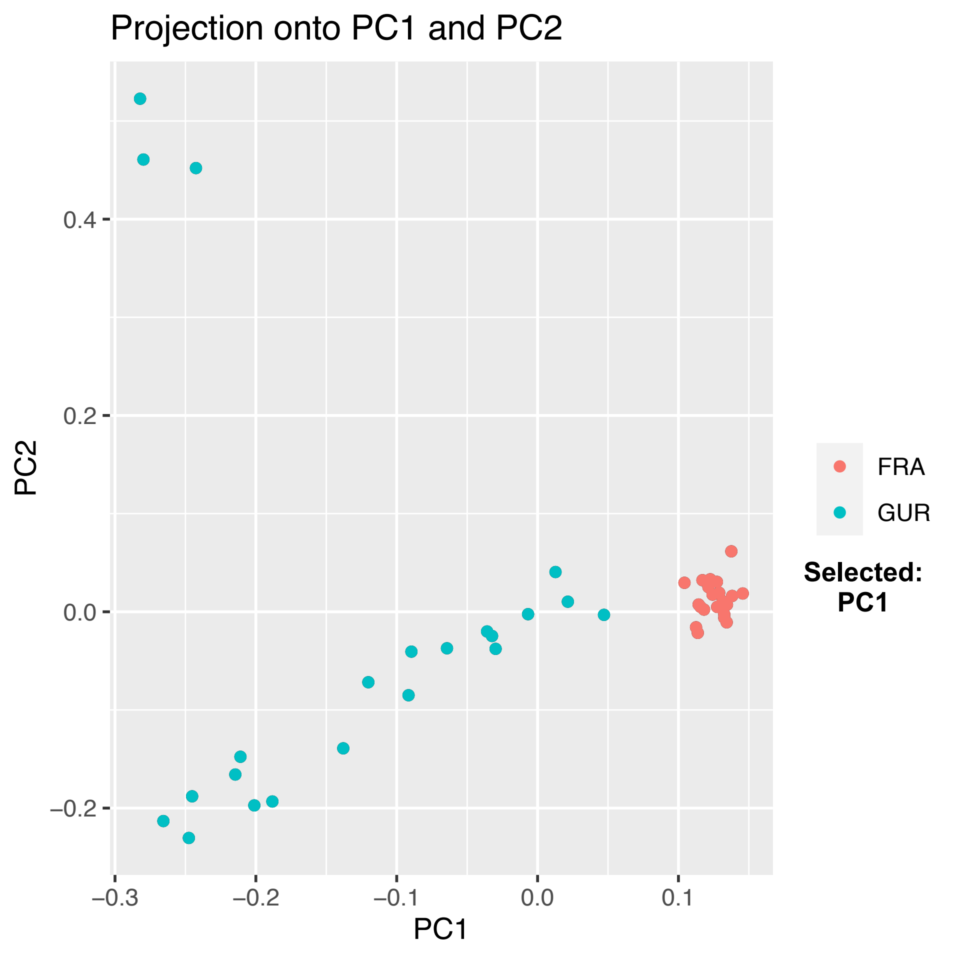
Projection onto PC1 and PC2
PC2
PC1
0.4
0.2
0.0
−0.2
−0.3
−0.2
−0.1
0.0
0.1
FRA
GUR
Selected:
PC1

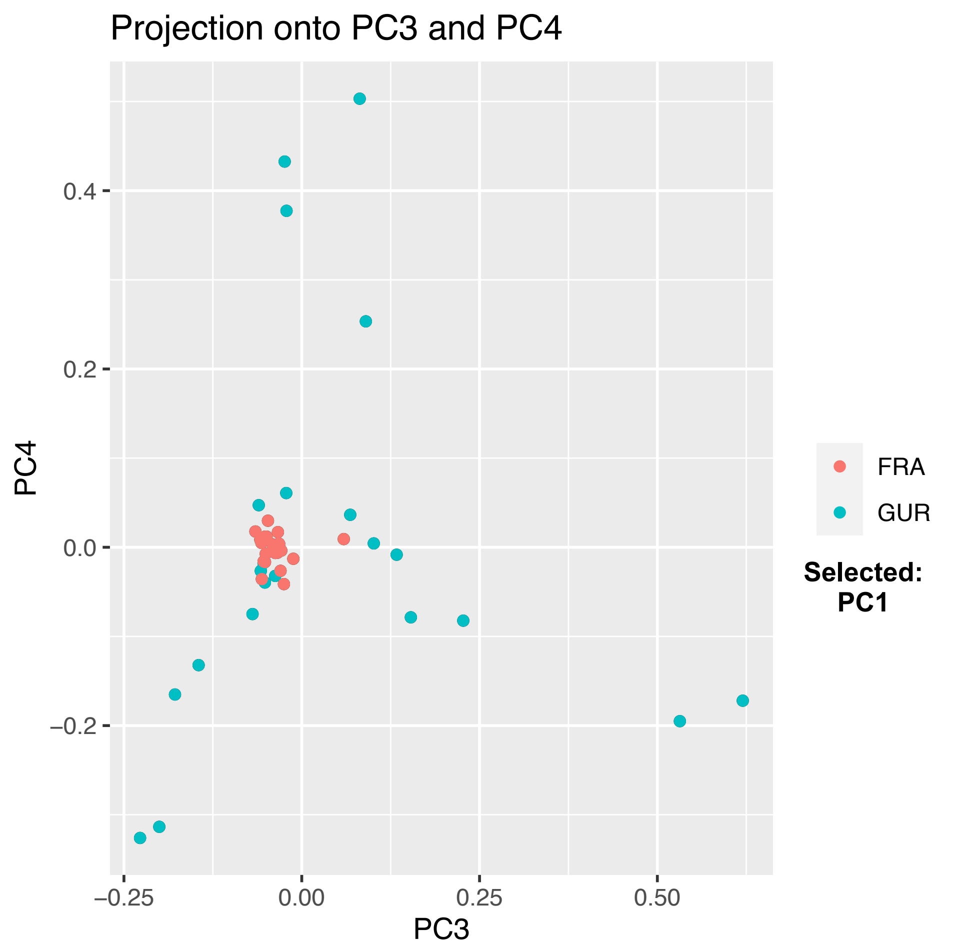
Projection onto PC3 and PC4
PC4
0.4
0.2
0.0
−0.2
−0.25
0.00
0.25
0.50
PC3
FRA
GUR
Selected:
PC1

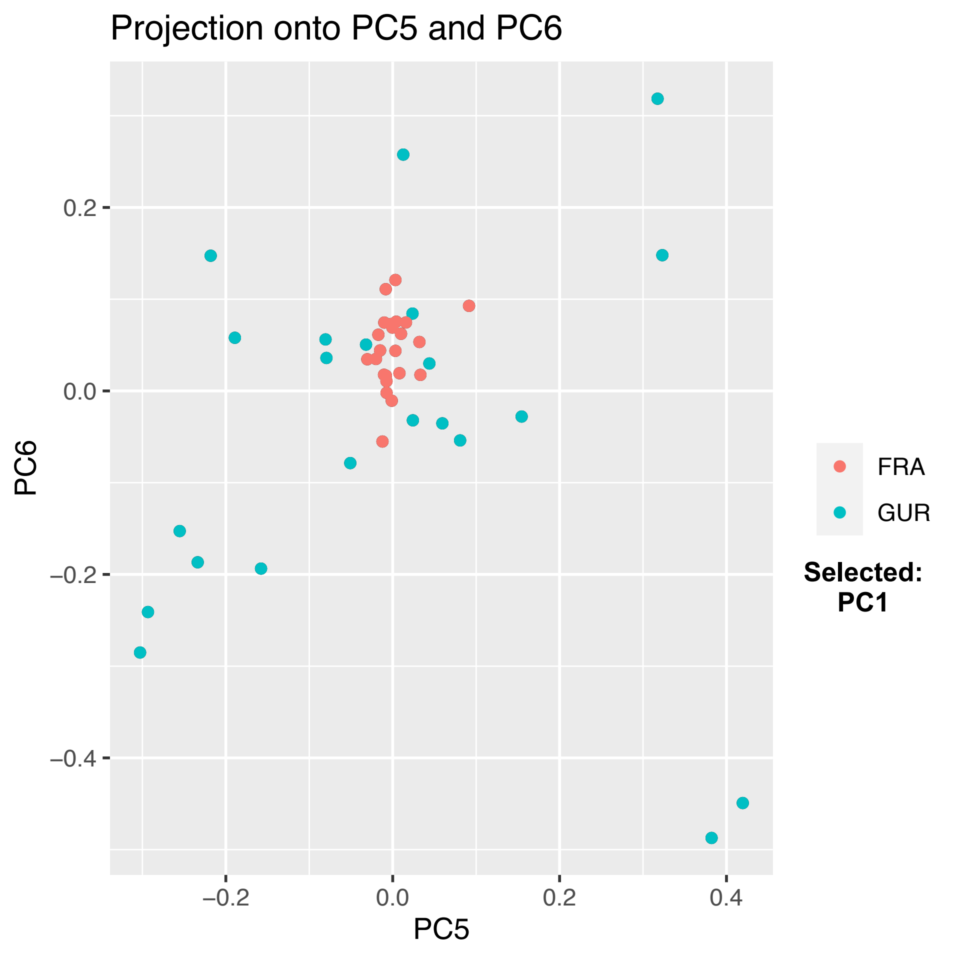
Projection onto PC5 and PC6
PC6
PC5
0.2
0.0
−0.2
−0.4
−0.2
0.0
0.2
0.4
FRA
GUR
Selected:
PC1

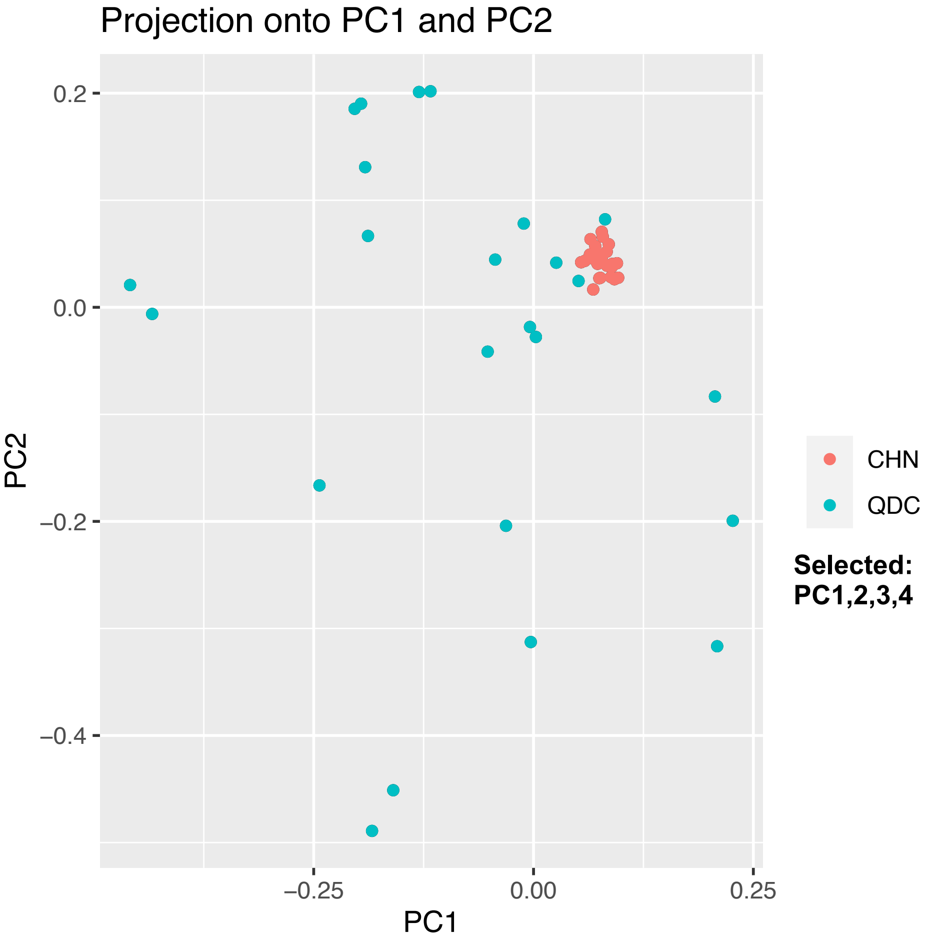

Projection onto PC1 and PC2
0.2
0.0
−0.2
−0.4
PC2
−0.25
0.00
0.25
PC1
CHN
QDC
Selected:
PC1,2,3,4

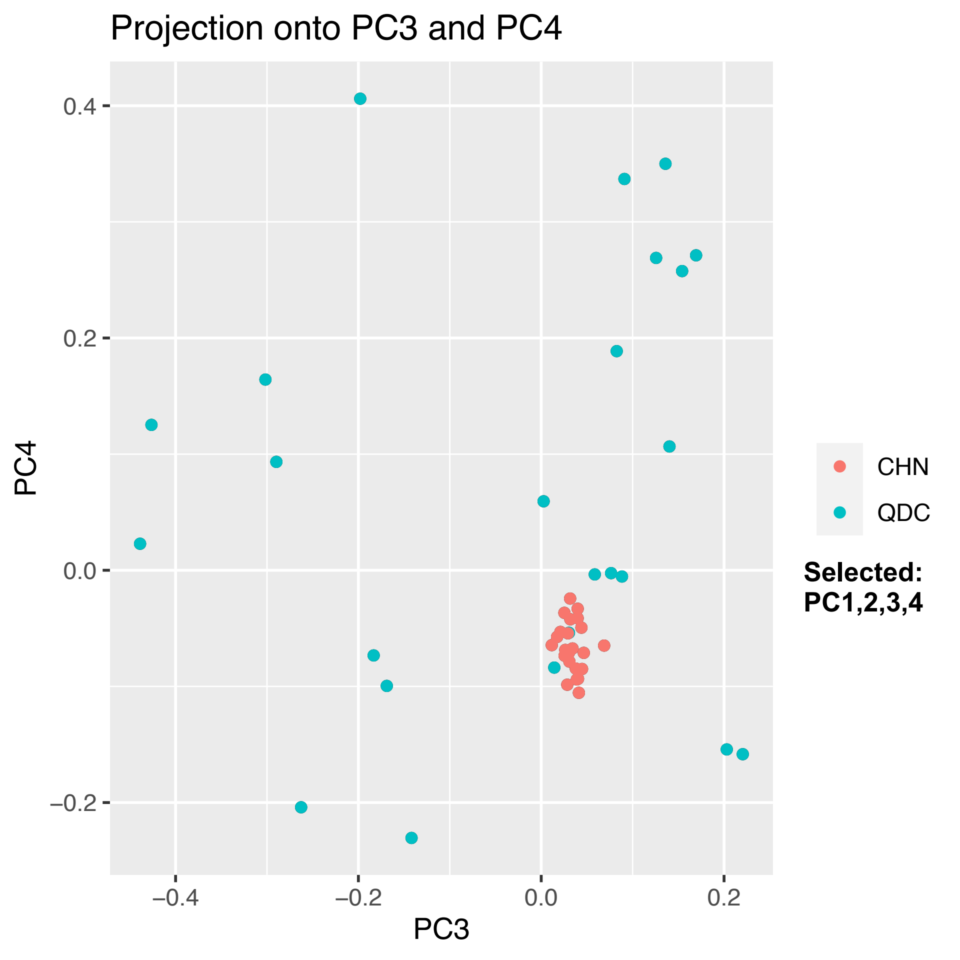
Projection onto PC3 and PC4
PC4
PC3
0.4
0.2
0.0
−0.2
−0.4
−0.2
0.0
0.2
CHN
QDC
Selected:
PC1,2,3,4

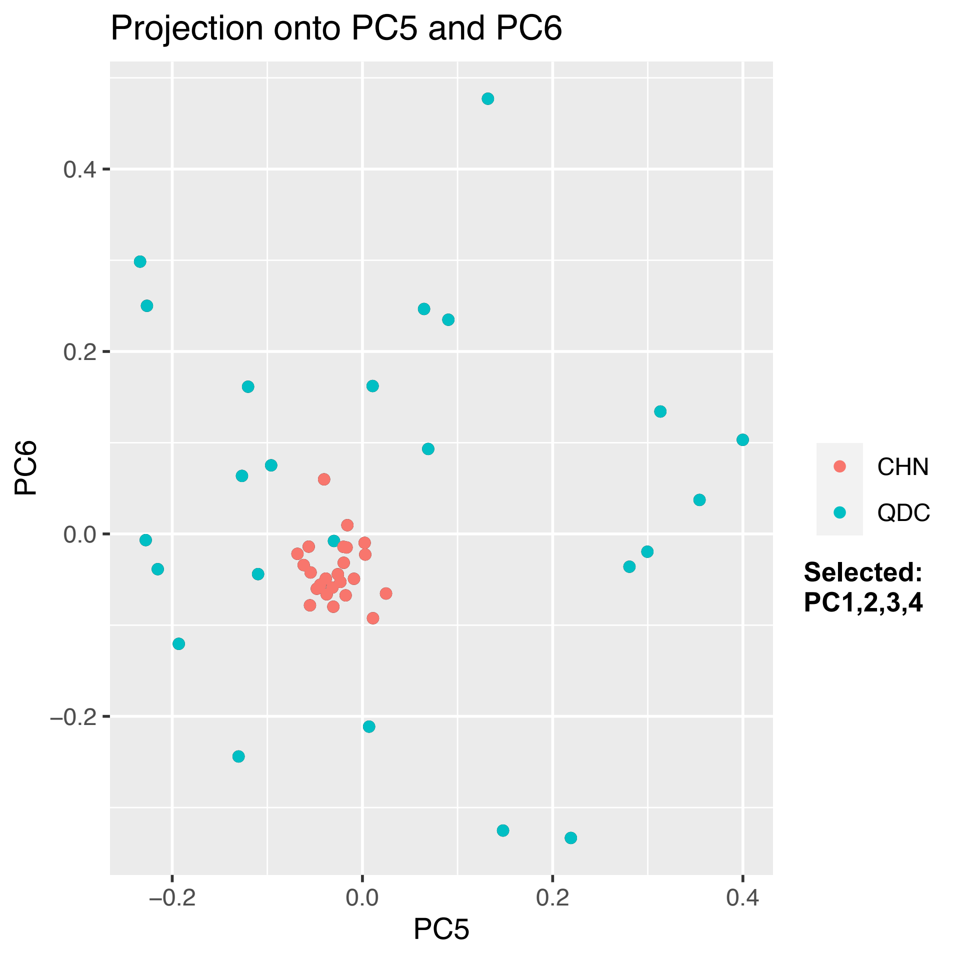
Projection onto PC5 and PC6
0.4
0.2
0.0
−0.2
PC6
−0.2
0.0
0.2
0.4
PC5
CHN
QDC
Selected:
PC1,2,3,4

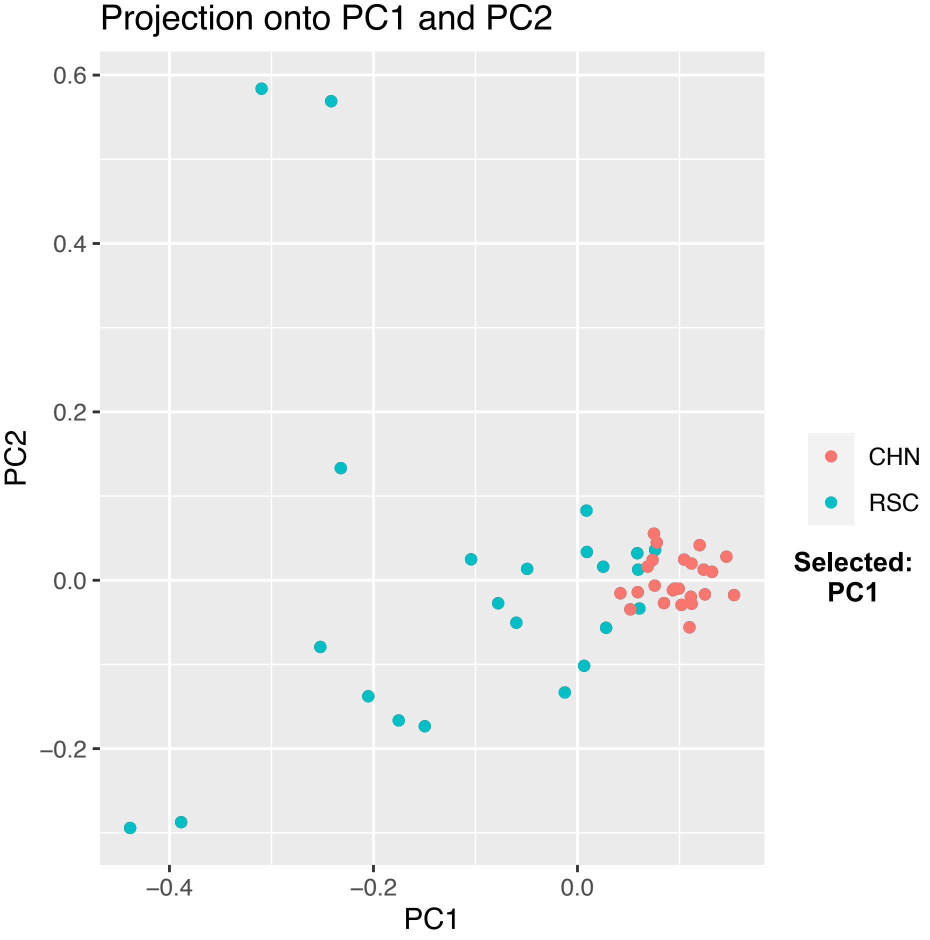

Projection onto PC1 and PC2
0.6
0.4
0.2
0.0
−0.2
PC2
−0.4
−0.2
0.0
PC1
CHN
RSC
Selected:
PC1

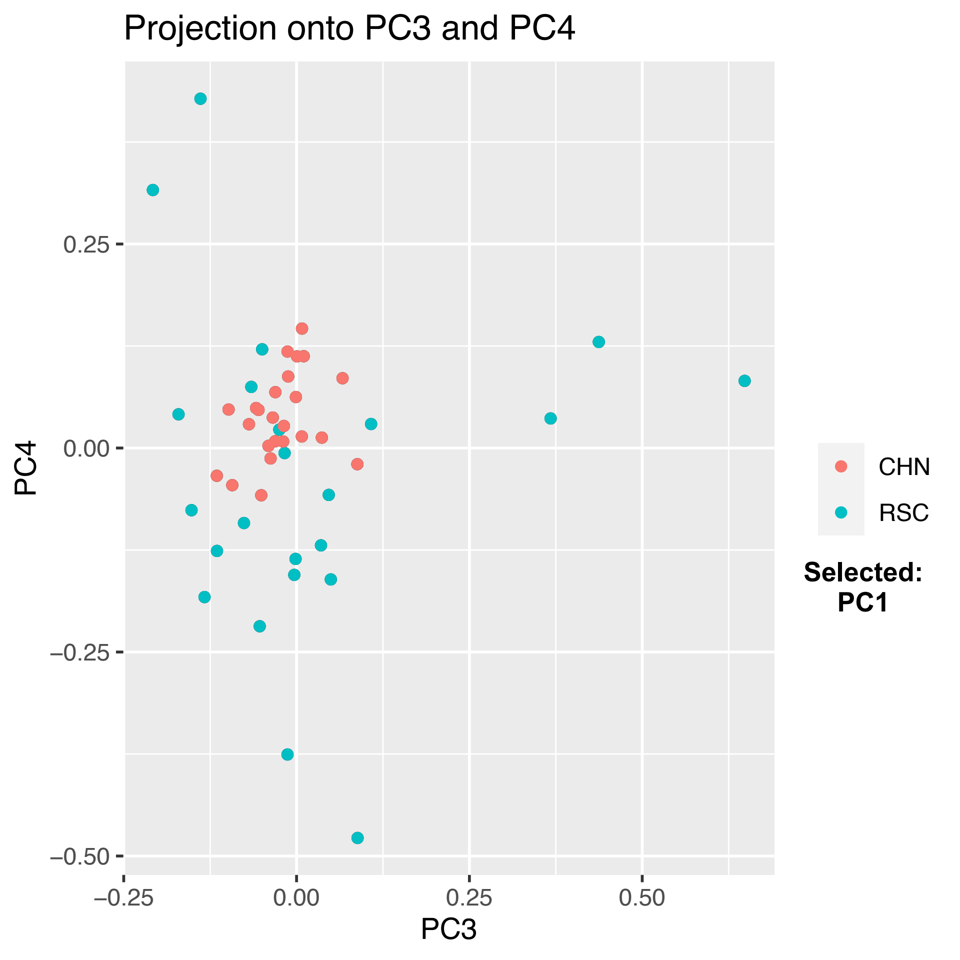

Projection onto PC3 and PC4
PC4
PC3
0.25
0.00
−0.25
−0.50
−0.25
0.00
0.25
0.50
CHN
RSC
Selected:
PC1

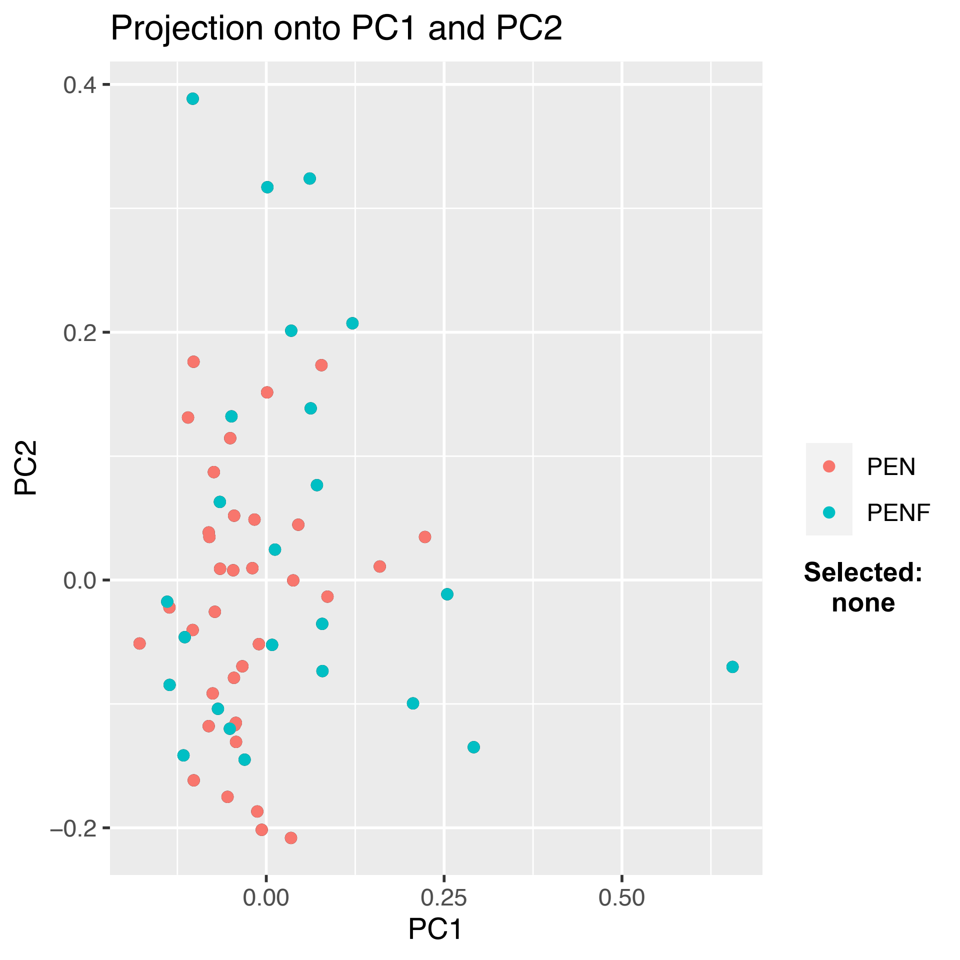

Projection onto PC1 and PC2
0.4
0.2
0.0
−0.2
PC2
0.00
0.25
0.50
PC1
PEN
PENF
Selected:
none

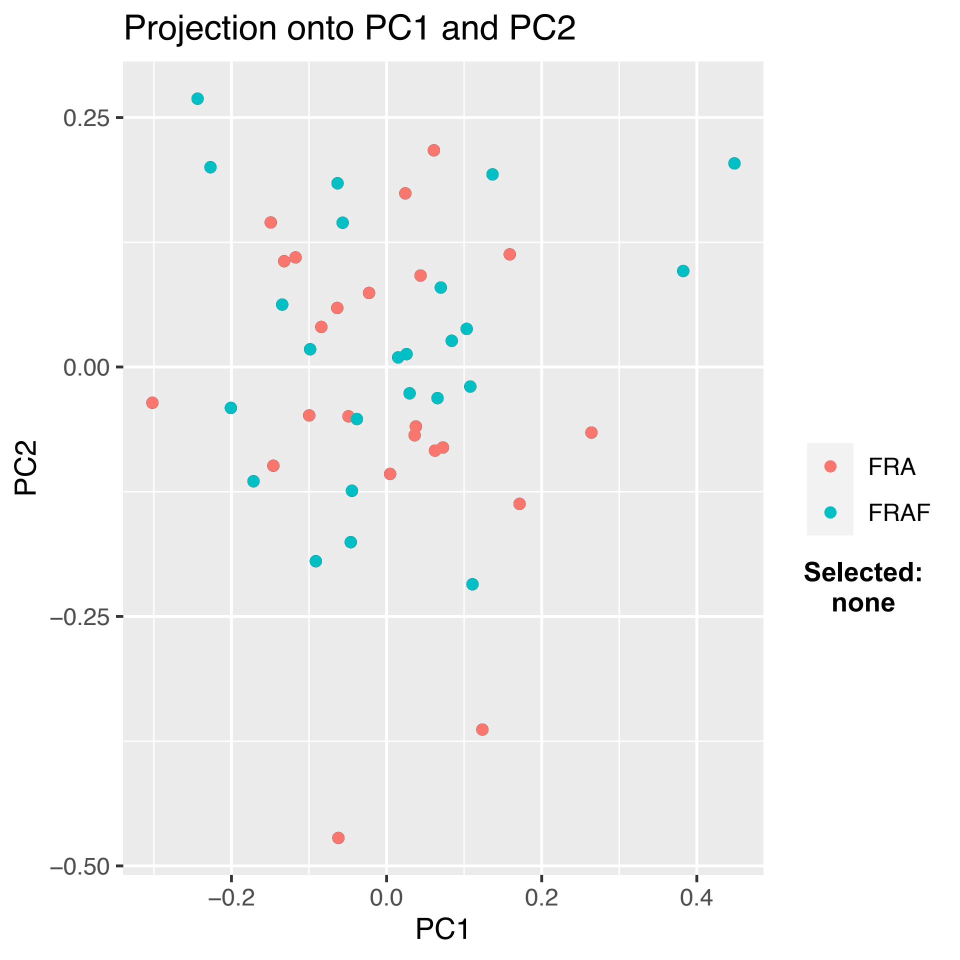

Projection onto PC1 and PC2
0.25
0.00
−0.25
−0.50
−0.2
0.0
0.2
0.4
PC1
PC2
FRA
FRAF
Selected:
none

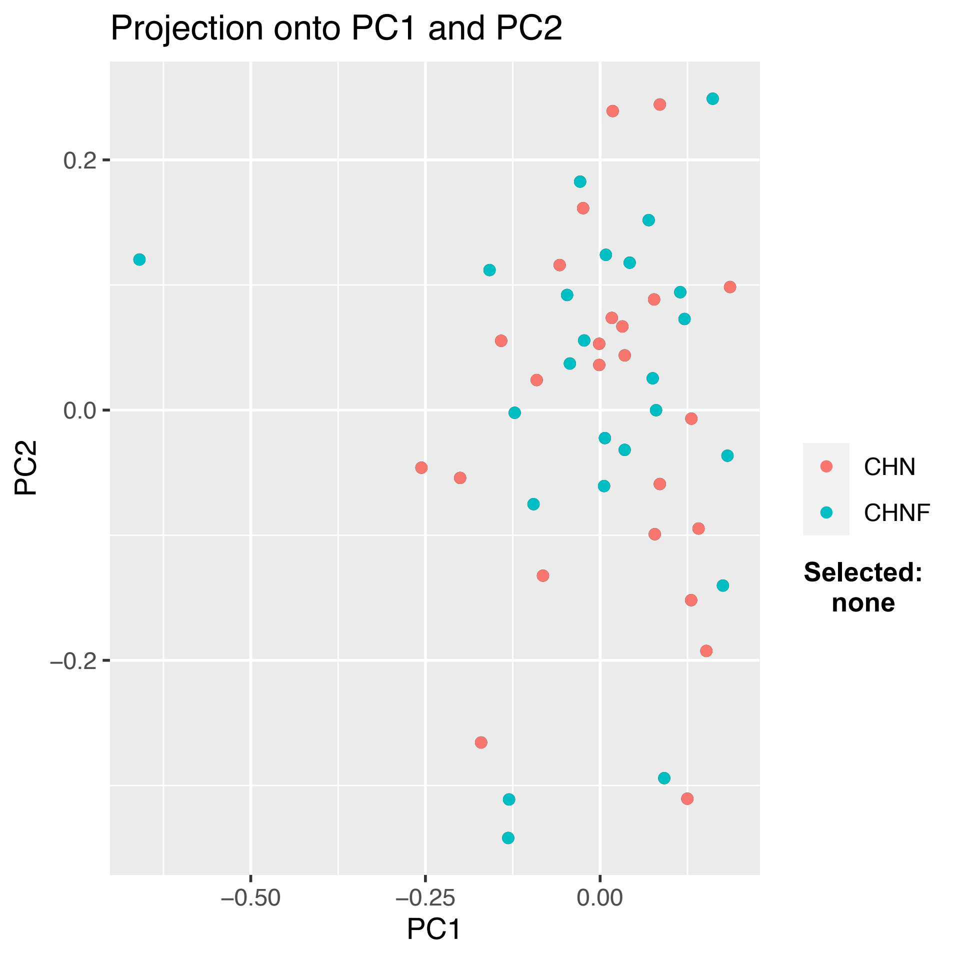

Projection onto PC1 and PC2
0.2
0.0
−0.2
PC2
−0.50
−0.25
0.00
PC1
CHN
CHNF
Selected:
none
