## Supplemental Results for "Relative genomic impacts of translocation history, hatchery practices, and farm selection in Pacific oyster *Crassostrea gigas* throughout the Northern Hemisphere"

### Supplemental Tables

**Table S1**. Mean genetic differentiation ($F_{ST}$) shown between pairs of populations for **(A)** populations in BC; and **(B)** populations in BC separated by size class; and **(C)** global populations. Full population names and locations are shown in Table 1 and Figures 1 and 2, respectively. Any $F_{ST}$ with a negative value was transformed to 0. Tables with these contrasts shown as 95% confidence intervals are presented in Table 4A-C.

**(A)**

| | HIS | PEN | PIP | SER |
|---|---|---|---|---|
| **HIS** | - | 0.0028 | 0.0022 | 0.0017 |
| **PEN** | | - | 0.0003 | 0.0025 |
| **PIP** | | | - | 0.0016 |
| **SER** | | | | - |

**(B)**

| | PEN_2 | PEN_3 | HIS_3 | HIS_6 |
|---|---|---|---|---|
| **PEN_2** | - | 0 | 0.0032 | 0.0030 |
| **PEN_3** | | - | 0.0036 | 0.0030 |
| **HIS_3** | | | - | 0.0015 |
| **HIS_6** | | | | - |

**(C)**

| | PEN | CHN | QDC | RSC | DPB | ROS | GUR | FRA | JPN |
|---|---|---|---|---|---|---|---|---|---|
| **PEN** | - | 0.0202 | 0.0396 | 0.0132 | 0.0604 | 0.0085 | 0.0288 | 0.0024 | 0.0060 |
| **CHN** | | - | 0.0196 | 0.0087 | 0.0795 | 0.0281 | 0.0459 | 0.0191 | 0.0241 |
| **QDC** | | | - | 0.0267 | 0.0964 | 0.0475 | 0.0635 | 0.0375 | 0.0416 |
| **RSC** | | | | - | 0.0722 | 0.0205 | 0.0383 | 0.0120 | 0.0165 |
| **DPB** | | | | | - | 0.0691 | 0.0728 | 0.0609 | 0.0635 |
| **ROS** | | | | | | - | 0.0352 | 0.0082 | 0.0129 |
| **GUR** | | | | | | | - | 0.0281 | 0.0313 |
| **FRA** | | | | | | | | - | 0.0048 |
| **JPN** | | | | | | | | | - |

**Table S2.** Population summary statistics from *Stacks* for (A) the single SNP per marker data only including variant sites and (B) from microhaplotype data including all variant and fixed RAD loci. The percentage polymorphic represents the proportion of nucleotides that are variant for the population within all RAD loci, including those RAD loci that have no polymorphisms present. The number of individuals is a fractional number due to missing data at some loci.

**(A)**

| Pop ID | Num. Indiv. | Obs. Het. | Std. Err. | Obs. Hom. | Std. Err. | Fis | StdErr | 95% C.I. |
|---|---|---|---|---|---|---|---|---|
| PIP | 23.1 | 0.07553 | 0.00084 | 0.92447 | 0.00084 | 0.072 | 0.016 | 0.041-0.103 |
| PEN | 32.5 | 0.07680 | 0.00082 | 0.92320 | 0.00082 | 0.086 | 0.021 | 0.045-0.127 |
| DPB | 18.9 | 0.08179 | 0.00107 | 0.91821 | 0.00107 | 0.031 | 0.012 | 0.008-0.053 |
| ROS | 21.5 | 0.07834 | 0.00089 | 0.92166 | 0.00089 | 0.064 | 0.013 | 0.038-0.09 |
| PENF | 22.7 | 0.07745 | 0.00084 | 0.92255 | 0.00084 | 0.074 | 0.012 | 0.05-0.098 |
| SER | 21.6 | 0.07735 | 0.00085 | 0.92265 | 0.00085 | 0.060 | 0.013 | 0.035-0.085 |
| HIS | 36.5 | 0.07720 | 0.00081 | 0.92280 | 0.00081 | 0.082 | 0.020 | 0.043-0.122 |
| GUR | 20.7 | 0.07918 | 0.00094 | 0.92082 | 0.00094 | 0.052 | 0.013 | 0.028-0.077 |
| RSC | 21.7 | 0.07732 | 0.00086 | 0.92268 | 0.00086 | 0.070 | 0.012 | 0.047-0.093 |
| QDC | 22.7 | 0.07847 | 0.00092 | 0.92153 | 0.00092 | 0.055 | 0.013 | 0.03-0.08 |
| CHN | 22.8 | 0.07531 | 0.00083 | 0.92469 | 0.00083 | 0.073 | 0.012 | 0.05-0.096 |
| CHNF | 22.6 | 0.07588 | 0.00085 | 0.92412 | 0.00085 | 0.073 | 0.013 | 0.048-0.098 |
| FRA | 21.8 | 0.07828 | 0.00084 | 0.92172 | 0.00084 | 0.079 | 0.011 | 0.057-0.101 |
| FRAF | 22.7 | 0.07866 | 0.00084 | 0.92134 | 0.00084 | 0.080 | 0.012 | 0.057-0.103 |
| JPN | 12.2 | 0.07931 | 0.00093 | 0.92069 | 0.00093 | 0.053 | 0.008 | 0.038-0.068 |

**(B)**

| Pop ID | Num. Indiv. | Polymorphic Sites | Percent Polymorphic Loci |
|---|---|---|---|
| **PIP** | 23.9 | 10951 | 0.28 |
| **PEN** | 33.6 | 12445 | 0.32 |
| **DPB** | 19.3 | 7244 | 0.19 |
| **ROS** | 22.1 | 10041 | 0.26 |
| **PENF** | 23.2 | 11056 | 0.29 |
| **SER** | 22.0 | 10702 | 0.28 |
| **HIS** | 37.4 | 12894 | 0.33 |
| **GUR** | 21.1 | 8623 | 0.22 |
| **RSC** | 22.0 | 10398 | 0.27 |
| **QDC** | 23.1 | 9174 | 0.24 |
| **CHN** | 23.1 | 10684 | 0.28 |
| **CHNF** | 23.1 | 10711 | 0.28 |
| **FRA** | 22.1 | 11134 | 0.29 |
| **FRAF** | 23.2 | 11291 | 0.29 |
| **JPN** | 12.5 | 8442 | 0.22 |

**Supplemental Figures**

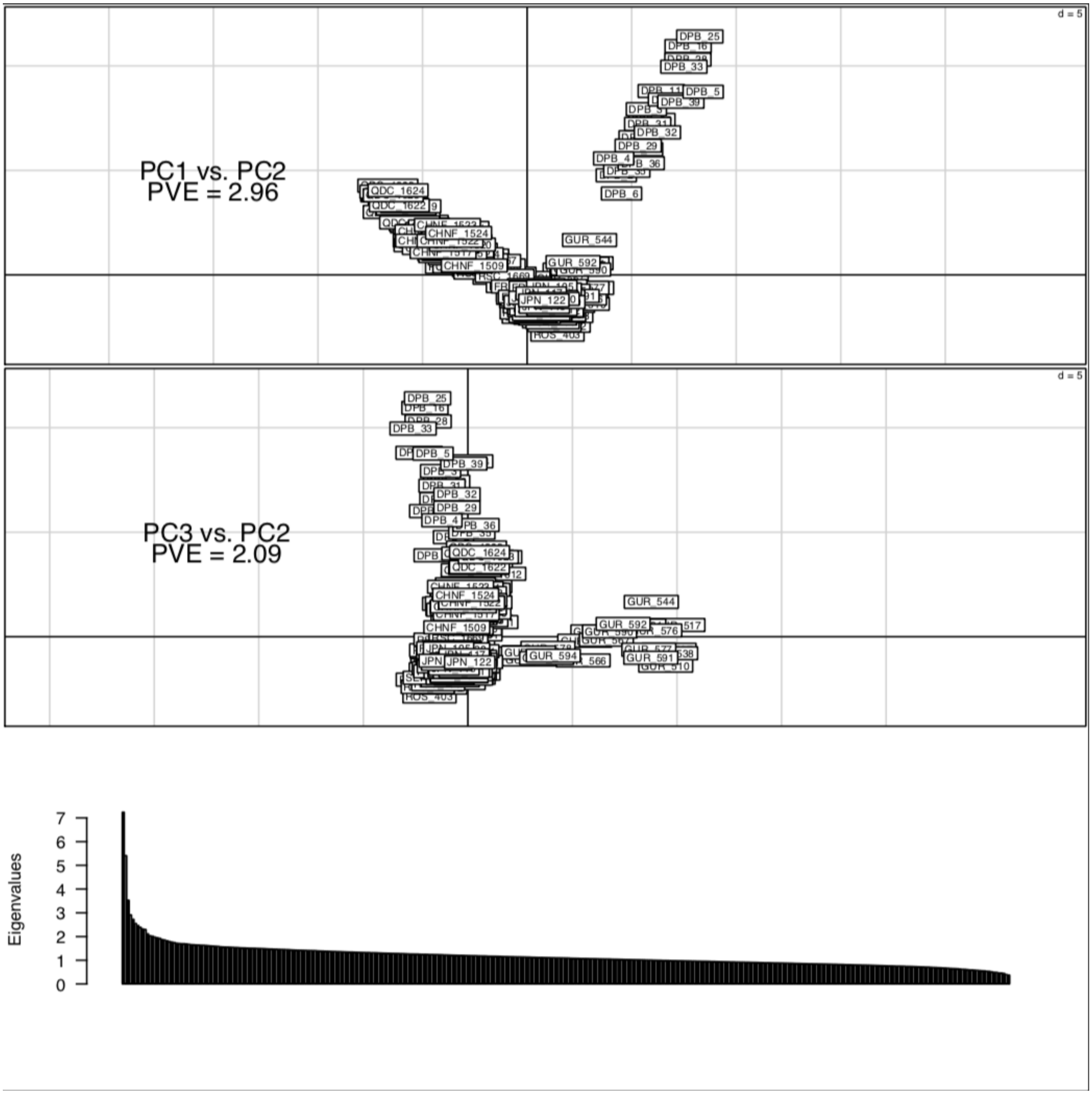


**Figure S1.** Principal Components Analysis (PCA) of all samples using a single SNP per locus with all loci. Sample IDs are plotted along PC 1 and 2 (A) or PC3 and 2 (B).

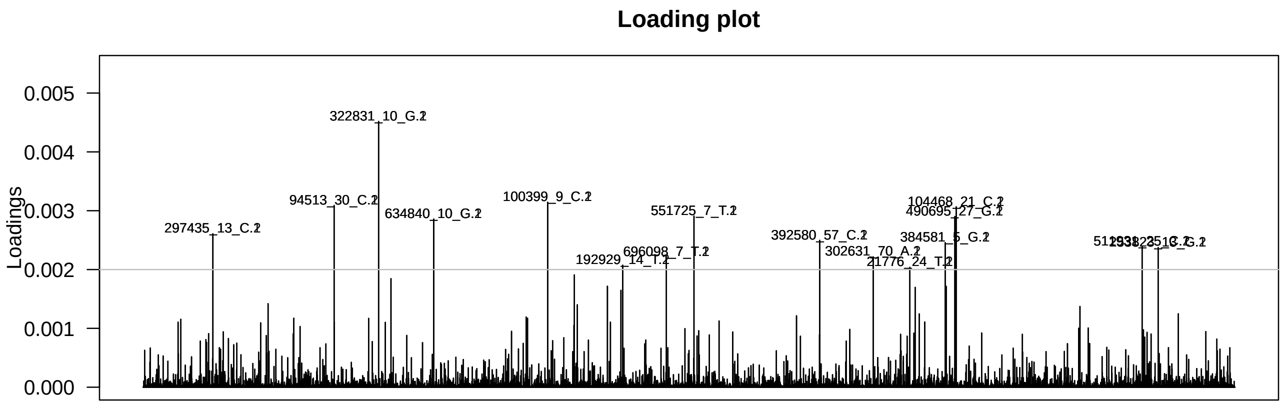


**Figure S2.** Discriminant Analysis of Principal Components (DAPC) marker loading values indicates that a small number of markers contributes the most to the DAPC differentiation. The 16 markers with loadings greater than 0.002 (grey line) are named.

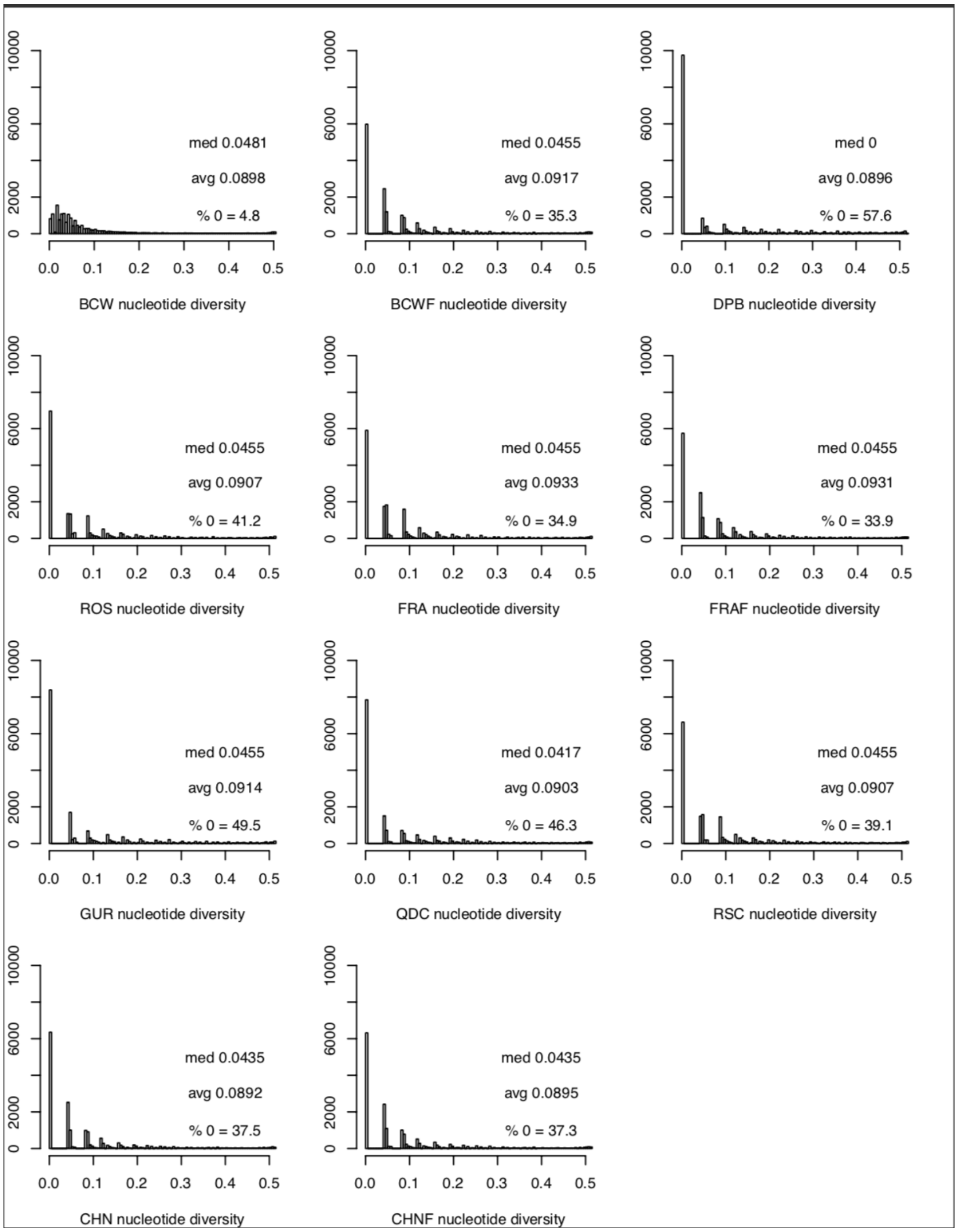


**Figure S3**. Per population distribution of nucleotide diversity (π), shown as frequencies of specific bins of nucleotide diversity as calculated by vcftools (--site-pi). Values are shown per population for the median, average, and percentage of markers with a zero value.

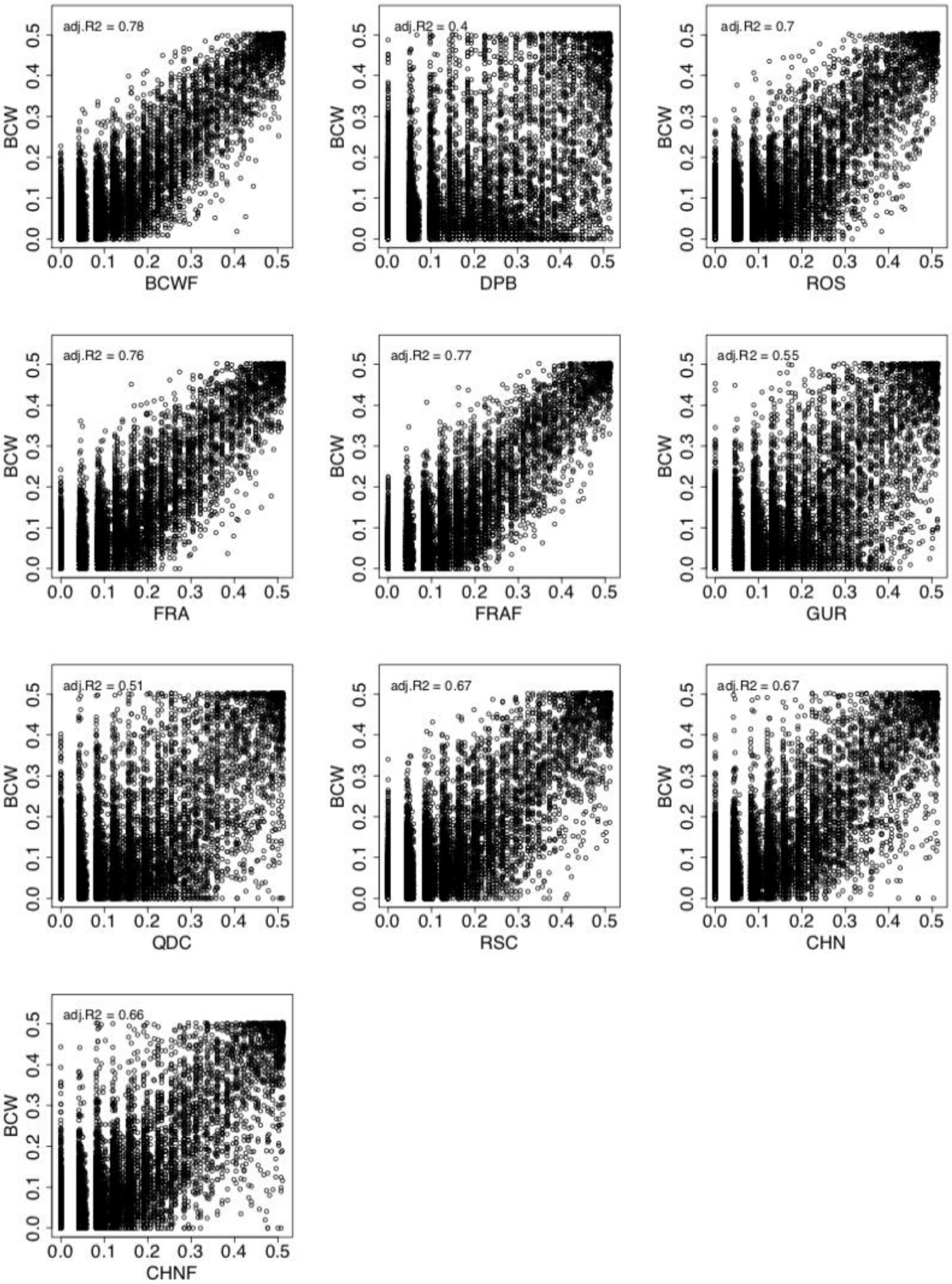


**Figure S4.** Nucleotide diversity (π) for each marker (dot) correlated between each population with the BC wild population. Correlated populations with BC wild samples, such as France (FRAF) show high correlation coefficients ($R^2 = 0.77$), whereas the hatchery-farmed populations that are more divergent have in general lower correlation, such as GUR ($R^2 = 0.55$), QDC (R2 = 0.51), and DPB (R2 = 0.4). Not all hatchery-farm populations have such low correlation, and are typically more similar to local wild populations, such as ROS and RSC ($R^2 = 0.7$ and 0.67, respectively).

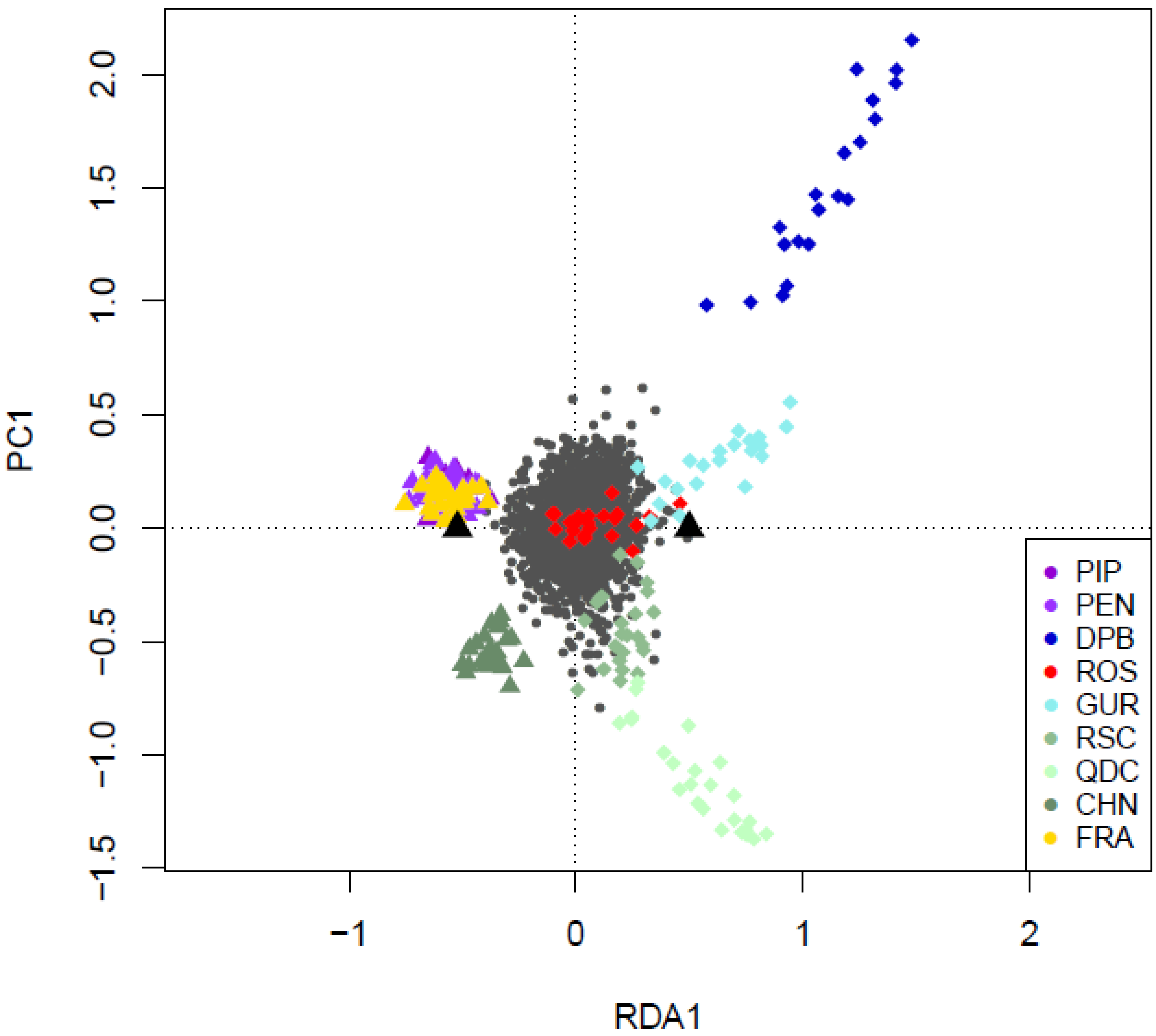


**Figure S5.** Comparison between Redundancy Analysis (RDA) and PCA in differentiating populations. RDA1 differentiated hatchery/hatchery-farmed samples (coloured triangles) from naturalized/wild samples (coloured diamonds). PC1 broadly separated the Chinese samples (negative PC1) from the BC samples, as well as separating out the DPB samples (positive PC1). The tight clustering of naturalized samples along PC1 and RDA1 contrasts that observed in the farmed samples, particularly for QDC, DPB, and GUR, which also display high genetic differentiation from other populations. The grey circles are individual loci. The black triangles indicate the average position along RDA1 of the two groupings, farms or wild.

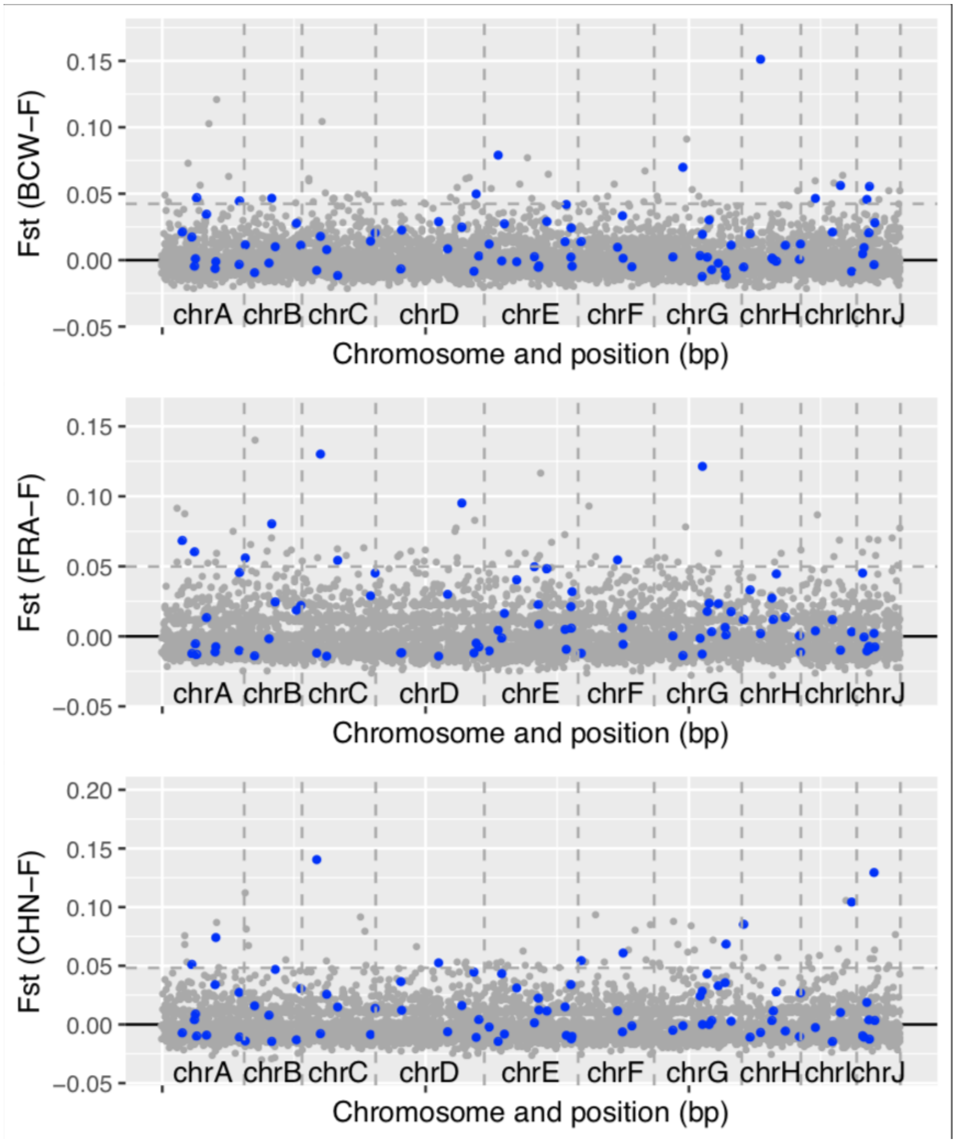


**Figure S6**. Outliers identified between naturalized/wild oysters moved onto a farm and naturalized populations (NAT-F vs. NAT) using RDA analysis (significant outliers shown by blue dots). The RDA was calculated using all NAT-F vs. NAT, whereas in plots, the $F_{ST}$ is calculated per contrast (see Table 2).

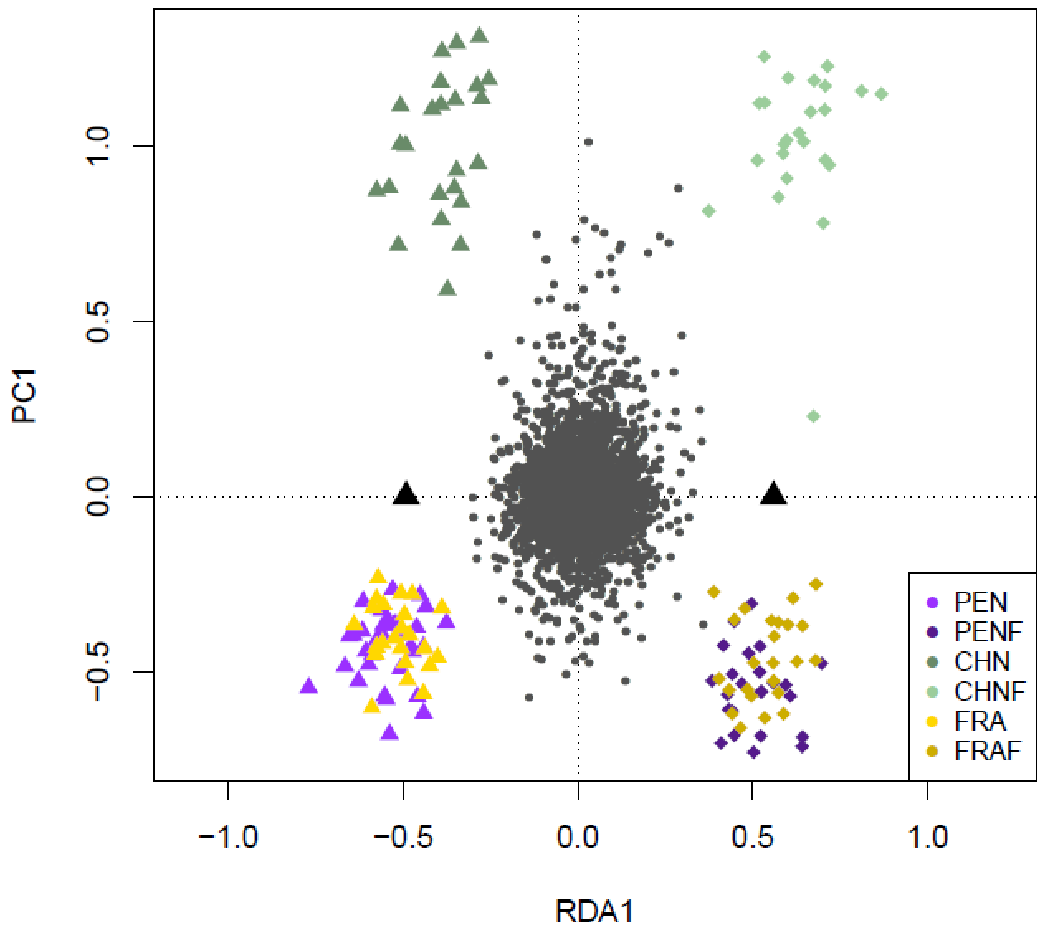


**Figure S7.** Comparison between Redundancy Analysis (RDA) and PCA in differentiating populations. RDA1 differentiated naturalized/wild oysters moved to farms (coloured diamonds) from naturalized/wild populations (coloured triangles), and PC1 differentiates the Chinese oysters from the oysters form France and Canada (PEN). The grey circles are individual loci. The black triangles indicate the average position along RDA1 of the two groupings, naturalized/wild oysters grown on farms or naturalized/wild oysters.

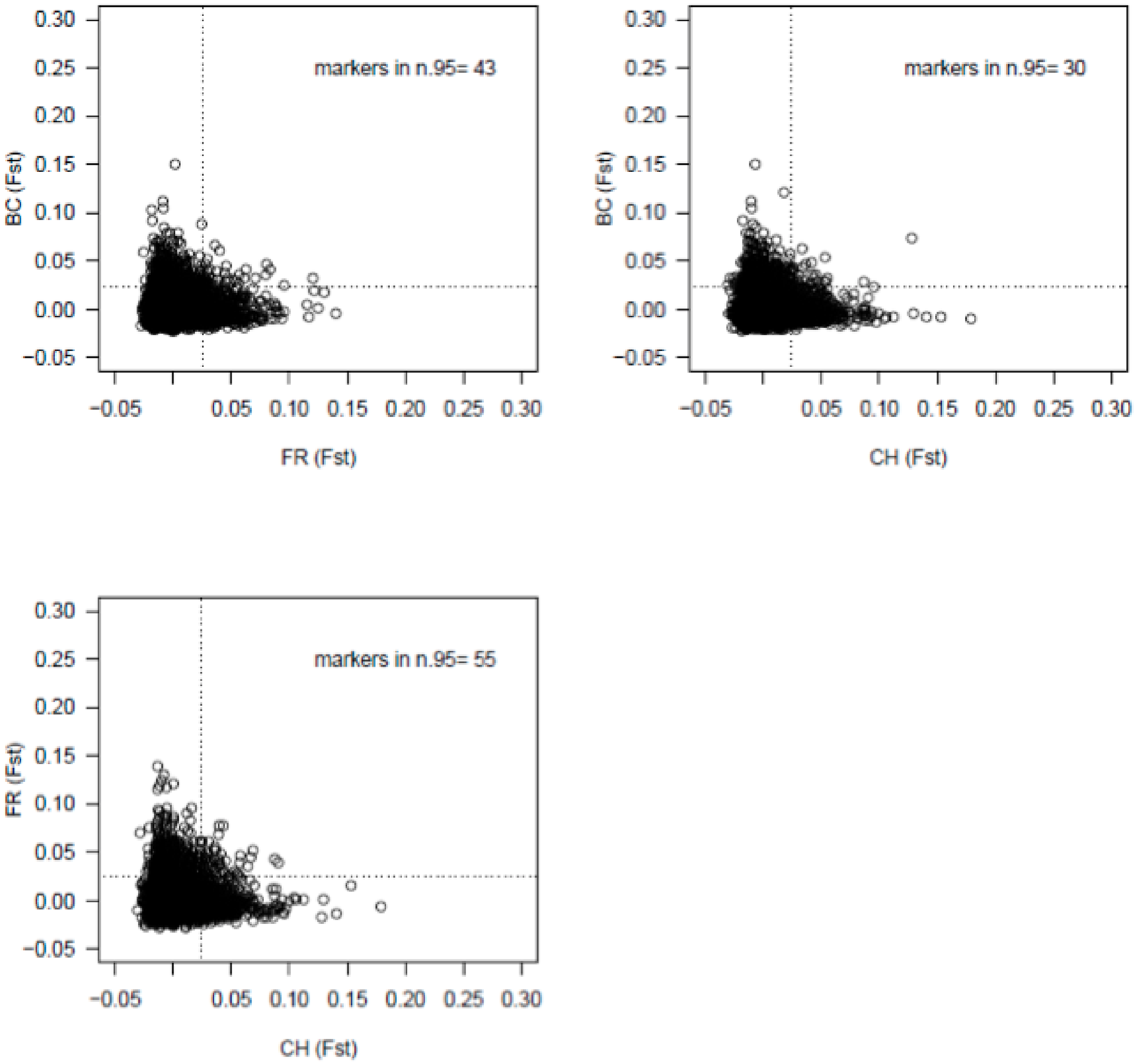


**Figure S8**. Per locus $F_{ST}$ values compared between different countries to detect signatures of parallel selection from being moved onto a farm. Genetic differentiation ($F_{ST}$) was calculated between oysters moved from naturalized populations into farms and compared per marker between sets of two contrasts each (e.g. $F_{ST}$ for FRA vs. FRAF compared to $F_{ST}$ for CHN vs. CHNF). The 95th percentile of $F_{ST}$ values are shown along each axis for each comparison, and the number of shared markers in both comparisons is shown on the graph (markers in n.95). Often high $F_{ST}$ markers were specific to one population, suggesting non-parallel selection from transplantation and growth on a farm.
